# Mining of natural diversity enables efficient and expressible peptide asparaginyl ligases

**DOI:** 10.64898/2025.12.16.694575

**Authors:** Wenyu Du, Shi Qi, Mingming Zhen, Miao Liao, Yang Zhao, Jiahui Ma, Yining Hao, Hui Jiang, Gengyu Lu, Chong Wai Liew, Niying Chua, Huachao Chen, Guangwan Hu, Xinya Hemu

## Abstract

Peptide asparaginyl ligases (PALs) are powerful tools for protein engineering but are limited by natural rarity and poor expression. We mined 23 cyclotide-rich *Viola* species, uncovering 29 new PALs that expanded the known repertoire to 47. A dual-objective screen identified VdiPAL1 as the best-performed natural PAL, with twice efficiency of *wt-*VyPAL2 and 12 mg L^-1^ soluble expression in *E. coli*. A broad P2’’ specificity including Trp/Ile/Leu/Phe/Tyr/Met was discovered across diverse PALs, which enables sequential click-compatible liposome dual-functionalization. 1.8 Å crystal structure of VdiPAL1 reveals a pre-organized near-attack conformation (NAC), supported by constant-pH MD simulations linking pH-dependent reactivity to NAC geometry. Our homology- and structure-based design yielded VyOpt1, a quintuple mutant of VyPAL2 with over 24-fold improved expression via enhanced cap-domain foldability in a single design-test cycle. This work expands the PAL family and demonstrates a transferable cap-domain-based engineering strategy, highlighting natural diversity as a powerful driver of enzyme discovery and optimization.

**TOC summary:** Mining of cyclotide-rich *Viola* genus expanded the total number of natural peptide asparaginyl ligases (PALs) to 47, including the discovery of highly expressible VdiPAL1. Its high-resolution structure provided new insights for mechanism, and its sequence guided us to a generalized PAL engineering strategy, leading to 24-fold increment in VyPAL2 expression.

## Introduction

Peptide ligases are pivotal biocatalysts that form backbone amide bonds, providing versatile tools for peptide and protein synthesis and modification^1^. They are widely valued for catalyzing the head-to-tail cyclization to improve peptide stability^2^, site-specific functionalization^3^, and orthogonal ligation strategies such as one-pot convergent synthesis and chemoenzymatic cascades^4^. These capabilities support a broad range of applications, including accessing conformationally rigid peptides and live-cell surface engineering^5, 6, 7^.

Among known peptide ligases, peptide asparaginyl ligases (PALs) have emerged as particularly versatile tools combining broad substrate tolerance, high catalytic efficiency, minimal hydrolysis, and scarless product formation, operating under mild and scalable aqueous conditions ^8,9^. These features make PALs especially attractive for scalable applications in synthetic biology and protein engineering, and complementary to other well-established systems such as intein^10,11^, sortase^12,13^, and subtiligase^14,15^, which have each found specific niches despite certain constraints like fusion-expression dependency or stringent sequence preferences^16^.

Despite their versatility, PALs are exceptionally rare in nature. Their discovery has largely relied on the elucidation of biosynthetic pathways for natural cyclic peptides and protein splicing products. In 2007, Saksa et al. proposed that asparaginyl endopeptidase (AEP) is responsible for post-translation backbone cyclization of cyclotide precursors, however the specific enzyme was not isolated or identified due to low abundancy in the chosen cyclotide-producing plants^17^. In 2014, butelase-1 was isolated from *Clitoria ternatea* as the first identified post-translational cyclase of cyclotides^18^. It was demonstrated that butelase-1, despite sharing high homology with asparaginyl endopeptidases (AEPs) and being structurally categorized under the C13 legumain family, functions as a ligase rather than a hydrolase. To clarify this distinction, the term peptide asparaginyl ligase (PAL) was introduced in 2019, following the identification of two Ligase Activity Determinants (LADs) in VyPAL2 from *Viola yedoensis* that control catalytic directionality^19^. The LAD concept not only enables rational conversion of AEPs into PAL-like enzymes via site-directed mutagenesis^20^, but also provides a molecular criterion for distinguishing PALs from AEPs^21^. Ligases discovered before that, including OaAEP1b from *Oldenlandia affinis* and HeAEP3 from *Hybanthus ennesaspermus*, have retained their earlier AEP nomenclature in literature.

LAD hypothesis has transformed PAL discovery from laborious protein isolation to efficient database mining, enabling large-scale in silico screening. Nevertheless, only 18 PALs (≈1%) were identified in a dataset of 1,500 legumain sequences, confirming their natural rarity^21^. Notably, all characterized PALs to date derive from cyclic peptide-producing plants, with more than half from the *Viola* genus. Given their cyclotide-rich biochemistry^22^, species diversity, and wide distribution, *Viola* represents the most promising natural reservoir for discovering new PALs^23, 24^.

To unlock the potential from natural diversity of PALs, we conducted a systematic investigation into *Viola* genus to expand the pool of PALs and deepen the mechanistic understanding of their catalytic efficiency. Through a pipeline combining sequence mining with high-throughput activity screening, we acquired novel legumain sequences, which not only enriched the natural PAL library but also led to the identification of VdiPAL1, a highly expressible and catalytically efficient ligase with extended substrate scope (**Fig. 1A**). Demonstrating its broad applicability, VdiPAL1 was successfully employed for the dual-functionalization of liposomes via a sequential click-compatible fluorescent targeting protocol (**Fig. 1B**). Furthermore, to elucidate the structural basis for catalysis, we determined the high-resolution crystal structure of VdiPAL1, leading us to propose a near-attack conformation (NAC) hypothesis as a potential spatial determinant of efficiency (**Fig. 1C**). Finally, leveraging evolutionary proximity analysis and computationally guided engineering, we developed a broadly applicable strategy to overcome heterologous expression challenges. This approach, featuring combinatorial mutagenesis within the cap domain, boosted the *E. coli* expression of the challenging ligase VyPAL2 by 24-fold (**Fig. 1D**)

**Fig. 1.**
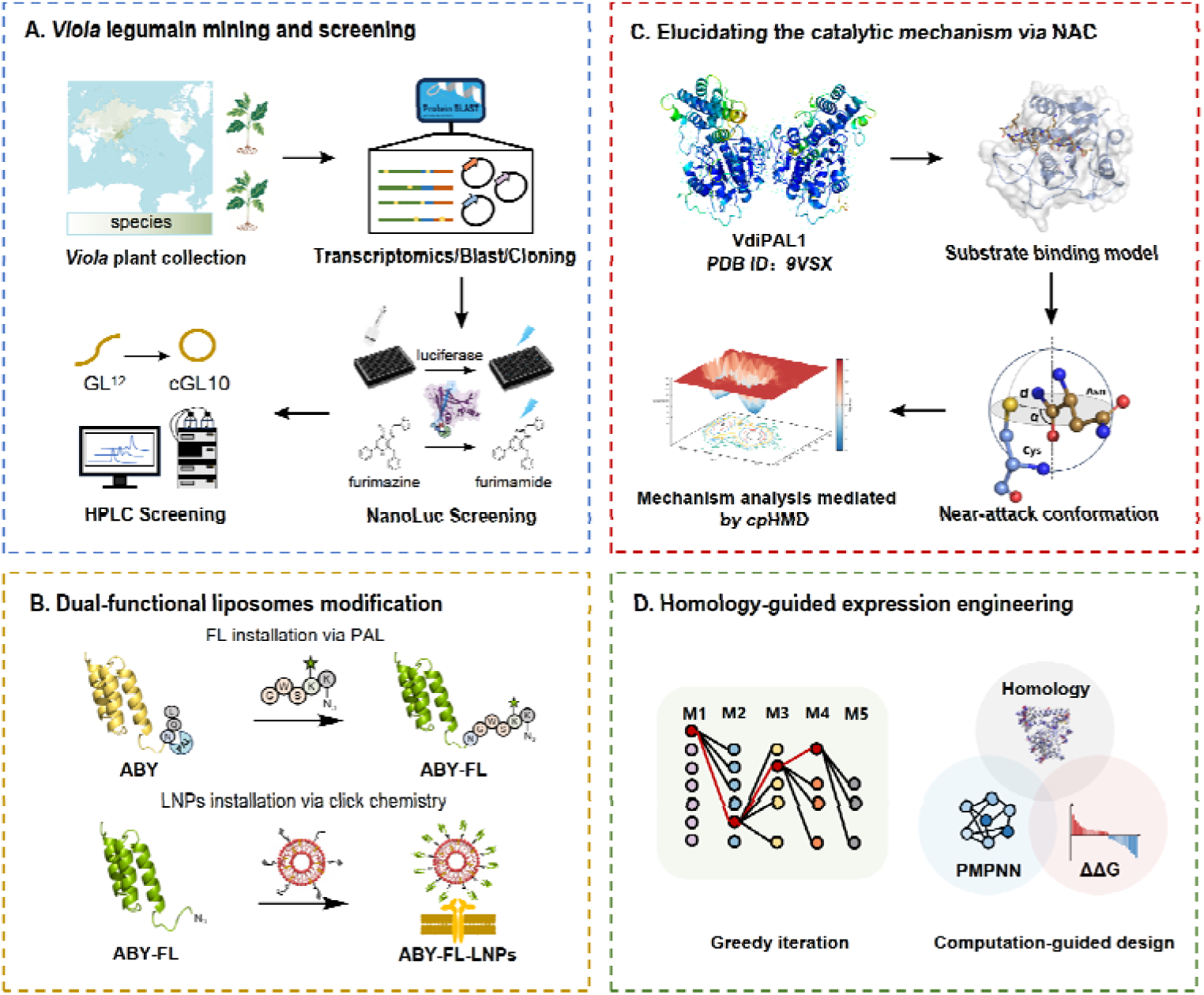
Integrated workflow for discovery, characterization, and engineering of *Viola*-derived PALs. (A) Schematic of the systematic mining, screening, and characterization of a novel legumain from *Viola* species. The workflow includes plant collection, BLAST analysis, sequence cloning, a NanoLuc-based primary screening, and subsequent characterization by HPLC. (B) Dual-functionalization of liposomes via PAL enzyme-mediated fluorescent-affibody preparation and one-pot click chemistry-mediated installation. (C) Elucidating the catalytic efficiency and pH-dependency via near-attack conformation (NAC). A NAC hypothesis was derived from the pre-organized conformation in crystal structure, and the prevalence of attack-favorable geometries was further supported in constant-pH MD (cpHMD) simulation. (D) Homology-guided expression engineering of VyPAL2. Key residues identified via multiple sequence alignment and single-mutant screening were subjected to both greedy accumulation strategy and computational-guided combinatorial mutagenesis.

## Results

### 1. Systematic mining of the *Viola* genus for shortlisted PALs

The *Viola* genus is recognized as a rich reservoir for cyclic peptides and an underexploited resource for peptide ligases. To systematically explore this resource, we procured 16 previously uncharacterized *Viola* species through extensive field collection and horticultural germplasm surveys from Global Biodiversity Information Facility (GBIF, https://www.gbif.org/). Together with seven previously reported *Viola* species^21, 25^, this curated collection of 23 species spans the native distribution of the genus across the Northern Hemisphere and Oceania (**Fig. 2A**), thus providing a geographically and climatically representative sampling of the approximately 400 known members of this genus^26^. Transcriptome sequencing of plant materials yielded a comprehensive legumain sequence library. We designed primers based on this dataset and successfully cloned 57 unique legumain-encoding genes (**Table S1**). Application of the LAD hypothesis enabled the prediction and naming of these sequences, giving 29 putative PALs and 28 putative AEPs (**Table S2**).

**Figure 2.**
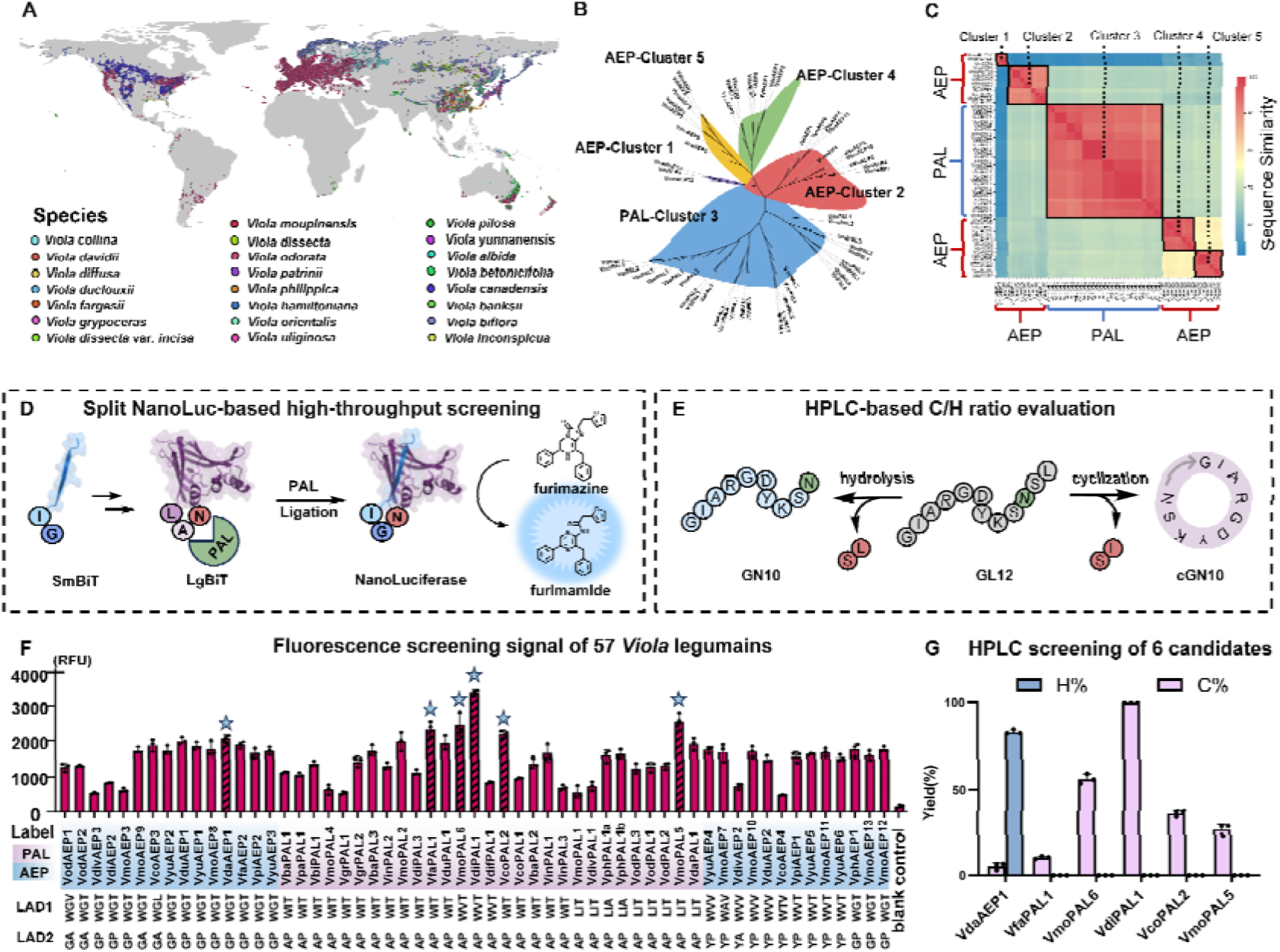
Sequence evolutionary features of legumains in newly collected *Viola* species. (A) Geographic and climatic coverage of 23 sampled *Viola* species. The species distribution map was compiled from publicly available occurrence records in GBIF and visualized using QGIS (Quantum Geographic Information System, version 3.40.9, https://qgis.org/). (B) Phylogenetic and sequence similarity analyses of 57 newly collected legumain sequences reveal a distinct PAL clade and four major AEP clades. (C) Sequence similarity matrix of AEP and PAL family. The heatmap displays the percentage of pairwise sequence identity, indicated by the color key on the right. The five major phylogenetic clusters (Cluster 1-5) identified in this study are outlined with black boxes. (D) Schematic of the split NanoLuc screening method. Blue GI-SmBiT and purple LgBiT-NAL are covalently joined by PAL to restore luciferase activity that converts furimazine into luminescent furimamide. (E) Schematic of the enzymatic reaction with substrate GL12 (gray). Legumain-mediated cyclization of GL12 results in cGN10 (pink) and hydrolysis results in GN10 (blue). (F) Discovery of high-performance VdiPAL1 via high-throughput dual-objective screening of *viola* legumains. Fluorescence-based ligase activity screening of 57 newly collected *Viola* legumains overlaid with phylogenetic clustering. Putative PALs were shown in purple and AEPs in blue. Top six enzymes exceeding a 2000 RFU threshold are marked with stars. (G) HPLC quantitation of the products catalyzed by the six best-performing candidates. Yield of GN10 (H%) was in blue. yield of cGN10 (%C) was in pink.

Sequence logo analysis of regions related to catalytic directionality, including LAD2, S2′, as well as the marker-of-ligase-activity (MLA) region^27^ show high conservation in the groups of putative PALs but less conserved in the group of putative AEPs (**Fig. S1**), agrees with previous discovery that plant AEPs are hydrolase with intrinsic transpeptidase potential^17^. Phylogenetic analysis further reveals that 29 putative PALs formed a tight, monophyletic clade with over 90% sequence identity, indicating an early evolutionary divergence that separates PALs from AEPs, followed by strong conservative selection within the *Viola* genus (**Fig. 2B**). In contrast, putative AEPs formed a separate branch that segregated into four distinct subclades, each exhibiting >80% internal sequence identity, a pattern suggestive of functional diversification supports the broad and distinct roles of legumains in plant life cycle^28^ (**Fig. 2C**). This high degree of conservation, alongside the broad geographical and climatic distribution of the source plants, suggests that our curated sequence library provides a representative snapshot of the natural diversity of *Viola* PALs.

To rapidly characterize the newly mined enzymes and screen for high-performance ligases, we adapted a split Nano Luciferase (NanoLuc^29^) reporter system previously described by Guo et al^30^. The split NanoLuc system was modified to carry PAL-specific motifs, as LgBiT-NAL and GI-SmBiT, respectively. Successful PAL-catalyzed ligation reconstitutes full NanoLuc activity, producing strong light emission at 460 nm by the conversion of substrate furimazine to furimamide (**Fig. 2D**). PALs are expressed as proenzymes composed by a core domain and a cap domain, which require acid-induced autocatalytic removal of a cap domain for their activation. Since PALs retain considerable ligation activity at pH 5.0, a condition that also permits activation, we integrated activation and activity detection into a single-step assay at pH 5.0. Using OaAEP1b-C247A^31^ proenzyme as a model, we confirmed a positive correlation between the luminescent signal and the amount and catalytic competence of the activated enzyme (**Fig. S2**), confirming this assay as a valid dual-objective screen for both expression and ligase activity. All 57 legumain candidates were cloned into pET28a(+) vector and expressed in *E. coli* SHuffle T7 in 24-well plates for high-throughput characterization. Crude cell lysate supernatants were directly subjected to the split-NanoLuc ligation assay. From this primary screen, six enzymes, including VdiPAL1, VmoPAL5, VmoPAL6, VfaPAL1, VcoPAL2, and VdaAEP1, yielded the highest luminescent signals, with VdiPAL1 from *Viola dissecta* being the top performer (**Fig. 2F**). A single amino acid difference between VinPAL1 and VinPAL3 resulted in a more than 2-fold difference in fluorescence signal, which we attributed to the variation in expression levels.

Since the split-NanoLuc assay cannot account for potential hydrolysis, the six candidates were subsequently analyzed by HPLC to specifically monitor the cyclization and hydrolysis of the peptide substrate GL12 (GIARGDYKSNSL, M.W = 1281.1 Da) (**Fig. 2E** and **S3**). Consistent with the LAD hypothesis, VdaAEP1 exhibited substantial hydrolytic activity, while the remaining five PALs were confirmed as promising ligases among which VdiPAL1 gave a near completion cyclization yield (**Fig. 2G**). Extended HPLC cyclization assays on other 24 putative PALs confirmed that all candidates exhibited cyclization activity with no detectable hydrolysis products except of 5 inactive ones (**Fig. S4**). These findings demonstrate the effectiveness of our dual-objective screening strategy by simultaneously selecting for high expression and catalytic activity. By combining rapid fluorescence-based screening with direct HPLC validation on crude supernatants, this approach bypassing laborious purification steps and enabling high-throughput screening.

### 2. VdiPAL1-mediated modular assembly of dual-functionalized liposomes

VdiPAL1, the top hit from our screen, was produced at a yield of 12 mg purified proenzyme per litre culture (**Fig. S5**). VdiPAL1 breaks the low-yield bottleneck of *Viola* PALs (0.1-2 mg L^-1^), showing significantly higher expression than the representative VyPAL2 (<0.5 mg L^-1^) **(Fig. S6)**. While its yield is slightly lower than the benchmark OaAEP1b-C247A (15 mg L^-1^) **(Fig. S7)**, VdiPAL1 offers a critical advantage by minimizing the hydrolysis by-products associated with OaAEP1b (**Fig. S8**), thus making it the most promising candidate for peptide ligation. In the cyclization of GL12 at pH 6.5 and 37 °C, VdiPAL1 exhibited a *k_cat_* similar to that of the variant VyPAL2-I244V^32^ (1.83±0.14 vs. 1.81±0.09 sL¹) and a slightly higher catalytic efficiency (*K_m_* = 4.14±1.31 vs. 5.77±0.99 μM; *k_cat_/K_m_*= 4.42 × 10L vs. 3.14 × 10L ML¹ sL¹) (**Fig. 3A**). Specificity profiling at the P2’’ position showed that VdiPAL1 has an extended substrate preference, efficiently accepting Trp, Met, and Tyr, which was broader than the previously reported scope including Leu, Ile, Val, Cys, and Phe^18, 19, 33^ (**Fig. 3B**). Interestingly, this extended preference was also observed in VyPAL2-I244V and OaAEP1b-C247A under identical conditions, which was not previously reported for these ligases (**Fig. S8**). This suggests that P2’’ tolerance may be more conserved across PALs than previously recognized and less stringent than P2’ for substrate recognition. Thermal stability assays comparing VyPAL2 and VdiPAL1 across pH 4-8 showed that VdiPAL1 has a Tm value of 41.5°C at pH 6.5, which is superior to that of VyPAL2-I244V (38.3°C) (**Fig. S9**). Taken together, despite its high sequence similarity to known PALs, VdiPAL1 is a unique wild-type PAL that outperforms both the widely used OaAEP1b-C247A and VyPAL2-I244V, by combining superior catalytic efficiency, high expression yield, extended substrate specificity, and enhanced stability, making it an ideal candidate for advanced bioengineering applications.

**Figure 3.**
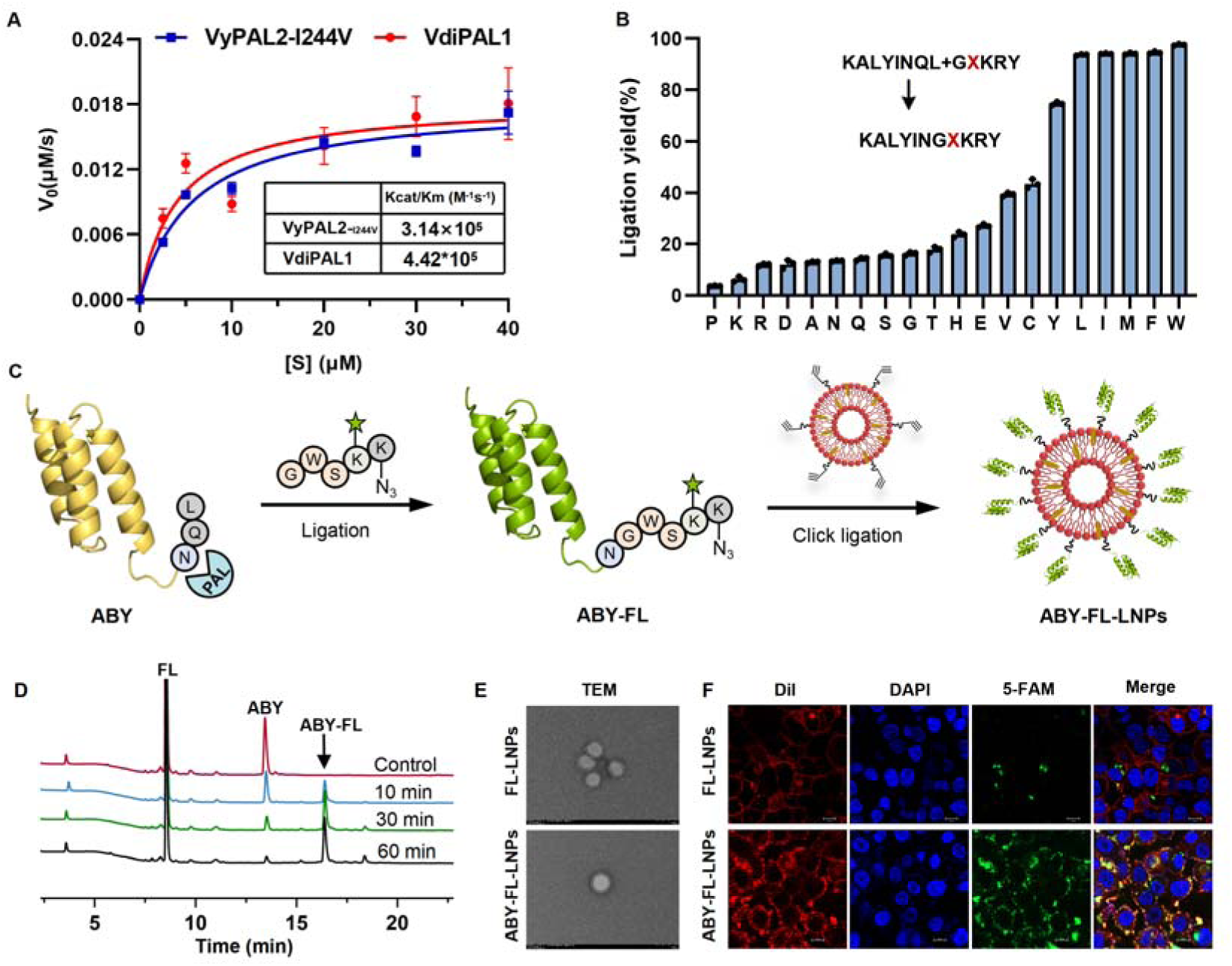
Characterization and modular assembly of EGFR-targeting fluorescent LNPs via VdiPAL1-click strategy. (A) Kinetic comparison of VyPAL2-I244V and VdiPAL1 in catalyzing peptide cyclization. (B) Substrate specificity of VdiPAL1 at the P2’’. Ligation yields are shown for reactions between KALYINQL and GXKRY peptides, where X represents each of the 20 standard amino acids. To enhance catalytic efficiency, a glutaminyl cyclase (QC) (**Fig. S10**) enzyme was co-introduced to minimize the reverse reaction^35^. Bars represent the mean of three independent experiments. (C) Schematic workflow for modular assembly of affibody-fluorescein dual-functionalized LNPs via VdiPAL1-mediated ligation and CuAAC click chemistry. (D) Efficiency of the VdiPAL1-mediated ligation analyzed by HPLC. (E) TEM images of FL-LNPs and ABY-FL-LNPs. (F) Confocal microscopy images showing the cellular uptake of FL-LNPs and ABY-FL-LNPs by A549 cells.

To demonstrate the versatility of VdiPAL1 in bioconjugation, we developed an efficient chemoenzymatic strategy combining VdiPAL1 mediated ligation with click chemistry^34^. This approach was used to generate affibody/fluorescein dual-functionalized liposomal nanoparticles (ABY-FL-LNPs) for targeting the overexpressed epidermal growth factor receptor (EGFR) on cancer cells (**Fig. 3C**). We ligated a recombinantly expressed zEGFR-affibody (ABY) (**Fig. S10**) bearing an NQL motif to an FL-peptide (GWSK(5-FAM)K(N_3_)) via PAL catalysis. The reaction was carried out at pH 6.5 and 37 °C with 400LnM VdiPAL1, 200 μM ABY, and 400 μM FL (a 1:500:1000 molar ratio), achieving 90% conversion within 60 minutes (**Fig. 3D** and **S11**). After centrifugal filtration to remove excessive FL-peptide, ABY-FL was directly conjugated onto alkyne-containing liposomes via Cu-catalyzed click reaction. Dynamic light scattering (DLS) (**Fig. S12**) and Transmission Electron Microscope (TEM) (**Fig. 3E** and **S13**) confirmed that the final ABY-FL-LNPs maintained their morphology with an average diameter of 140 nm. Cellular uptake studies on A549 (EGFR^+^) cells showed that ABY-FL-LNPs were successfully internalized. In contrast, uptake of FL-LNPs was limited, confirming the crucial role of affibody on EGFR-targeting (**Fig. 3F**). This PAL-click strategy provides an orthogonal and modular platform for the programmable assembly of functional LNPs as both the targeting and tracing modules can be readily changed. The approach is broadly compatible with diverse liposome formulations and allows for pre-loading with therapeutic cargos, enabling the construction of target-specific and traceable nanomedicines.

### 3. Structural basis of pre-organized catalytic geometry and pH-dependent reaction mechanism

To elucidate the structural basis for its high efficiency of VdiPAL1, we determined the crystal structure of the recombinant proenzyme at a resolution of 1.87 Å (PDB ID 9VSX, **Fig. 4A, S14,** and **Table S3**). The structure reveals dimer formation, with each core domain monomer displaying the canonical αLβL legumain fold (**Fig. S15**). Structural alignment with VyPAL2 (PDB ID 6IDV) shows a Cα root-mean-square deviation (RMSD) of 0.74 Å. Notably, the high resolution allowed the first identification of a stably bound water molecule within the catalytic cleft between the core and cap domains (**Fig. 4A** and **S16**). This water molecule is coordinated through a hydrogen bond network with four highly conserved residues, including three S1-pocket-forming residues Glu211, Ser241, and Asp262 from the core domain, and Gln342 from the cap domain that protrudes into the active site. Mutation of these residues either abolished activity or reduced expression level (**Fig. S17**). We propose that this hydrogen-bond network plays a crucial role in stabilizing the proenzyme conformation by strengthen the cap domain anchoring to the core.

**Figure 4.**
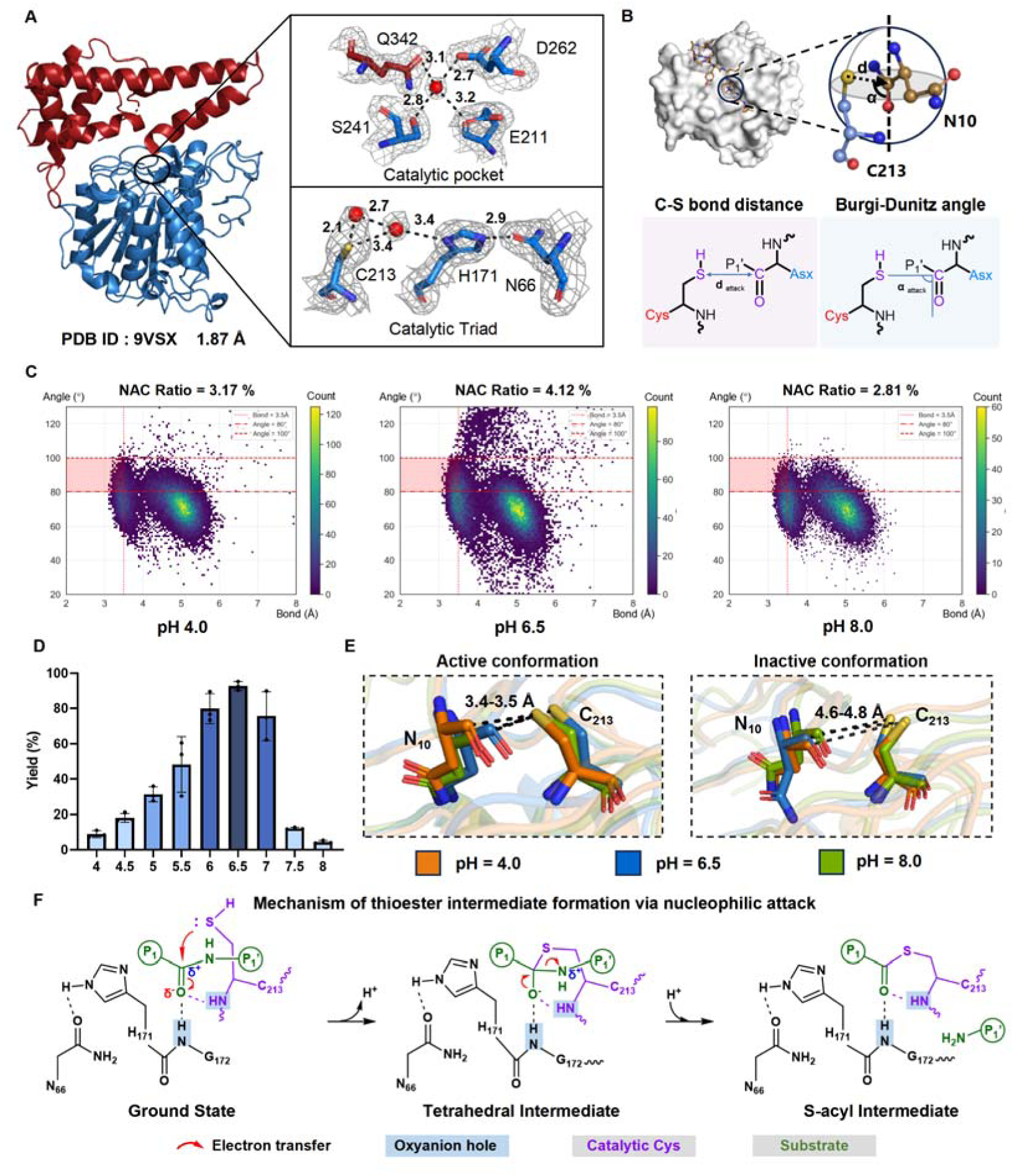
Structural and catalytic mechanism of VdiPAL1. (A) High-resolution crystal structure of VdiPAL1 proenzyme (PDB ID: 9VSX) reveals three stable water molecules forming hydrogen bond with the S1 pocket residues and with the catalytic triad C213-H171-N66 within the cleft between the cap (red) and core (blue) domains. The red balls represent water molecules. The gray mesh shows the 2*F*o-*F*c electron density map at a contour level of 1.0 r.m.s.d. (B) Structural depiction of NAC, highlighting the electrophile (carbon)-nucleophile (sulfur) distance and the Bürgi-Dunitz angle between the substrate Asn (brown) and the catalytic Cys (blue). Schematic below shows the two-dimensional NAC geometry parameters. (C) Distribution heatmaps of NAC distances (C-S) and angles from cpHMD simulations at pH values 4.0, 6.5 and 8.0. NAC-compliant populations are boxed in red. (D) Cyclization yields of VdiPAL1-catalyzed reaction across pH 4–8. (E) Representative conformational snapshots of VdiPAL1 and GL12 complexes from cpHMD simulations at pH 4.0 (orange), 6.5 (blue), and 8.0 (green). (F) Mechanism of thioester intermediate formation via nucleophilic attack. The substrate is shown in green and the catalytic Cys213 in purple. Key residues opposite the scissile bond (His171, Asn66, Gly172) are shown in black. The oxyanion hole, formed by backbone amides of Gly172 and Cys213, is boxed in blue. Electron flow is indicated by red arrows. Schematic of initial attack by Cys forms an enzyme-substrate thioester intermediate. The substrate is shown in green, catalytic Cys213 in purple, residues at opposite site in black (His171, Asn66, Gly172), and the oxyanion-stabilizing amides on G213, C213 are boxed in blue. Electron flow is indicated by red arrows.

In the active center, we observed another two stable water molecules bridging the catalytic residues His171 and Cys213 in a “pre-organized” geometry (**Fig. 4A**). Especially, superimposed with the substrate-bound-mimicking structure of VyPAL2 (PDB 7F5P, **Fig S18**), Cys-Sγ is 2.5 Å from the carbonyl carbon of substrate P1 residue, matching the catalytically competent state. In contrast, His-Nδ is 3.5 Å from the P1-corbonyl carbon and 3.9 Å from the P1’ amine group, suggesting a much weaker potential for direct catalysis.

Unlike most serine and cysteine proteases, which typically rely on a tightly coupled catalytic dyad or triad to mediate proton transferring and sequential nucleophilic attacks, PALs exhibit a spatial separation between the catalytic dyad Cys and His, positioning on opposite sides of the substrate scissile bond. This geometric arrangement suggests that the nucleophilic attack by Cys may proceed independently of His. Interestingly, previous mutagenesis studies indicated that Ala substitution of the catalytic Cys could retain weak activity, particularly the acid-induced autoactivation, raising the possibility that His alone may act as an alternative nucleophile in the initial acylation step in the absence of Cys^32^. Computational enzyme design has also validated that His could act as an independently catalytic nucleophile^36^.

To assess the relative competence of these residues and determine the major nucleophilic residue, we conducted both computational simulation and mutagenesis study. We constructed substrate-enzyme complex models and evaluated the geometric parameters associated with the NAC, including distance (*d*_attack_) between the nucleophile and the substrate’s carbonyl carbon and the Bürgi-Dunitz angle^37^ (*a*_attack_) (**Fig. 4B**). Given that a C–S bond length of ∼1.8 Å^38^ is too restrictive for the initial encounter complex, we used van der Waals constraints (*d*_attack_ < 3.5 Å)^39^ and attack-ready pre-reactive geometry angle (*a*_attack_ = 90° ± 10°) based on a recent reported model of serine protease as proxies for productive nucleophilic pre-organization^40^. In addition, to reflect the pH-dependency of PAL activity, which is higher at a near-neutral environment and reduced under acidic or basic conditions, we performed constant-pH MD (cpHMD) simulations^41,42^ at pH 4, 6.5, and 8 to monitor the changes of *d*_attack_ and *a*_attack_.

In 15 independent 100 ns cpHMD simulations performed at each pH and assuming the catalytic Cys thiol as the nucleophile, the fraction of frames adopting NAC (*d*_attack_ (C-S) < 3.5 Å and α_attack_ = 90° ± 10°) peaked at pH 6.5 (4.12%), and decreased markedly at pH 4.0 (3.17%) and pH 8.0 (2.81%) (**Fig. 4C, S19-S21**). This pH-dependent trend aligns closely with experimentally measured cyclization yields (**Fig. 4D**), validating NAC frequency as a structural correlate of catalytic competence. We also found that the side chain of the catalytic Cys residue mainly adopts two conformations. In the active state, the thiol is oriented toward the substrate’s carbonyl, positioning it for nucleophilic attack, whereas in the inactive state, the thiol group is rotated to the opposite direction, increasing the distance to the carbonyl carbon by more than 1 Å (**Fig. 4E**). In contrast, simulations assuming His as the nucleophile yielded markedly lower NAC frequencies across all pH conditions and failed to reproduce the observed pH-depended activity profile in 100 ns (**Fig. S22-S26**). These data argue against a primary nucleophilic role for His in the presence of catalytic Cys. Prior computational studies on human legumain^43^ and caspases^44^ suggest that His may function in stabilizing the oxyanion intermediate or facilitate proton transfer to the leaving amine. To further probe the role of His, we mutated it to Ala in MD simulations and observed a significantly lower substrate binding affinity that caused C-S distance increased from average 4.6 Å to 11.9 Å, indicating a loss of NAC pre-organization (**Fig. S27**). In MM/PBSA analysis, the average binding free energy for the H171A mutant complex was −29.9 kcal/mol, indicating a substantially weakened interaction compared to the −67.9 kcal/mol calculated for the wild-type complex (**Fig. S28**). These results suggest that the critical roles of His171 in catalysis, likely include stabilizing the substrate in NAC, a geometric arrangement crucial for high catalytic efficiency. This hypothesis is further supported by Alanine scan of the theoretical catalytic triad, N66A, H171A and C214A (**Fig. S17**). Among them, N66A failed to be expressed, while both H171A and C214A retained low but detectable activity. Validated by one-pot auto-activation and substrate-cyclization reactions (30 °C, pH 4.0, overnight) with high concentration of purified proenzymes. Notably, H171A achieved full substrate conversion despite being much more difficult to activate than C213A (**Fig. S29**-**S30**), which left ∼15% unreacted substrate (**Fig. S31**). This finding supports the hypothesis that His can serve as an alternative nucleophile in the absence of catalytic Cys (**Fig. S32**) and the lower substrate conversion rate of C213A reaffirms the role of Cys as the primary catalytic nucleophile. Furthermore, the impaired activation of H171A underscores the critical role of His in the substrate positioning, which was consistent with the observed loss of substrate pre-organization and weakened binding affinity in the MD simulations and MM/PBSA analysis.

To further support the reliability of the NAC analysis, we extended the simulation time to 1 μs for three independent replicates. Similarly, the NAC ratio using Cys as the nucleophilic residue at the three pH levels captures the experimental observations (**Fig. S33-S34**), while the scenario with His as the nucleophile fails to reproduce the experimental trend (**Fig. S35-S36**). Unlike the 100 ns simulations, extended simulations revealed a relaxation process within the system, with the NAC proportion reaching convergence by 600 ns (**Fig. S37**). Notably, despite the observed decrease in the absolute NAC proportion in the extended simulations, the pH-dependent trend (pH 6.5 > pH 4.0 > pH 8.0) remained consistent with both the 100 ns simulations and experimental results.

Collectively, these results support a model in which catalytic Cys executes the nucleophilic acylation step, while His and Asn contribute to catalysis by stabilizing the substrate conformation other than proton transfer. The oxyanion hole, formed by the backbone amides of C213 and G172, is also essential for stabilizing the negative charge developed during the transition state (**Fig. 4F**). This pre-organized NAC-assisted mechanism, distinct from classical serine/cysteine protease triads, underlies the high catalytic proficiency and unique pH dependence of PALs.

### 4. VdiPAL1-informed rescue of VyPAL2 expression via cap redesign

Despite sharing high sequence and structural homology, VdiPAL1 and VyPAL2 exhibit dramatically different expression profiles in *E. coli*. VyPAL2 is an attractive PAL owing to its minimal hydrolytic side activity, yet its practical application is hindered by poor prokaryotic recombinant yields (<0.5 mg L^-1^) and the need for costly insect cell expression system. To address this, we employed the highly expressible VdiPAL1 as a template to guide VyPAL2 engineering.

Sequence alignment identified 30 amino acid differences between the two enzymes (**Fig. S38**), To pinpoint key residues influencing expression, we construct 30 single-residue-substitution mutants of VyPAL2 and used a GFP fusion reporter system (**Fig. S39**) for high-throughput screening^45^. 19 of the 30 mutants exhibited higher fluorescent signals than the wild-type, with the M357N mutant showing the most significant increase (1.4-fold, **Fig. 5A**). Notably, seven of the top ten expression-enhancing mutations located in the cap domain, predominantly at the termini of helices or loops, suggesting a role in modulating local folding or cap dynamics.

**Figure 5.**
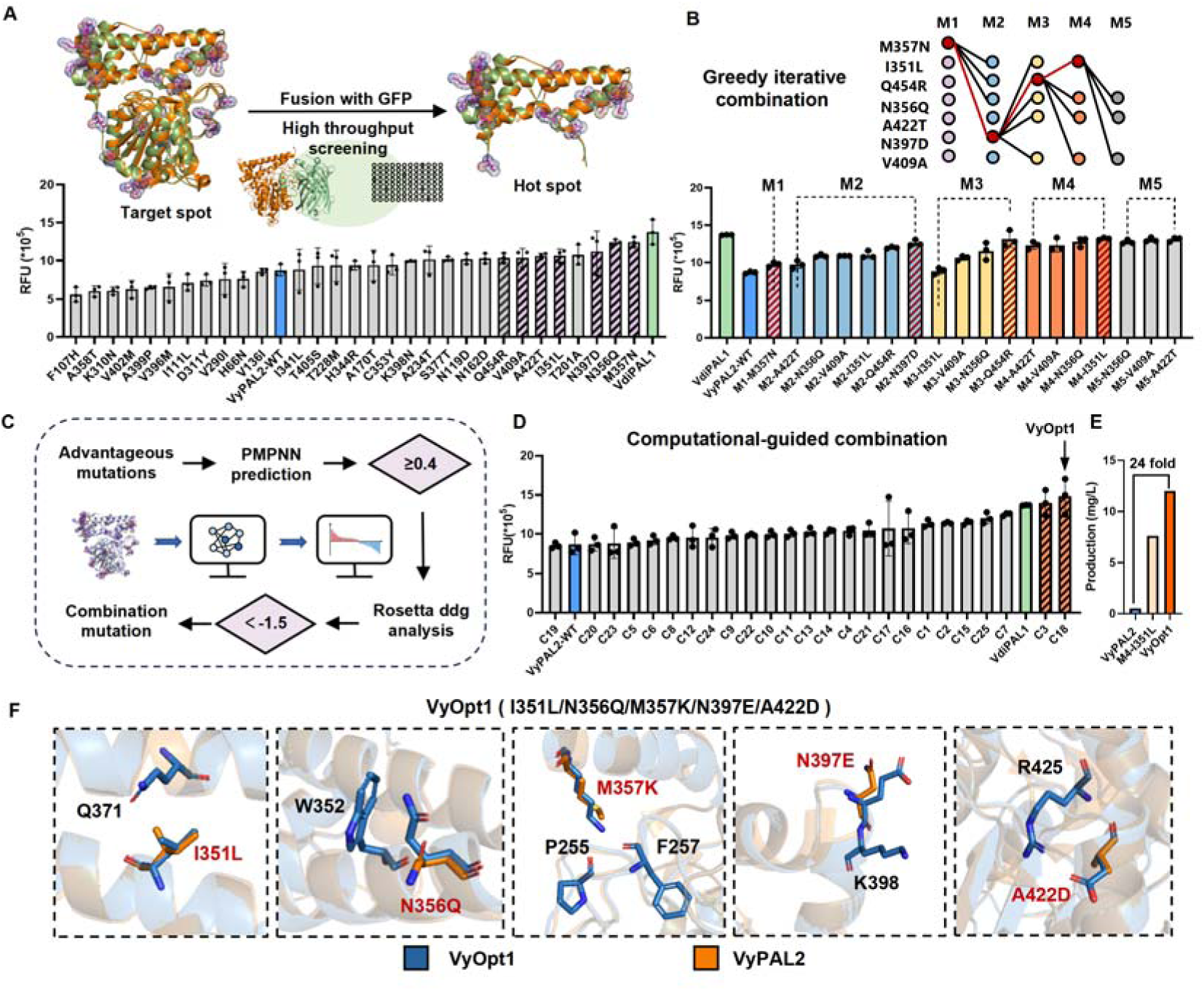
Homology-guided engineering of VyPAL2 to enhance recombinant expression. (A) High-throughput screening of 30 divergent residues between VyPAL2 (blue bar) and VdiPAL1 (green bar) using a GFP-fusion reporter resulted in the identification of seven cap-domain hotspots. Candidate hotspot mutations are highlighted with purple hatched bars. (B) Schematic of greedy combinatorial mutagenesis. Beneficial mutations (red dots) were iteratively combined over five rounds and reached peak after round four, yielding a quadruple mutant M4-I351L. Top-performing mutant in each round is highlighted with hatched bars. (C) Computational pipeline for enhanced expression design. Random combinatorial mutations at hotspot residues were predicted using ProteinMPNN and subsequently filtered based on experimental data of advantageous single mutations and Rosetta_ddg predicted ΔΔG, resulting in 25 candidate combinations. (D) Experimental validation of 25 top candidates from computational design. (E) Expression yield comparison among the wild-type VyPAL2 (blue), the conventionally iterated quadruple mutant M4-I351L (light orange), and the computationally designed combinatorial variant VyOpt1 (dark orange). (F) Structural comparison of the mutation sites in VyOpt1 (blue) versus wild-type VyPAL2 (orange) which improve packing by introducing new hydrogen bonds and electrostatic interactions.

To further enhance VyPAL2 expression while maintaining its native activity, we focused on combinatorial integration of the seven beneficial cap-domain mutations. A widely-applied greedy mutagenesis strategy^46^ was implemented, starting with the top-performing single mutant M357N and progressively incorporating additional mutations through iterative pairwise combinations. After four rounds, fluorescent signal reached peak with a quadruple mutant, VyPAL2-I351L/M357N/N397D/Q454R. Following removal of the GFP tag, this engineered variant achieved a soluble yield of 7.6 mg/L (**Fig. S40**), representing a 15-fold increase over the wild-type enzyme (**Fig. 5B**).

To overcome the local-optimum limitations often encountered in conventional iterative mutagenesis, we implemented a computation-guided strategy in parallel to explore a broader combinatorial landscape. Based on our observation that beneficial variants were enriched in the cap domain, we hypothesized that enhanced foldability of the cap domain and stabilized cap-core interactions could underlie improved recombinant expression. To test this hypothesis, we devised a rational design strategy that integrates our experimental screening data with computational modeling. We began by leveraging our advantageous single-mutation data to define a focused set of seven hotspot sites. This experimentally-constrained set was served as the input for ProteinMPNN to generate a comprehensive library of 2–5 site combinatorial mutations. To retain evolutionary relevance, we ensured that at least 40% of mutated sites in each variant were substituted with VdiPAL1 residues. 265 candidates generated were then subjected to Rosetta_ddg to estimate the change in folding free energy (ΔΔG) upon mutation^47^. The 25 top-scored variants (ΔΔG < −1.5 REU) were selected for experimental validation (Table S4, **Fig. 5C**). Among these, a quintuple mutant VyOpt1 (I351L/N356Q/M357K/N397E/A422D) gave the highest signal (**Fig. 5D**). After GFP removal and process optimization, VyOpt1 reached a final yield of 12 mg L^-1^ (**Fig. 5E**, **S40** and **S41**), representing 24-fold improvement over the wild-type VyPAL2 and significantly outperforming the 7.6 mg L^-1^ obtained by greedy combinatorial approach.

Structural modelling of VyOpt1 provide mechanistic insights into its improved expression. The I351L mutation replaces a β-branched isoleucine with a more conformationally flexible γ-branched leucine, potentially reducing steric hindrance and introduces a stabilizing hydrophobic interaction with Q371. The N356Q mutation, characterized by an extended side chain, shortens the hydrogen bond distance to W352, thereby strengthening the interaction. At the α-helix turn, M357K stabilizes the local structure through hydrogen bonds formed between the Lys side chain and residues P255 and F257. Similarly, the N397E mutation, located at a loop turn, enhances surface hydrophilicity, contributing to improved solubility and stability. Finally, A422D reshapes the local hydrogen bond network by forming new hydrogen bonds and salt bridges with Arg425, significantly increasing structural rigidity^48^. In summary, these residues enhance the stability of the cap domain, thereby facilitating the folding of the core domain and increasing overall enzyme expression (**Fig. 5F and S42-S45**). To our knowledge, this is the first report that explicitly links cap-domain stabilization to improved expression in PALs.

## Discussion

By retracing the evolutionary landscape and functional limitations of previously characterized PALs, our study reinforces the value of mining natural diversity for more efficient and expressible ligases. Drawing on the botanical richness of China, we surveyed together 23 *Viola* species spanning the Northern Hemisphere and Oceania, effectively covering the principal biogeographic clades of the genus, offering a representative overview of *Viola* legumains. Phylogenetic analysis revealed a distinct clade of PALs with high conservation associated with the cyclotide biosynthesis and 29 new PAL sequences were identified. Among them, VdiPAL1 stood out for its high soluble yield (12 mg L^-1^) and robust ligation activity, enabling efficient chemoenzymatic dual-functionalization of liposomes. These finding highlight how systematic exploration of natural diversity can yield functionally superior enzymes, even within evolutionarily conserved families. These results suggest that PALs from diverse plant lineages share a conserved tolerance for bulky hydrophobic residues at the P2’’ position. Notably, the observed acceptance of Met and aromatic Trp/Tyr, which were not favored at the corresponding P2’ position of the leaving group. This finding raises the possibility that S2’’ and S2’ sites are not fully overlapping, with the S2’’ site exhibiting a broader substrate tolerance. This expanded acceptance spectrum at P2’’ opens new opportunities for designing orthogonal P1’-P2’ and P1’’-P2’’ substrate pairs, potentially enabling multi-site peptide ligation and diversification in synthetic applications.

To dissect the basis of the efficiency and pH-dependency of PALs, we combined crystallography, cpHMD simulations and mutagenesis. While classical serine and cysteine proteases rely on well-defined NACs involving a hydrogen-bonded Ser/Cys-His-Asp triad and proton-relay coupling^49^, the Cys-His dyad in PALs lacks direct hydrogen bonding yet still pre-organized into an NAC stabilized by bridging water molecules, as revealed by the high-resolution structure of VdiPAL1. This pre-organization of the catalytic Cys-His dyad likely lowers the entropic barrier for substrate engagement and primes the catalytic pocket for nucleophilic attack, which may account for the unusually low K_m_ and high *k*_cat_ of PALs. Mutagenesis and simulations confirmed Cys as the primary catalytic nucleophile, while His act mainly as the substrate-stabilizer rather than a general base when Cys is present. Moreover, the oxyanion hole, formed by the backbone amides of Gly172 and Cys213, likely pre-activate the carbonyl carbon by enhancing its electrophilicity. This may underlie the low but measurable activity observed in the Cys-to-Ala mutant, supporting a model where His can function as a secondary nucleophile under favorable electrostatic preorganization.

Simulation of the enzyme-substrate complex under varying pH showed that catalytically competent geometries indicated by optimal C-S bond distance and Bürgi-Dunitz angle were most enriched at pH 6.5, closely matching with the experimental activity profiles. These results suggest that the local electrostatics modulate NAC ensemble and thereby tune the catalytic efficiency across pH ranges. To further support the reliability of the NAC analysis, we extended the simulation time to 1 μs (**Fig. S36**). These longer simulations revealed a relaxation no later than 600Lns, with cumulative NAC occupancy reaching equilibrium (**Fig. S37**). This delayed convergence indicates that while the initial AlphaFold3 predicted structure was close to the NAC conformation, it still required solvent reorganization and side-chain rearrangements to reach true dynamic equilibrium. Although the absolute NAC frequencies decreased compared to 100Lns simulations (<2% vs. <5%), the pH-dependent trend (pH 6.5 > 4.0 > 8.0) remained consistent and agreed with the experimental results. A literature survey confirms that NAC fractions reported in MD simulations vary significantly across enzyme systems (ranging from <1% to >70%) and low NAC fractions are common^50^, since NAC is transient near-attack geometry rather than a stably populated ground state. For example, MD study of HIV-1 protease reported NAC populations as low as 0.042% for bound water nucleophiles^51^. Therefore, given the strict geometric criteria applied in this study, the observed NAC frequencies fall within a scientifically reasonable range and reflect the intrinsic energetic barrier associated with this catalytic step. Considering the critical role of the oxyanion hole, we also re-evaluated our simulations by adding both N-O van der Waals radii sum in the NAC criteria. Consequently, the 100 ns simulation was not yet converged but the 1 μs simulation retained the pH-dependent trend in-line with the experimental data (**Fig. S46**). Although substrate-bound crystal structure could not be obtained after multiple attempts (data not shown), likely due to rapid turnover and conformational flexibility, converging evidence from kinetics, high-resolution structure, and extended cpHMD simulations support a model in which pre-organized active-site geometry and electrostatic environment cooperatively stabilize the NAC and govern pH-dependent reactivity.

Besides the enzyme side, the protonation state of the substrate also critically influences the ligation outcome. In principle, the incoming peptide nucleophile prefers to be in the neutral -NHL form rather than protonated -NHLL to perform a nucleophilic attack, which thermodynamically favors mildly basic conditions. However, the observed decline in ligation efficiency at higher pH suggests that the mere presence of a reactive nucleophile is insufficient. At alkaline pH, the preorganized NAC geometries become less populated, likely due to conformational perturbations of the active site. Thus, efficient catalysis requires a dual prerequisite: both a deprotonated nucleophile and a conformationally competent enzyme pocket. The optimal pH (6.5) represents a delicate balance between these two factors, where substrate reactivity and enzyme geometry are simultaneously satisfied.

Beyond mechanistic insight, VdiPAL1 served as a template to improve the poor expression of Viola PALs. Through homology-guided hotspot scanning in VyPAL2, we identified key residues in the cap domain, particularly at loop termini as potential modulators of recombinant expression. The greedy iterative combination reached a plateau (7.6 mg L^-1^) after four rounds, prompting us to explore a computation-guided strategy to expand the combinational search space. We integrated experimentally identified beneficial sites with ProteinMPNN for sequence regeneration and Rosetta_ddg for stability-based filtering. This pipeline yielded a quintuple mutant, VyOpt1, with a 24-fold expression increase in just one design-build-test cycle. The final design combined two mutations directly derived from VdiPAL1 and three MPNN-generated substitutions not found in natural homologs, underscoring the ability of deep learning models to propose beneficial mutations that improve foldability beyond the natural sequence landscape^52^. This method leverages the synergy between sequence homology and structural stability, enabling rapid prioritization of combinatorial mutations that optimize expression with minimal experimental iterations.

Analysis of these beneficial mutation sites points to cap-domain foldability as a pivotal determinant of expression efficiency. This insight extends the known function of the cap domain beyond its canonical role as conditional activation control. We propose that the cap also acts as a structural scaffold essential for proper folding and solubility of the PAL proenzyme. This concept not only explains the divergent expression levels of homologous PALs but also offers a broadly applicable strategy for enhancing the production of other difficult-to-express PALs through targeted cap engineering.

In summary, this work provides an integrated workflow from sequence mining to enzyme optimization. We expanded the known diversity of peptide asparaginyl ligases, identified a highly efficient and expressible nature VdiPAL1, uncovered structural and catalytic features underlying its activity, and established a generalizable strategy for improving expression of homologous enzymes. Overall, these advances will facilitate further mechanistic and engineering studies of PALs, promoting their development as biosynthetic bricks for versatile peptide functionalization.

## Materials & Methods

### Sequencing and transcriptome analysis

Fresh *Viola* plants were obtained through field collection and horticultural seed collection. Fresh plant leaves were sent to BGI Genomics for RNA sequencing, followed by data processing steps (assembly and annotation) to obtain transcriptomes. CDS annotated to be legumain, AEP, or vacuolar processing enzymes were cross-validated by their sequence similarity with VyPAL2. Candidate genes were used as templates for the design of cloning primers (**Table S1**).

### RNA extraction, cloning, recombinant expression and protein extraction

Fresh leaves were cut to extract plant mRNA using Trizol method, and cDNAs were obtained by reverse transcription using a HiScript III 1st Strand cDNA Synthesis Kit (Vazyme, R312-01, Nanjing, China) and then used for cloning of legumain sequences. Legumain sequences were amplified by PCR and cloned into pET28a(+) vectors using ClonExpress II One Step Cloning Kit (Vazyme, C112-01) with NdeI/XhoI restriction enzymes in-frame with both N-terminal and C-terminal His6-tags. Plasmids were transformed into *E. coli* DH5α competent cells for amplification. After sequencing, correct legumain-encoding plasmids (**Table S2** and **supplementary dataset 1**) were transformed into *E. coli* SHuffle T7 competent cells, and the transformants were picked and incubated with LB/Kanamycin medium to obtain the seed solution. 10 uL overnight cultured seeds were added 24-well plate containing 1 mL LB/Kanamycin medium and continued incubating at 37 L, 220 rpm until the OD_600_ reached 0.4. Induction of the target proteins was performed with 0.1 mM IPTG at 16 L, 180 rpm for 24 to 48 h and cells were harvested by centrifugation at 4000 *g*, 30 min at 4°C. The cell pellets were resuspended in the lysis buffer (50 mM HEPES, 0.1 M NaCl, 1 mM EDTA, 5 mM β-mercaptoethanol (β-ME), 0.1% TritonX-100 at pH 7.5, 0.2 mg mL^-1^ lysozyme, and the crude enzyme supernatant was collected by centrifugation after overnight lysis.

### Split-NanoLuc activity screening

Highly expressible OaAEP-C247A was used for condition optimization of the split-NanoLuc assay. Ligation reactions (25 μL) were prepared with 25 μM LigBiT and 100 μM SmBiT in 20LmM citrate buffer (pH 5.0) containing 1 mM EDTA and 5 mM β-mercaptoethanol. Crude enzyme (1, 2, 4, or 8 μL of cell lysate supernatant) was added to initiate the reaction, and mixtures were incubated at 37 °C for 30 or 60 min. For luciferase activity detection, 2-8 μL of the ligation product was transferred into 100 μL of PBS containing 5 μM furimazine, and chemiluminescence was measured immediately using a microplate reader (**Fig. S2**). Based on signal-to-background ratio, the final optimized condition was determined as 1 μL of crude enzyme, 37 °C incubation for 60 min, followed by transfer of 4 μL reaction product into 100 μL of detection buffer for measurement.

### Scale-up expression, purification and acid-induced auto-activation of VdiPAL1

PALs were expressed using the procedure mentioned above in a 1 L scale. A volume of 10 mL lysis buffer was added to resuspend every 1 g cell pellet. After sonication on ice (40% power, 1 s on / 3 s off cycle for total 30 min), cell debris was removed by centrifugation at 12,000 x *g* for 30 min at 4 °C. The supernatant was filtered using 0.45 μm syringe filter and loaded onto a nickel column (Cytiva, HisTrap HP 1 mL, Uppsala, Sweden) for His-tag affinity purification (Cytiva, AKTA Start^TM^ purification system). Target protein was eluted with elution buffer (50 mM HEPES, 0.1 M NaCl, 1 mM EDTA, 5 mM β-ME, 0.5 M imidazole, pH 7.5). The eluted fractions were concentrated and buffer-exchanged with binding buffer (50 mM HEPES, 0.1 M NaCl, 1 mM EDTA, 5 mM β-ME, pH 7.5) using a ultrafiltration device (Amicon® Ultra, 10 kDa MWCO, Merck) to remove excessive imidazole. Purified proenzymes in a concentration of 0.2 mg mL^-1^ were activated by lowering the pH to 4.2 and incubating at 30°C for 30 min. Activated enzymes were purified on a HiLoad 16/600 Superdex 200 column with AKTA pure 25L protein purification system (Cytiva) with 20 mM sodium citrate buffer, 1 mM EDTA, 5 mM β-ME, 0.1 M NaCl and 5% (w/v) glycerol, pH 4.2. The concentration of purified enzymes was determined by measuring the absorbance at 280 nm using microvolume spectrophotometer (Yomim Instruments, Unano-1000, Hangzhou, China). The eluents were neutralized to pH 6.5 and stored at 4 °C or −80 °C after the addition of 20% sucrose.

### Kinetic and thermal stability analysis of VdiPAL1 and VyPAL2-I244V

In vitro cyclization assays were performed in 50 μL volumes containing 10 nM VdiPAL1 or VyPAL2-I244V, 2.5-40 μM substrate GL12 (GIARGDYKSNSL), and reaction buffer (20 mM phosphate buffers, pH 6.5, 1 mM EDTA, 5 mM β-ME). Reactions were performed in triplicate at 37 °C and quenched at 30, 60, 90, and 120 s by addition of 2 μl 1 M HCl. Cyclization rates were calculated by converting the HPLC peak areas of the remaining linear precursors or the cyclized products into concentrations. Initial velocities were plotted using GraphPad Prism 8.0 to obtain the Michaelis-Menten curve and the kinetic parameters (*K*_cat_ and *K*_m_) for each enzyme (**Fig. 3A**).

Mixtures containing 0.3 mg mLL¹ enzyme and 10× Sypro Orange (Thermo Fisher Scientific) were prepared with a series of buffers (50 mM sodium citrate, 0.1 M NaCl, 5 mM β-ME, pH 4–5.5; and 50 mM sodium phosphate, 0.1 M NaCl, 5 mM β-ME, pH 6–8) to a final volume of 25 μL per well in the 8 strip real-time PCR tubes (BBI Life Sciences, Shanghai, China). The ThermoFluor Assay was conducted in a Real-Time PCR Detection System (Thermo Fisher Scientific, QuantStudio™ 5) with the temperature increased from 25 to 99 °C. The melting temperature (Tm) was calculated by plotting the change of RFU per degree against temperature.

### Substrate specificity profiling of VdiPAL, VyPAL2-I244V and OaAEP1b-C247A

In vitro ligation assays were performed in 50 μL volumes containing 200 nM enzyme (VdiPAL1, VyPAL2-I244V, or OaAEP1b-C247A), 200 μM donor peptide (KALYINQL), 1 mM nucleophilic peptide (GXKRY, where X represents one of the 20 standard amino acids), and the same reaction buffer as above. Reactions were performed in triplicate at 37 °C for 30 min and quenched by heating at 100 °C for 5 min. Ligation and hydrolysis yields were determined by the proportion of HPLC peak areas corresponding to KALYIN-GXKRY and KALYIN, respectively. Yield percentages were plotted in GraphPad to visualize substrate specificity profiles (**Fig. 3B** and **S8**).

### Preparation of FL-LNPs and ABY-FL-LNPs

The preparation of blank liposomes was carried out using a standard thin-film hydration method. 40 mg DSPE (CAS: 1069-79-0), 5 mg DSPE-PEG2000-Alkyne, and 8 mg cholesterol (CAS: 57-88-5) were dissolved in 20 mL dichloromethane and 10 mL methanol. The mixture was sonicated for 40 min (300 W) followed by rotary evaporation at 35°C to form the lipid film. The lipid film was then hydrated in 10 mL aqueous solution (PBS) and sonicated using an ultrasonic cell disruptor (Weimi, Shanghai, China) on ice bath for total 5 min (100 W, 3 s on/10 s off). The particle size of the liposomes was measured using a particle size analyzer (Brookhaven instruments, NanoBrook Omni, New York, USA) (**Fig. S8**). The morphology of the liposomes was observed using transmission electron microscope (Hitachi, Japan, **Fig. S9**). To prepare ABY-FL conjugate, affibody-NQL-His6 used for EGFR-targeting was recombinantly expressed and purified on AKTA Start^TM^. Peptide GISK(5-FAM)K(N_3_) was chemically synthesized (SYNPEPTIDE, Nanjing, China). VdiPAL1-mediated ligation was carried out with 200 μM ABY, 400 μM FL-peptide, 400 nM VdiPAL1 in a pH 6.5 buffer (20 mM sodium phosphate, 1 mM EDTA, 5 mM β-ME) at 37 °C with 10 min, 30 min, and 60 min.

The blank liposomes were then reacted with either the ligation product ABY-FL or the FL peptide to prepare ABY-FL-LNP and FL-LNP, respectively, through click chemistry^53^. Specifically, 100 μM liposomes (equivalent to 0.5 mM alkyne), 2.2 mM CuSOL solution, 50.5 mM sodium ascorbate solution, 4.5 mM 1,10-phenanthroline disodium salt, and 204.4 μM ABY-FL or FL solution were added sequentially. The reaction was conducted at room temperature under N_2_ protection for 6 h. The morphology of both LNPs were examined using TEM (**Fig. S9**).

### Validation of the targeting function of modified liposomes to cells

EGFR-overexpressing A549 cells were treated with either non-targeted fluorescent liposomes (FL-LNP) or affibody-functionalized fluorescent liposomes (ABY-FL-LNP) at a final concentration of 30 μM and incubated for 2 h under standard culture conditions. Following incubation, cells were washed three times with PBS and stained with membrane dye DiI (Beyotime, Shanghai, China) at a final concentration of 5 μM in PBS by incubating at 37 °C for 15 min in the dark. After three additional PBS washes, cells were fixed with 4% paraformaldehyde for 20 min at room temperature. DAPI staining was then performed for 15 min at 37 °C in the dark to visualize nuclei. After a final PBS wash, an anti-fade mounting medium (Beyotime) was applied. Confocal imaging was performed using a Zeiss LSM800 laser scanning confocal microscope equipped with a 63× oil-immersion objective to assess cellular uptake and subcellular localization of the liposomes.

### Protein crystallization and structural determination

Initial crystal screening for VdiPAL1 was performed manually using an Eppendorf electronic dispenser in 96-well sitting drop plates (FAstal BioTech, Shanghai, China). Equal volumes of protein (17 mg mL^-1^) and reservoir solutions were mixed and incubated at 16 °C. After 48 h, rectangular crystals suitable for structural determination were observed in 0.2 M Ammonium sulfate, 0.1 M HEPES (pH 7.7), and 27% w/v polyethylene glycol 3350. For X-ray data collection, crystals were immersed in a cryoprotectant buffer containing 35% PEG 3350 and flash-frozen in liquid nitrogen. X-ray diffraction data were collected on Beamlines BL02U1 of the Shanghai Synchrotron Radiation Facility^54^ (Shanghai Institute of Applied Physics, China) under cryo-conditions (100 K). Data were processed using iMosflm. Phasing was performed by molecular replacement in PHASER-MR^55^, using the AlphaFold3-simulated VdiPAL1 structure as the starting model. Initial automated model building was performed using the Molecular Replacement in PHENIX ^56^, followed by manual model building in COOT^57^. Structure refinement was performed using PHENIX, followed by manual adjustment in COOT (**Table S3**). The final protein structure was resolved as a dimer and deposited in the Protein Data Bank with the accession code 9VSX.

### Complex structure prediction

To investigate the binding mode of VdiPAL1 with its substrate and provide a structural basis for exploring the effect of NAC on the optimal reaction pH, the VdiPAL1-GL12 complex structure was predicted using AlphaFold3 server (https://alphafoldserver.com). Chemically reasonable complex filtered by structural inspection was used in following study.

### System preparation for GROMACS simulation

Simulations were performed at pH values of 4.0, 6.5 and 8.0. The protein and peptide were parameterized using the modified CHARMM36m force field (https://gitlab.com/gromacs-constantph/force-fields). By specifying the pH parameter, the subtool *gentopol* within phbuilder (version 1.3, https://gitlab.com/gromacs-constantph/phbuilder) determined the appropriate initial lambda values automatically. The topological structure and coordinates of the protein-substrate complex were prepared with *editconf* and *solvate* modules in the GROMACS constant pH MD beta version^41,42^ (https://gitlab.com/gromacs-constantph/constantph). To achieve a net-neutral simulation system, the *neutralize* subtool in phbuilder introduced ions to establish charge neutrality at t = 0, and incorporated buffer particles to maintain this neutrality throughout the simulation when t > 0. NaL and ClL were used as counterions.

### Molecular mechanics (MM) minimization

To minimize the potential energy of the system with correct parameters, the subtool *genparams* embedded in phbuilder generated “EM.mdp” file according to the specified pH value. The real space non-bonded cutoff, encompassing van der Waals and short-range electrostatic interactions, was set to 12 Å in accordance with the CHARMM force field parameters. Long-range electrostatic interactions were calculated using the Particle Mesh Ewald (PME) algorithm. Steepest descent minimization, with a maximum of 5000 steps, was conducted using *grompp* and *mdrun* modules in the GROMACS constant pH MD beta version.

### Constant pH molecular dynamics (cpHMD) simulation

In addition to the energy minimization phase, the subtool *genparams* also generated appropriate cpHMD parameter files for both the equilibration and production MD phases. To be consistent with the experimental conditions, a constant temperature was maintained at 310.15 K using the V-rescale thermostat with a coupling time constant of 0.5 ps. Pressure was stabilized at 1 bar using the C-rescale barostat with a coupling time constant of 5.0 ps. During the MD simulation, the LINCS algorithm was applied to constrain the covalent bonds involving hydrogen atoms, thereby enabling the use of a 2 fs time step. Equilibration was performed for 10 ps in the NVT ensemble, followed by 10 ps of equilibration in the NPT ensemble. Subsequently, cpHMD simulations were conducted under the NPT ensemble using the GROMACS constant pH MD beta version. Two sets of simulations were performed with different initial velocities: fifteen independent 100 ns simulations and three independent 1 μs simulations. Atomic coordinates were recorded every 50 ps, yielding 2,000 snapshots per 100 ns trajectory and 20,000 snapshots per 1 μs trajectory^42^.

### Near Attack Conformation (NAC) analysis

To evaluate the feasibility of amide bond formation at different pH levels, the percentage of frames meeting the NAC criteria was calculated. The distance between the sulfur atom of VdiPAL1-Cys213 and the carbonyl carbon of GL12-Asn10 was measured. Additionally, the angle formed by Cys213 (S)/His171 (ND1) of VdiPAL1, carbonyl carbon (C) and carbonyl oxygen (O) of GL12-Asn10 was also determined. Given that the Bondi radii of carbon, nitrogen and sulfur are typically 1.70 Å, 1.55 Å and 1.80 Å, respectively, and the attack-ready Bürgi-Dunitz angle is 90 degrees, the NAC condition was defined as a distance of less than 3.5 Å and an angle ranging from 80 to 100 degrees (90 ± 10 degrees). Interatomic distances and bond angles were calculated using the MDAnalysis toolkit based on the atomic coordinates extracted from the trajectories.

### Classical molecular dynamics simulation

The initial structures of VdiPAL1-C213A and H171A in complex with the GL12 peptide were predicted by AlphaFold3, solvated in a rectangular TIP3P water box with a minimum solute-box distance of 1.5 nm, and neutralized with NaL/ClL ions. All simulations were performed using GROMACS 2023 with the AMBER14SB force field. Following a 50,000-step steepest descent energy minimization, each system was equilibrated for 100 ps in the NPT ensemble at 310.15 K and 1.0 bar, applying positional restraints to the complex’s heavy atoms. This stage utilized a V-rescale thermostat and a Berendsen barostat. Finally, 100 ns production simulations were conducted in the NPT ensemble, switching to the Parrinello-Rahman barostat. The simulations employed a 2 fs timestep, with hydrogen bonds constrained by the LINCS algorithm and long-range electrostatics treated with the Particle Mesh Ewald (PME) method.

### Binding free energy (ΔG_bind) calculation

To quantify the binding affinity between the protein and the peptide, the binding free energy (ΔG_bind) was calculated using the Molecular Mechanics/Poisson-Boltzmann Surface Area (MM/PBSA) method. All calculations were performed using the gmx_MMPBSA tool^58^. The solvation free energy (G_solv) was decomposed into polar and non-polar components. The polar solvation energy (G_polar) was obtained by solving the Poisson-Boltzmann (PB) equation, with the ion concentration of 0.15 M to simulate physiological salt conditions. The non-polar solvation energy (G_nonpolar) was estimated using the solvent accessible surface area (SASA) model (γ = 0.0072 kcal·molL¹·ÅL², β = 0.00 kcal·molL¹).

### Mutagenesis and activity study of VdiPAL1 mutants

Alanine substitution of Q342, D262, S241, E211, C213, H171, and N66 were carried out with mutagenesis kit C214 (Vazyme, Nanjing, China). Mutants were expressed in 24 well plates. The fermentation products were lysed using the same method described above to obtain crude enzyme supernatants. Western Blot analysis was performed to detect expression levels. For the activity screening assay, 10 μL of each mutant’s crude supernatant was incubated with 20 μM substrate GL-12 in citrate buffer (20 mM sodium citrate, 1 mM EDTA, 5 mM β-ME, pH 5.0) at 37 °C for 30 min. The cyclized product and substrate were monitored by HPLC. H171A and C213A were scaled up to 2 L fermentation cultures, purified via Ni-column chromatography, and subjected to acidification using the same method described above. One-pot autoactivation and cyclization reactions were conducted with 10 μM purified proenzyme and 20 μM GL12 in citrate buffer (20 mM sodium citrate, 1 mM EDTA, 5 mM β-ME, pH 4.0) at 30 °C overnight. Upon completion, the reaction mixtures were analyzed by HPLC to determine the cyclized product and substrate.

### Design of combinatorial mutation for VyPAL2-I244V

Top seven expression-enhancing single-point mutation of VyPAL2-I244V (I351L, N356Q, M357N, N397D, V409A, A422T, Q454R) were selected for combinatorial design. To avoid the potential instability of ΔΔG prediction with excessively high mutational loads, we limited the number of simultaneous mutations to 2-5 per design. The input structural model of VyPAL2 was generated using AlphaFold3 (RMSD < 0.25 Å compared with crystal structure 6IDV and 7F5J). Targeted sampling of specified mutation sites was implemented using the circular design script submit_example_4_non_fixed.sh from the official ProteinMPNN repository (https://github.com/dauparas/ProteinMPNN). This allowed mutational variation exclusively at selected positions, while keeping the remaining sequence fixed. For each site combination, up to 100 cycles of sequence generation were performed. The process was terminated early under any of the following criteria: (1) no new valid sequences were added for 5 consecutive cycles, (2) the candidate list reached 200 sequences, or (3) the maximum of 100 cycles was completed. Candidate sequences were retained if >40% of their mutated residues overlapped with the known expression-enhancing positions. In total, 265 combinatorial mutant designs were selected for downstream ΔΔG filtering and experimental testing.

### Free energy change (ΔΔG) prediction

To further enrich for high-confidence variants, we applied a structural stability filter based on predicted changes in folding free energy (ΔΔG) upon mutation. For each of the 265 combinatorial mutant designs, Rosetta ΔΔG predictions were performed using the cartesian_ddg protocol (Rosetta v3.13) with the ref2015 scoring function. Each mutant was evaluated over five independent replicates, and average ΔΔG values were computed. All calculations were automated using the Python wrapper RosettaDDGPrediction^59^. Designs with >40% overlap with known expression-enhancing positions and a predicted ΔΔG ≤ –1.5 Rosetta Energy Units (REU) were retained. This yielded 25 candidate variants for experimental validation.

## Acknowledgement

We thank the botanist Ce-hong Li (Emei Mountain Biotic Resource Experimental Station of China) for aiding on field collection of *Viola* plants, Julien Lescar (Nanyang Institute of Structural Biology, Singapore) for advice on crystallography data analysis, and the staff of beamline BL02U1 at the Shanghai Synchrotron Radiation Facility for assistance during the data collection. This work was supported by the National Key Research and Development Program of China (2023YFA0916000), National Natural Science Foundation of China (32371324), the High-level Talent Startup Fund by China Pharmaceutical University, the Key R&D Program of Hubei Jiangxia Laboratory (JXBS010), and the National Natural Science Foundation of China (32270228).

## Author contributions

W.D.: conceptualization, methodology, wild-type and mutant library construction, data analysis, activity assays, crystal structure solving and refinement, writing-original draft, writing-editing; S.Q.: conceptualization, datamining, modeling and simulation, computer-assisted engineering, writing-original draft; M.Z.: methodology, LNP preparation, functionalization and examination. M.L., H.J., and G.L.: resources, field collection. Y.Z. and Y.H: mutant library construction and screening; J.M.: protein purification and crystallization; C.W.L. and N.C, crystal structure solving; C.C., methodology, LNP preparation; G-W.H:, resources, writing-editing, funding; X.H: conceptualization, methodology, writing-editing, supervision, funding.

## Data Availability

Authors declare that all data supporting the findings of this study are available within the paper and its supplementary information files. HPLC and LC-MS data files are available at Figshare (https://doi.org/10.6084/m9.figshare.30170467).

