## supporting information 1 for "Mining of natural diversity enables efficient and expressible peptide asparaginyl ligases"

Table of Contents

Supplementary Figure S1-S46

Supplementary Tables S1-S4

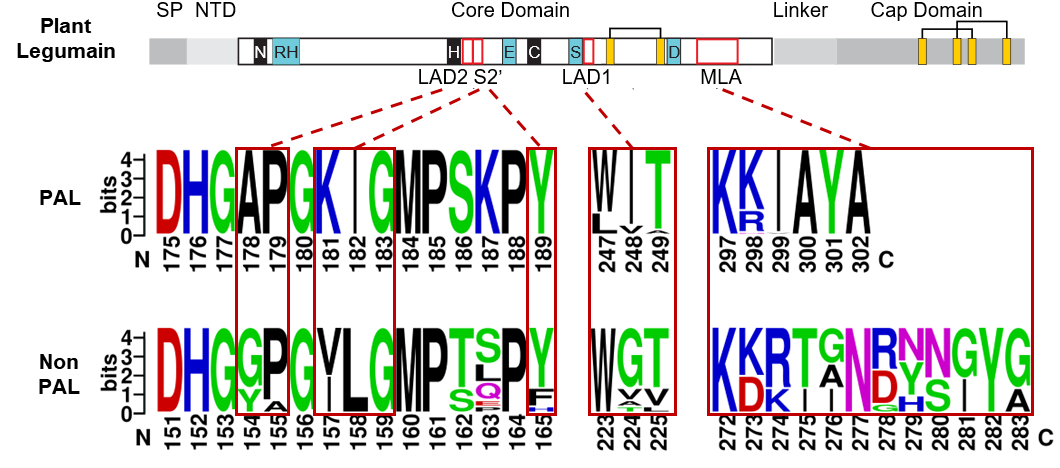

**Figure S1 Sequence logos for the highly conserved motifs of *Viola* legumain.** The top panel shows a schematic of the protein structure with the relative positions of conserved motifs indicated by red boxes. The following sequence logos compare the LAD2, S2', LAD1, and MLA motifs between PAL and non-PAL proteins. The size of each character indicates its occurrence frequency within the respective protein family.

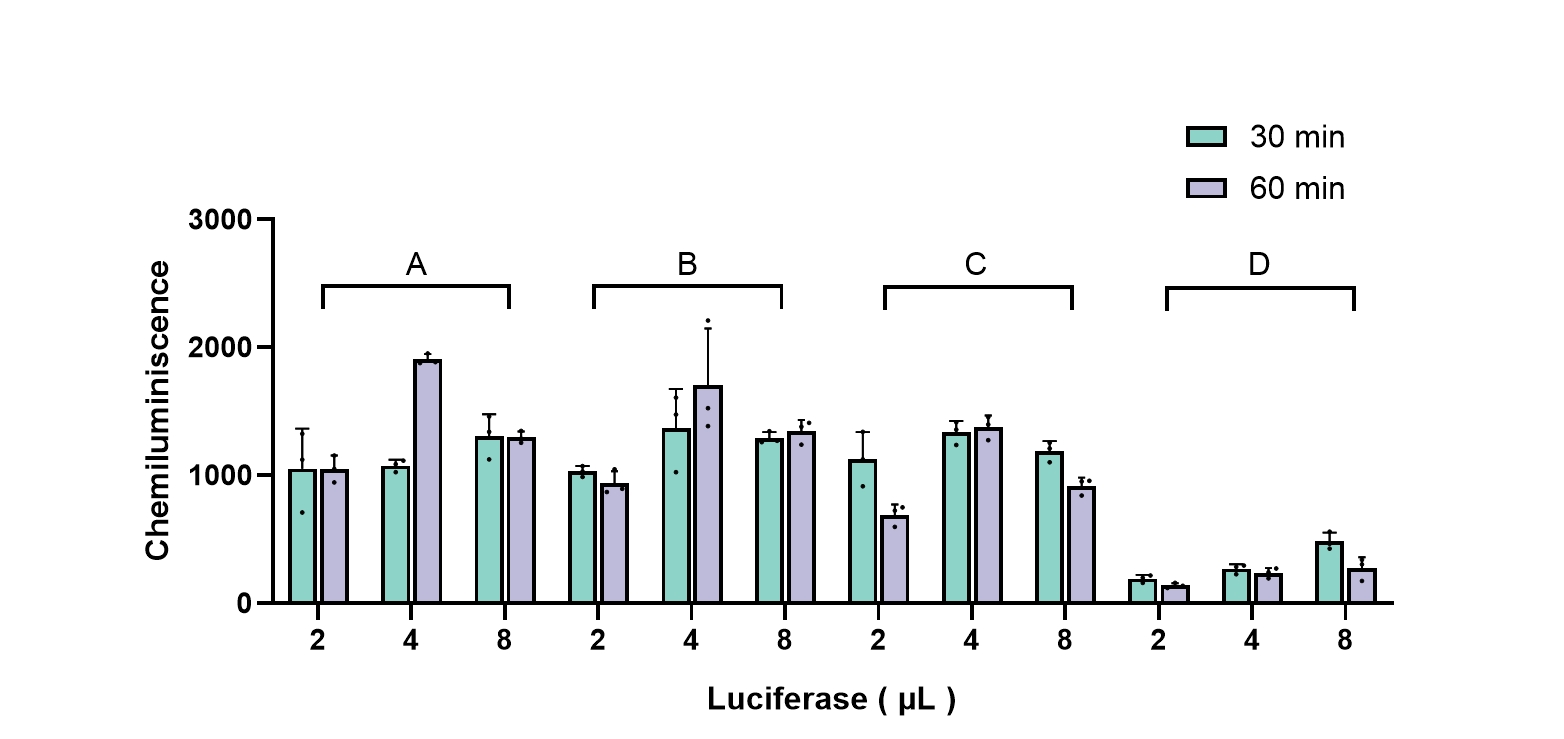
Figure S2 Effect of linkage reaction time and product addition volume on luminescence signal in the first step of the NanoLuc screening methodology using OaAEP1b-C247A. The green and purple colors in the figure represent ligation reactions carried out for 30 and 60 min, the volume of crude enzyme supernatant was set up in four gradients A-D, which were 1, 2, 4, and 8 μL, respectively. The second step of luminescence signal detection was performed by taking 2, 4, and 8 μL of product from each set of linkage reactions, respectively.

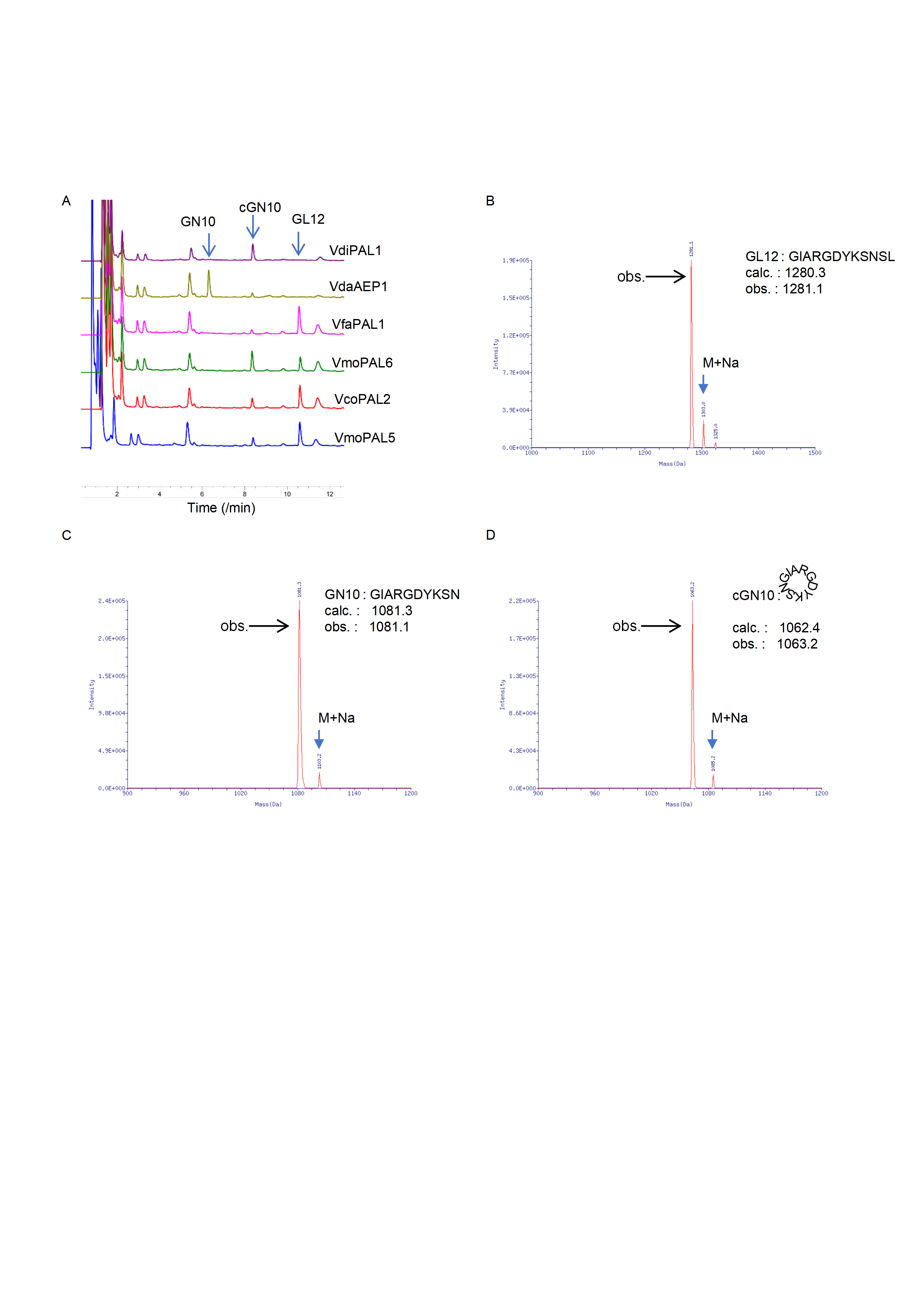

Figure S3 HPLC detection of the cyclization reactions of six candidate enzymes screened by the NanoLuc method. (A) HPLC chromatograms of the cyclization reactions of the six candidate enzymes. The substrates GL12, linear ligation products cGN, and hydrolysis products GN10 are indicated. (B) Mass spectrometry spectrum of the peptide substrate GL12. (C) Mass spectrometry spectrum of the peptide hydrolysis product GN10. (D) Mass spectrometry spectrum of the peptide cyclization product cGN10.

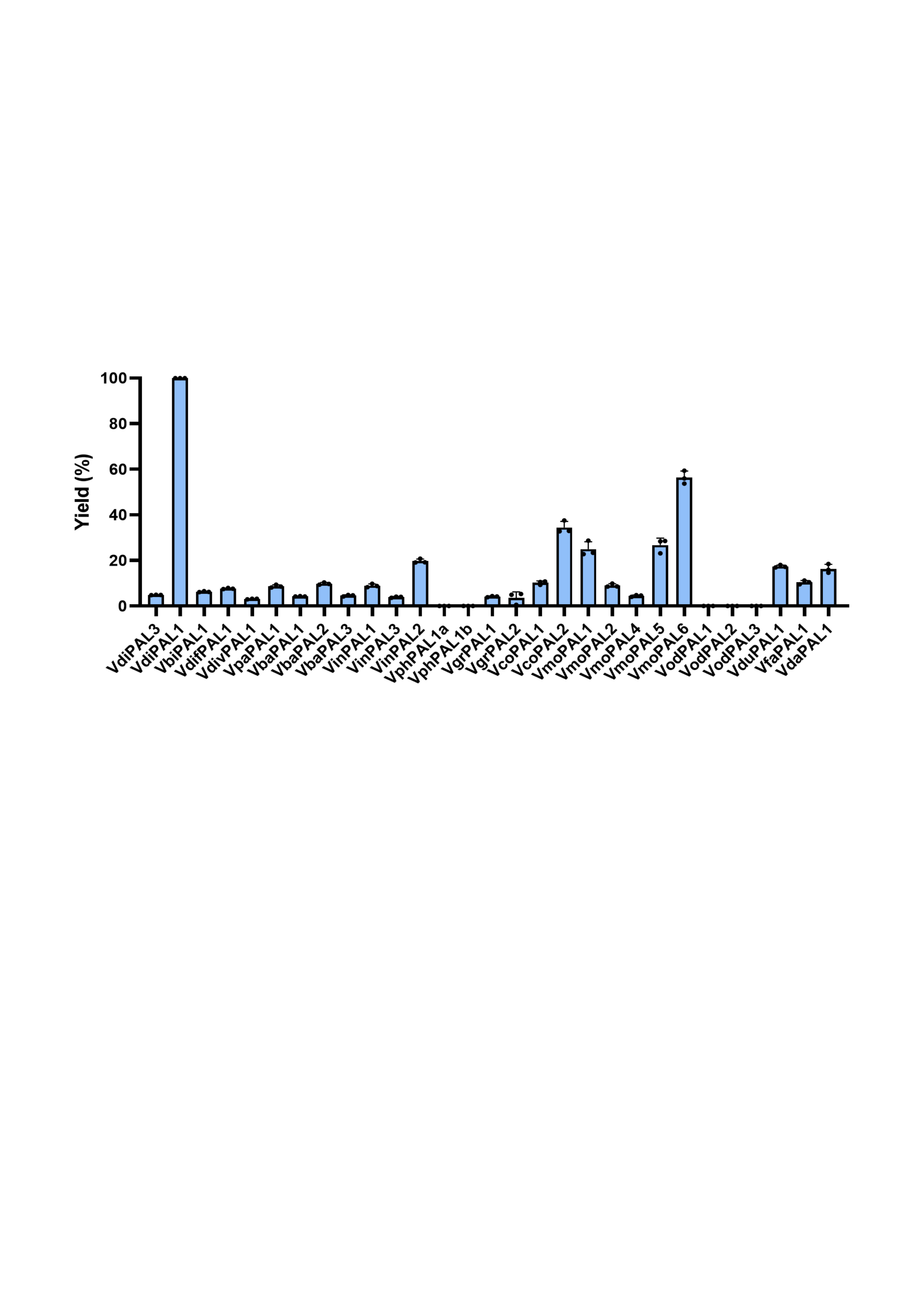

Figure S4 HPLC analysis of cyclization reactions catalyzed by 29 novel PALs. Yield of cGN10 (%C) was in blue. 10 μL of crude enzyme supernatant was added to a reaction buffer (50 mM sodium citrate, 0.1 M NaCl, 5 mM β-ME and pH 5) containing 20 μM substrate peptide GL12. The mixture was incubated at 37 °C for 60 min. The conversion rate of the cyclized product was determined by HPLC. Notably, no hydrolysis products were detected for these enzymes.

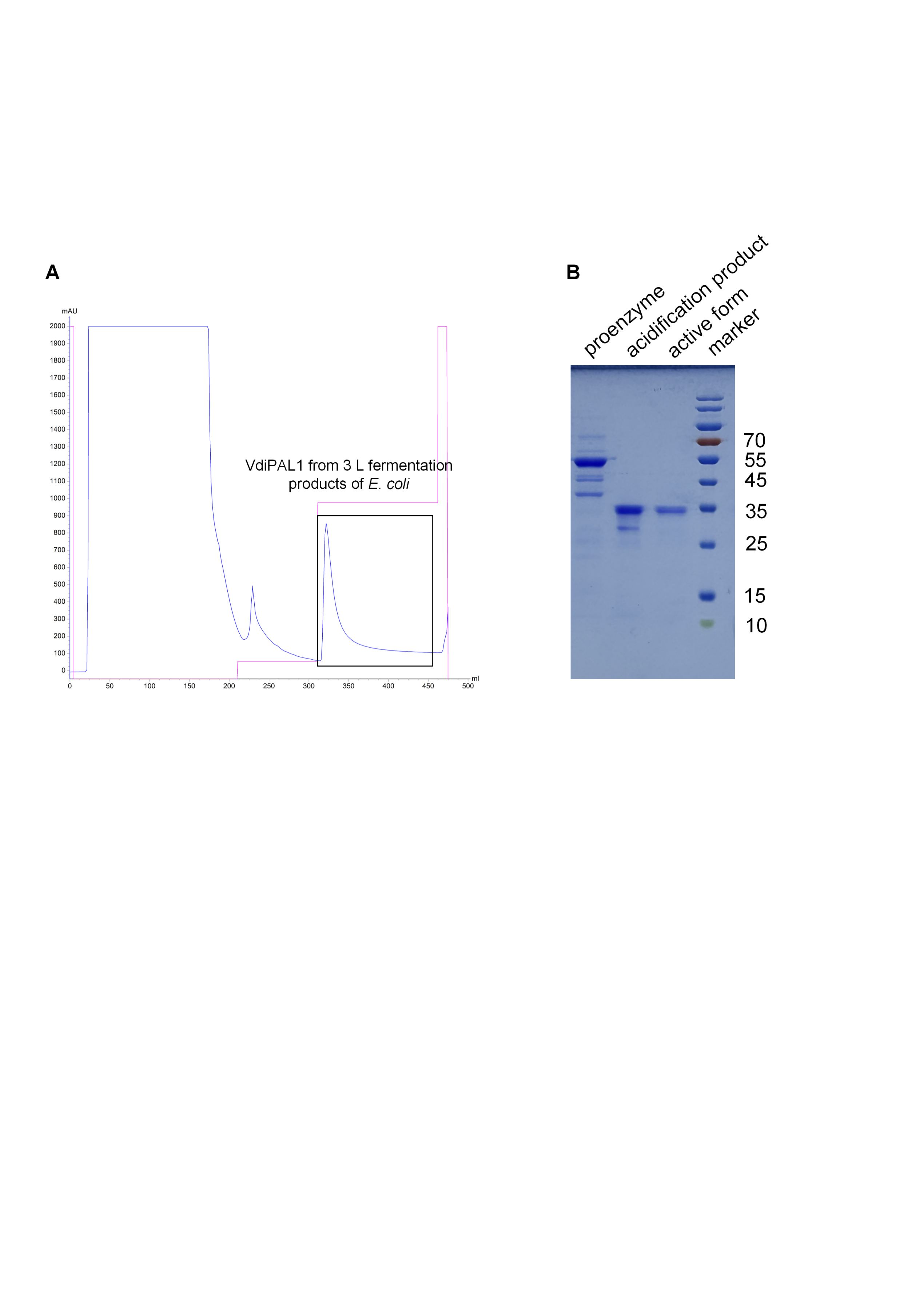

Figure S5 Purification of the VdiPAL1. (A) The blue line represents the UV absorbance at 280 nm, and the red line indicates the imidazole concentration. Fractions containing the target protein, purified from a 3 L *E. coli* fermentation using Ni-affinity chromatography, are enclosed in the black box. (B) The SDS-PAGE of purified VdiPAL1 proenzyme (55 kDa) and active form (35 kDa).

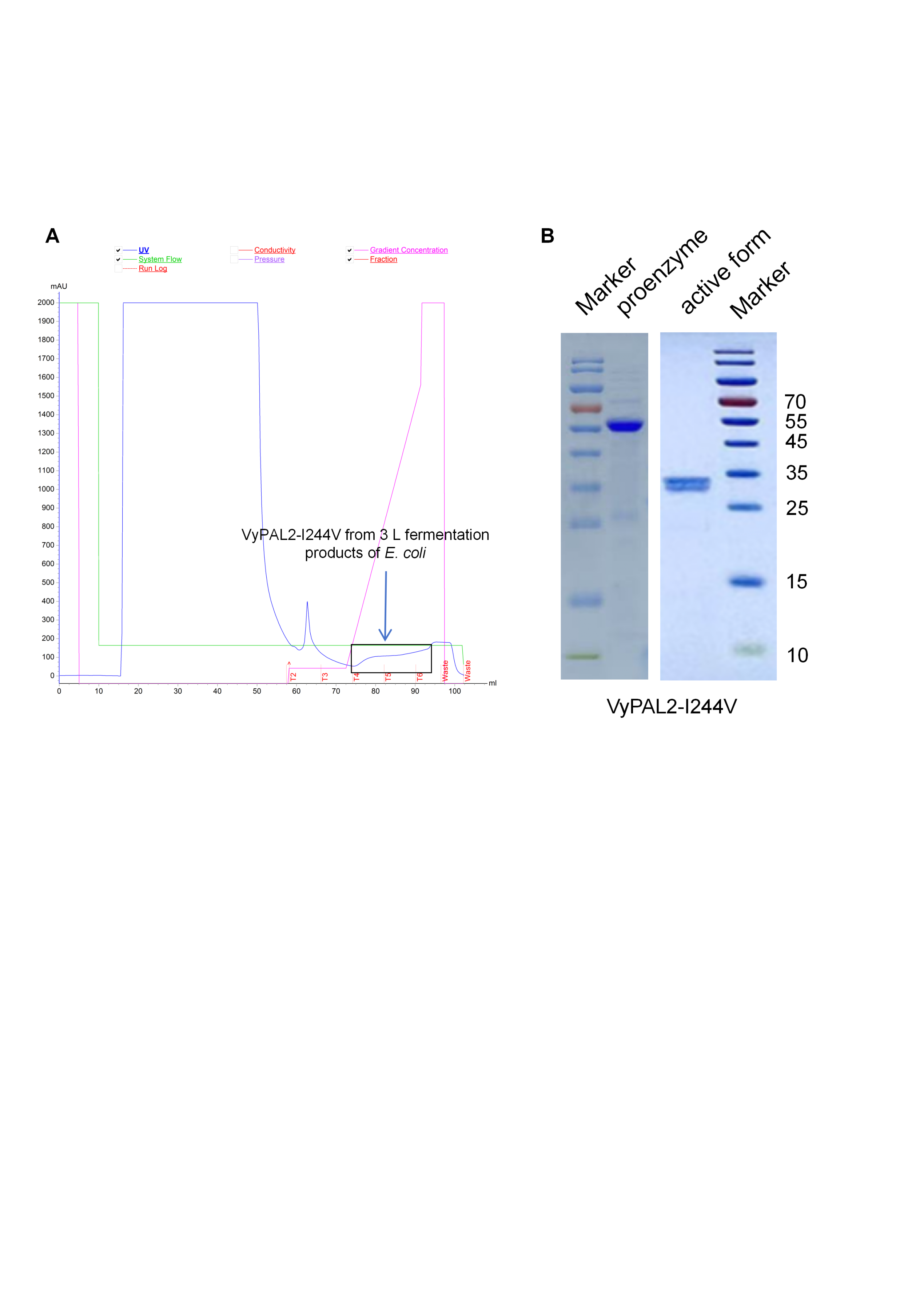

Figure S6 Purification of the VyPAL2-I244V. (A) The blue line represents the UV absorbance at 280 nm, and the red line indicates the imidazole concentration. Fractions containing the target protein, purified from a 3 L *E. coli* fermentation using Ni-affinity chromatography, are enclosed in the black box. (B) The SDS-PAGE of purified VdiPAL1 proenzyme (55 kDa) and active form (32 kDa).

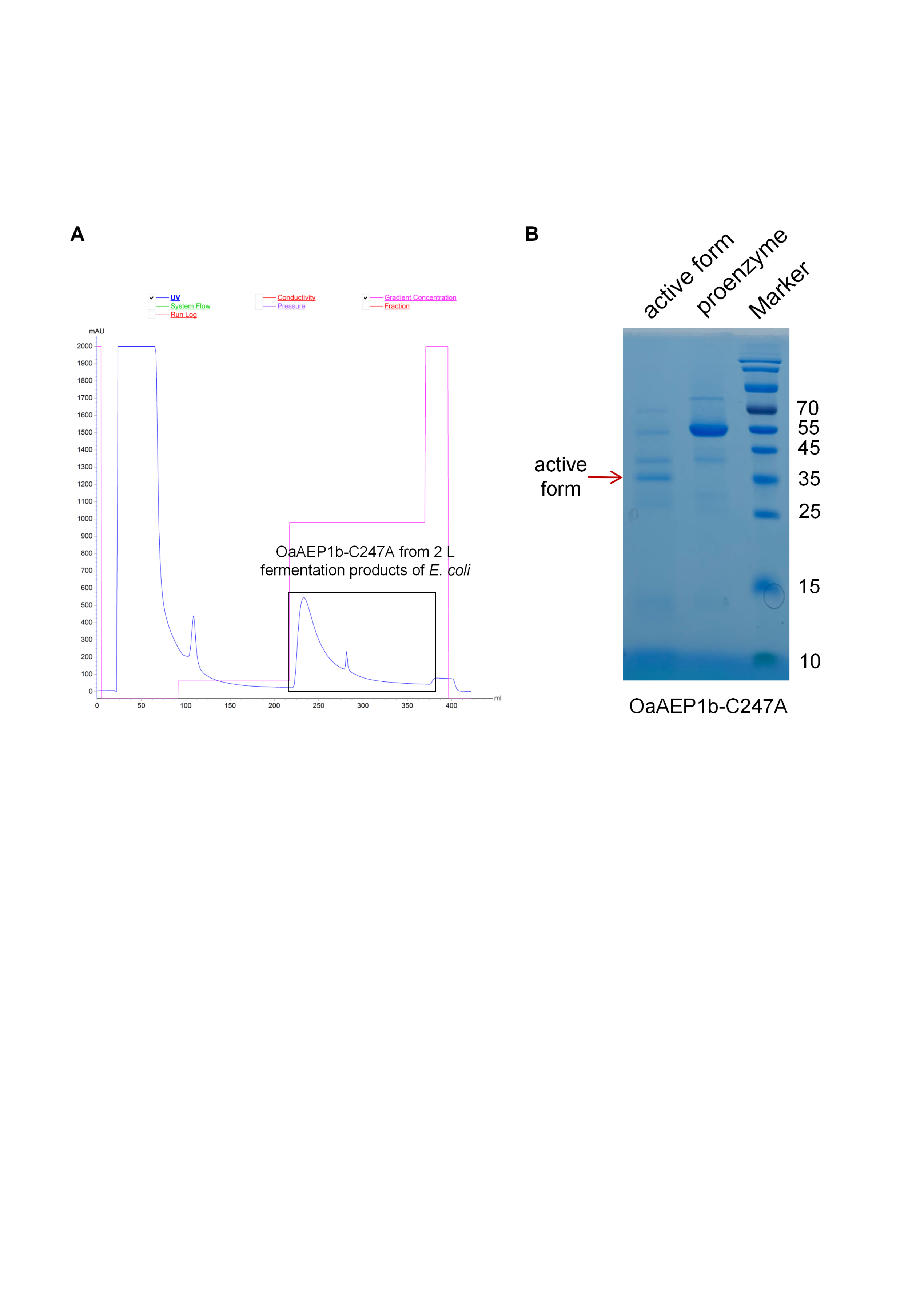

Figure S7 Purification of the OaAEP1b-C247A. (A) The blue line represents the UV absorbance at 280 nm, and the red line indicates the imidazole concentration. Fractions containing the target protein, purified from a 2 L *E. coli* fermentation using Ni-affinity chromatography, are enclosed in the black box. (B) The SDS-PAGE of purified VdiPAL1 proenzyme (55 kDa) and active form (35 kDa).

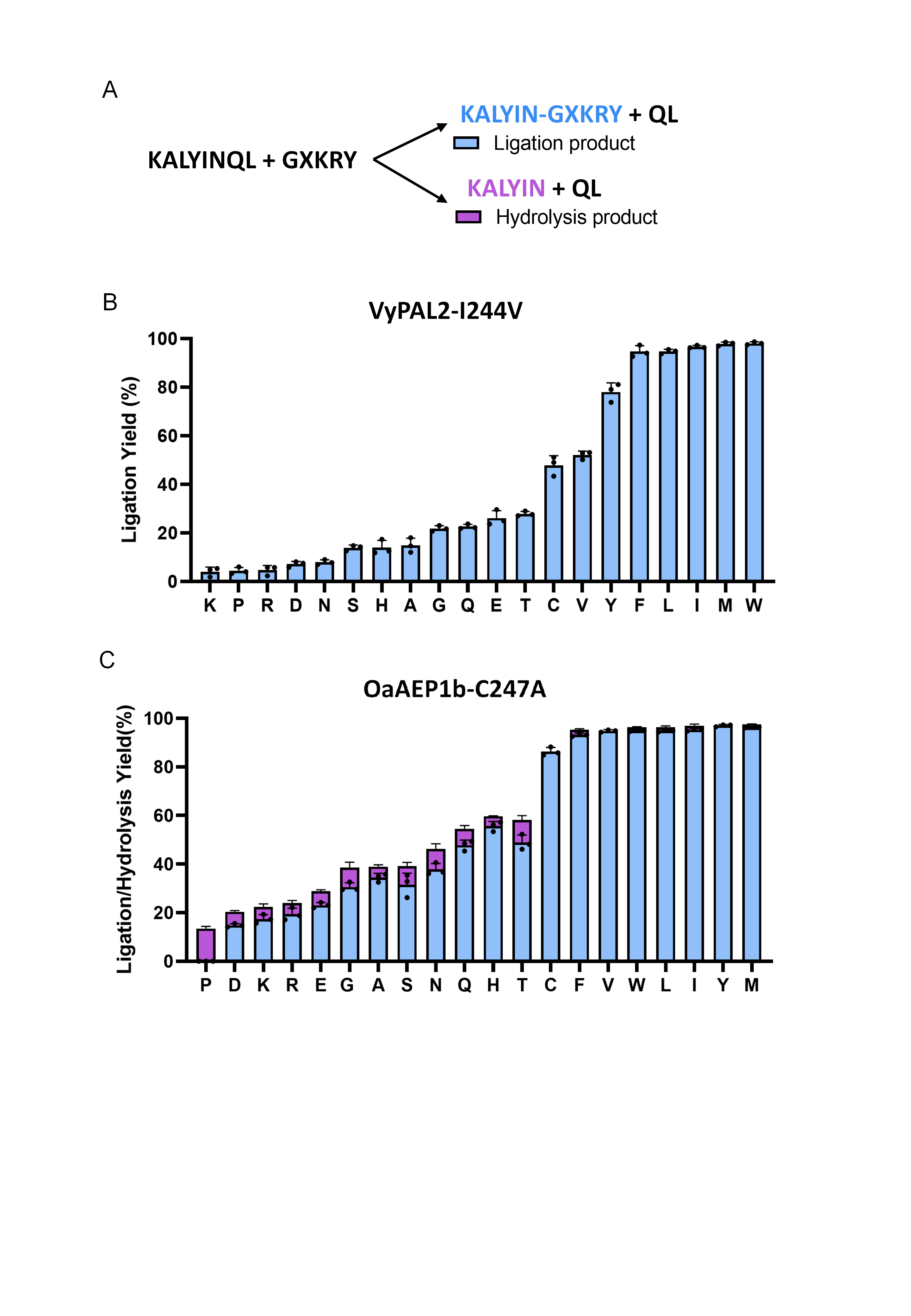

Figure S8 Substrate specificity of VyPAL2-I244V and OaAEP1b-C247A at the P2''. (A) Schematic representation of ligation substrate specificity, ligation and hydrolysis yields are shown for reactions between KALYINQL and GXKRY peptides, where X represents each of the 20 standard amino acids. Ligation products and hydrolysis products are represented by blue and purple bars, respectively. (B) Ligation yields of VyPAL2-I244V with different P2'' substrates. (C) Ligation and hydrolysis yields of OaAEP1b-C247A with different P2'' substrates.

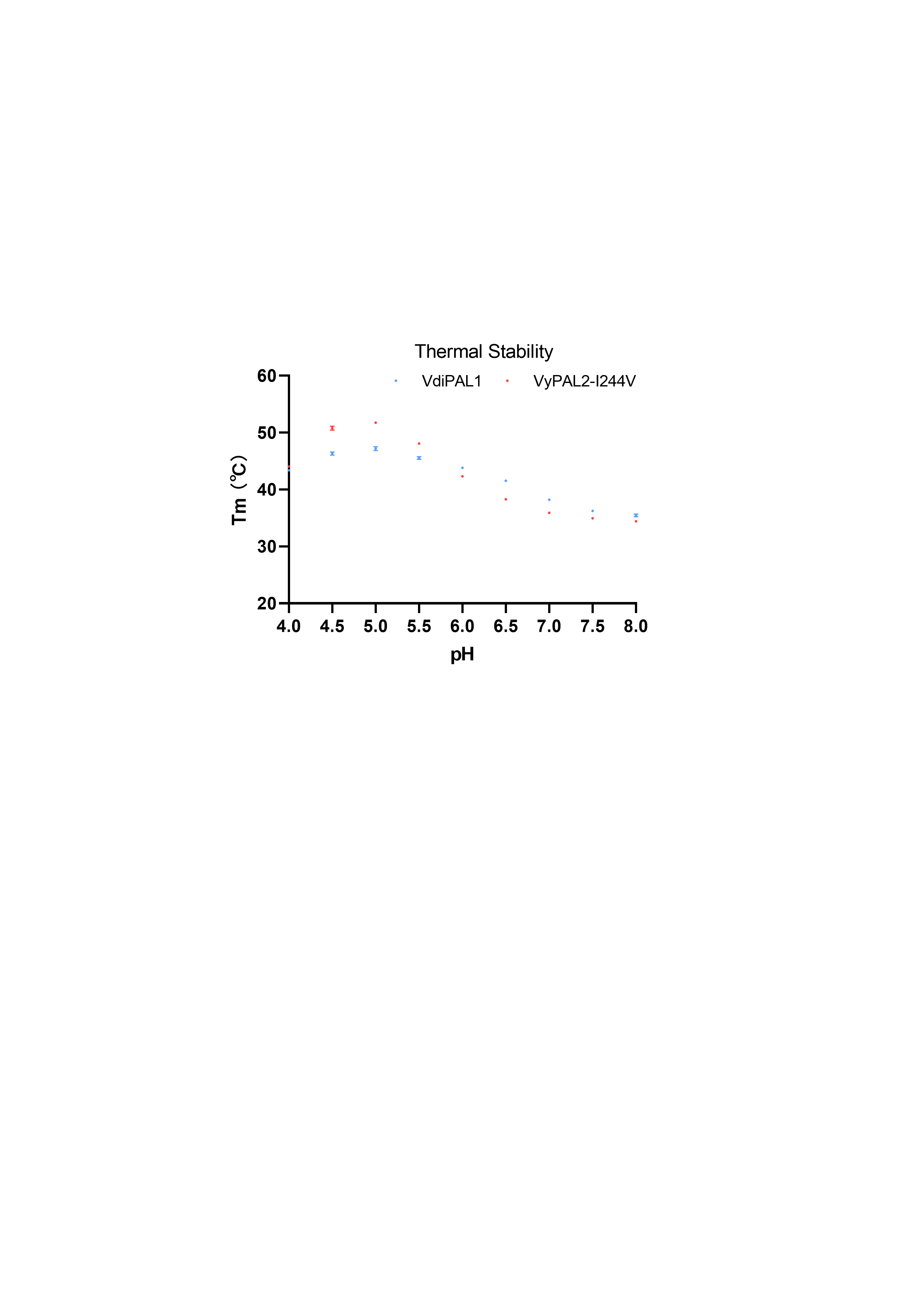

Figure S9 Thermal Stability of VyPAL2 and VdiPAL1. Thermal shift assays were conducted using SYPRO Orange dye with salt-free buffers at a pH ranging from 4 to 8.

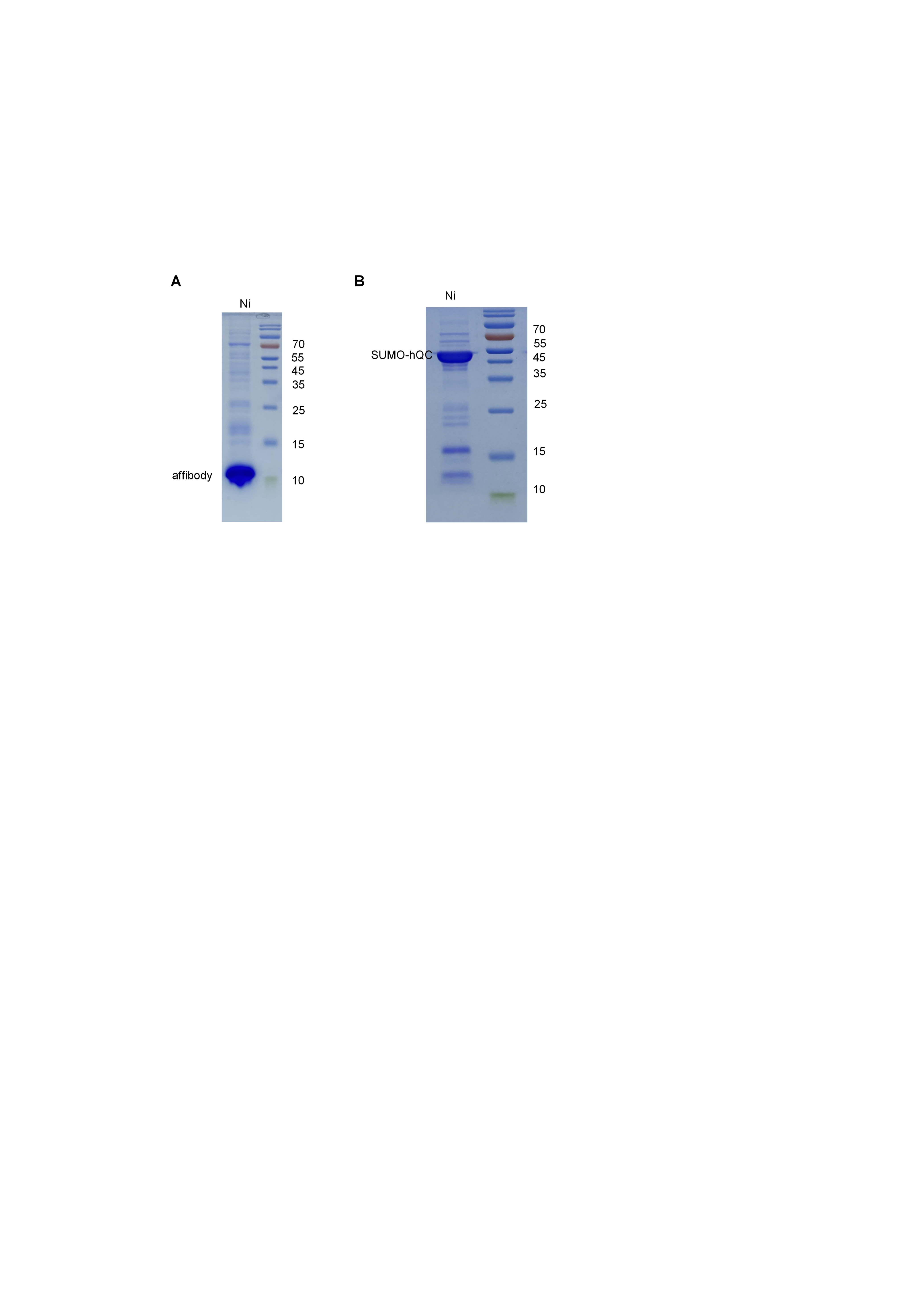

Figure S10 Expression and purification of proteins related to the reactions. (A) Affibody protein purified by Ni-affinity chromatography, with a target molecular weight of 9.7 kDa. (B) Human QC protein with a SUMO tag purified by Ni-affinity chromatography, with a target molecular weight of 50 kDa.

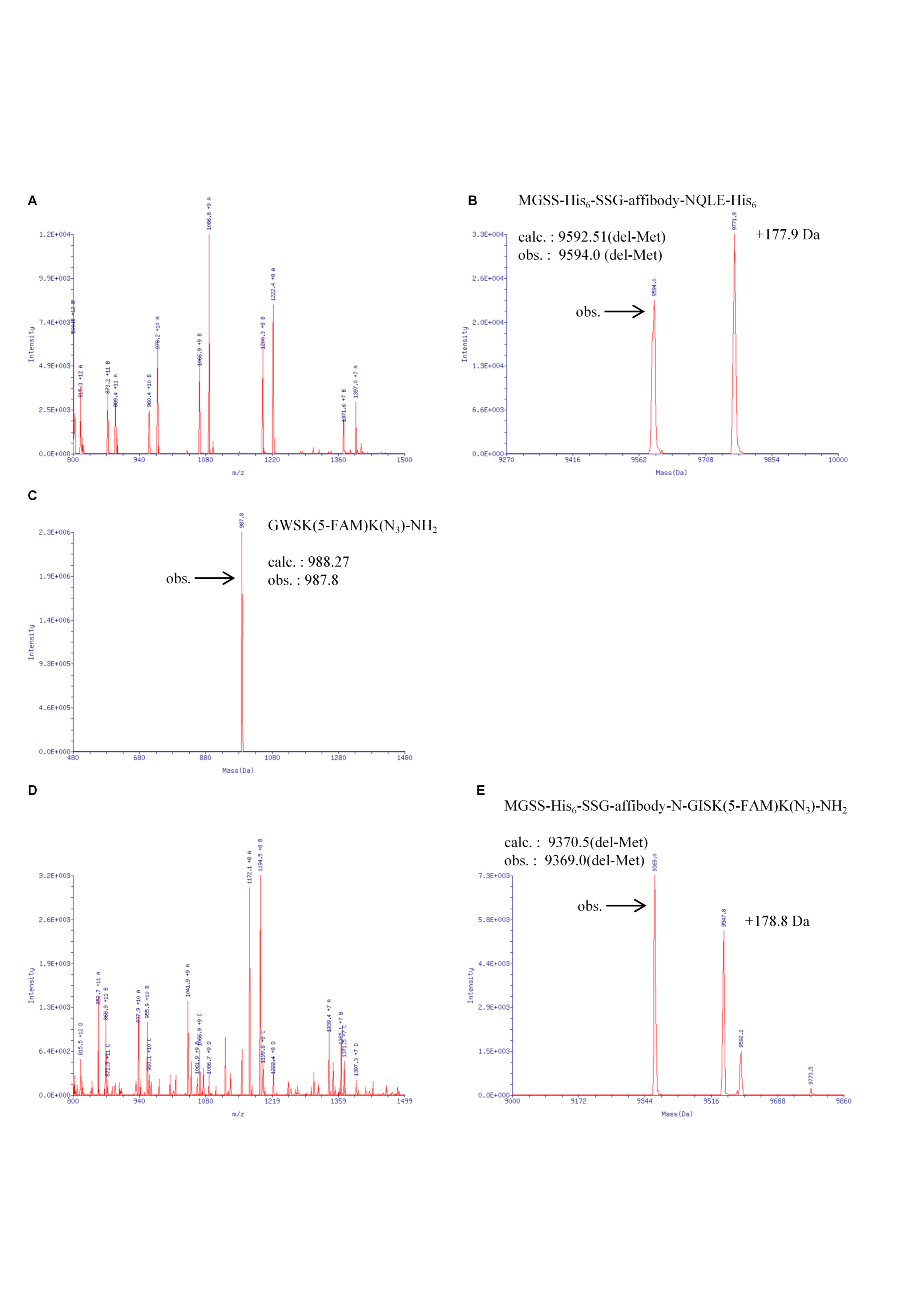

Figure S11 VdiPAL1-mediated ligation of an affibody and a fluorescent peptide. (A) m/z mass spectrum of the Affibody protein. (B) Deconvoluted mass spectrum of the Affibody protein, with the obs. molecular weight of 9594.0 Da (with N-terminal methionine removed). (C) Mass spectrum of the fluorescent peptide, showing the obs. molecular weight of 988.27 Da and an obs. molecular weight of 9987.8 Da. (D) m/z mass spectrum of the conjugation product. (E) Deconvoluted mass spectrum of the conjugation product, with a calc. molecular weight of 9370.5 Da and an obs. molecular weight of 9369,0 Da (accounted for by formic acid adduct formation).

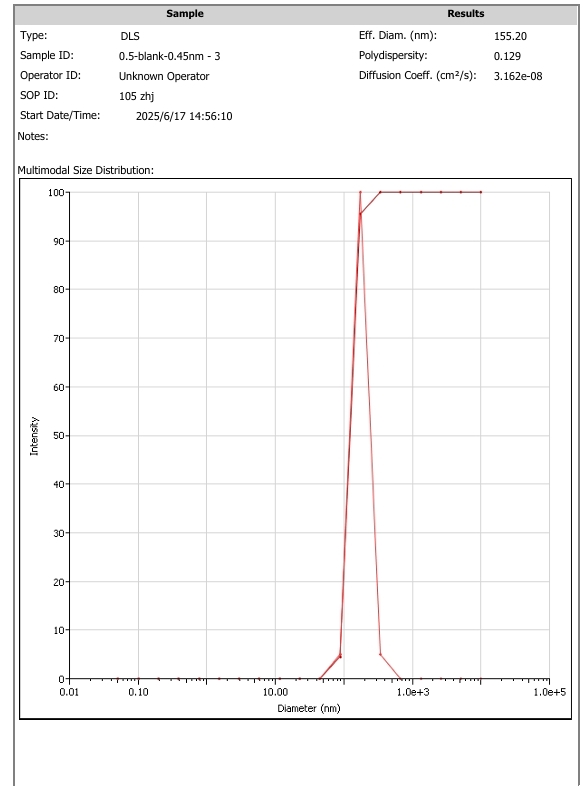

Figure S12 Dynamic Light Scattering (DLS) profiles of liposome samples.

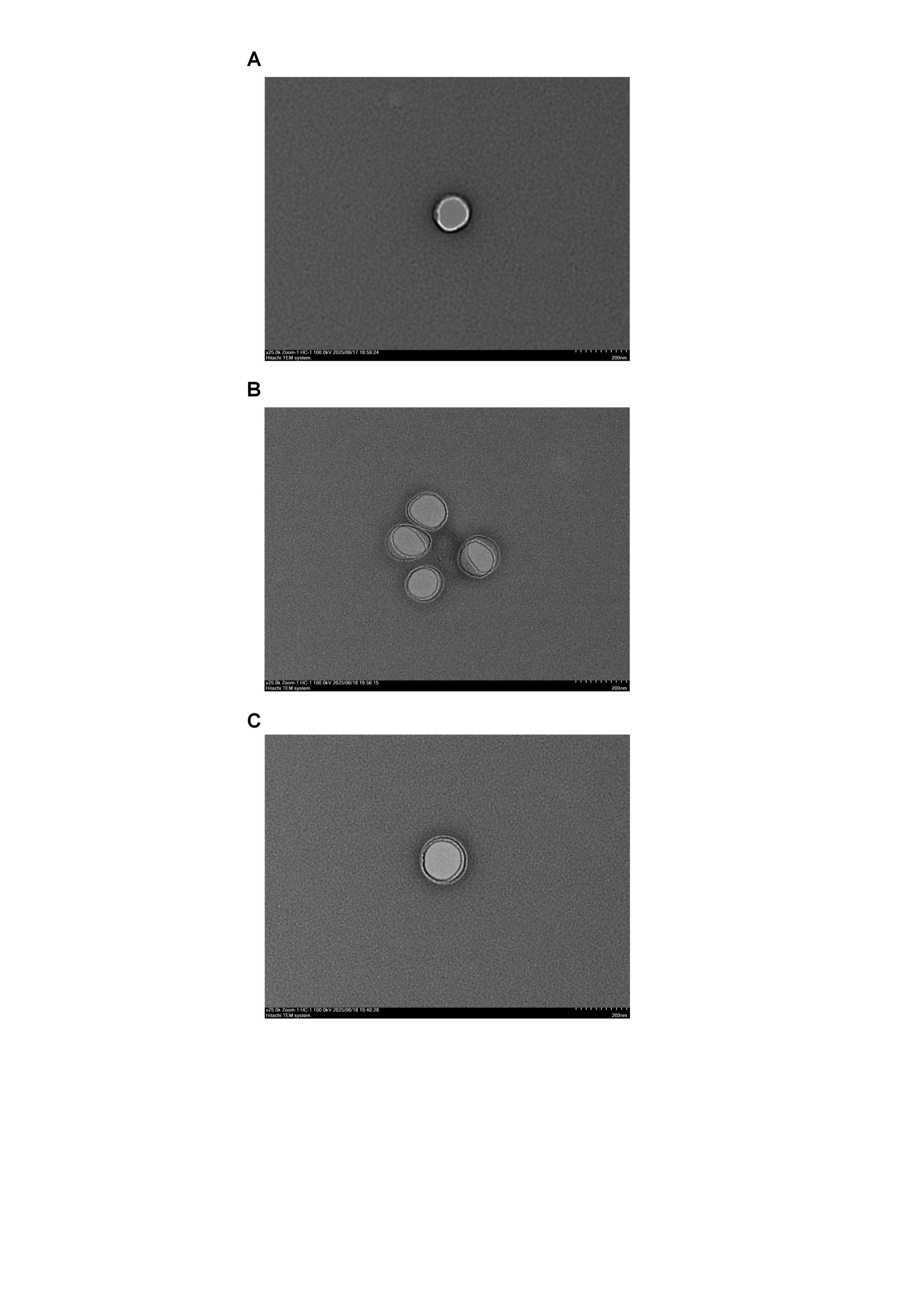

Figure S13 Transmission Electron Microscopy (TEM) images of the three types of liposomes. (A) blank liposomes, (B) FL liposomes, and (C) ABY-FL liposomes.

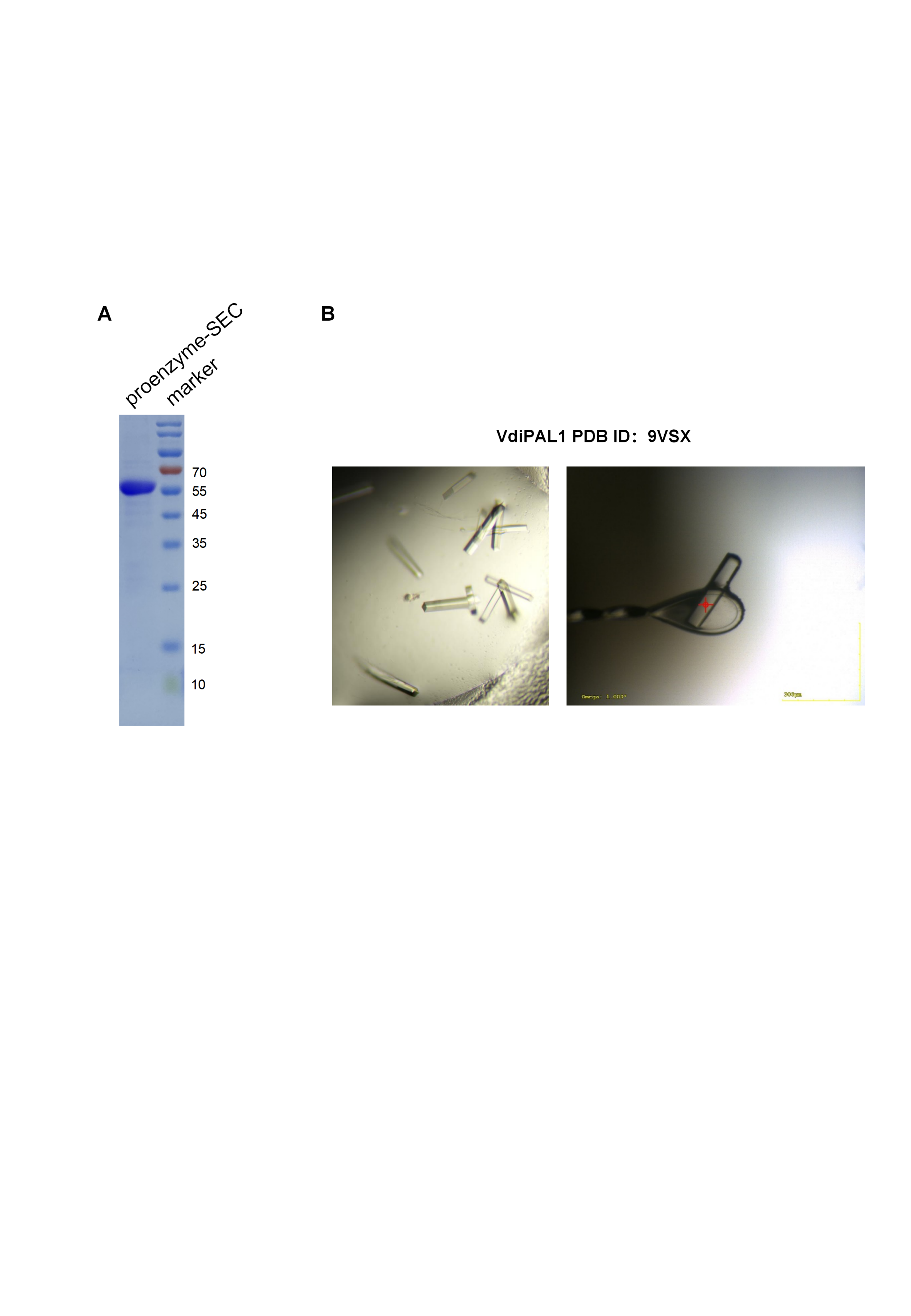

Figure S14 Purification and crystal structure of VdiPAL1. (A) The SDS-PAGE of the purified VdiPAL1 proenzyme using SEC (55 kDa). (C)The crystal morphology of the VdiPAL1 (PDB ID: 9VSX).

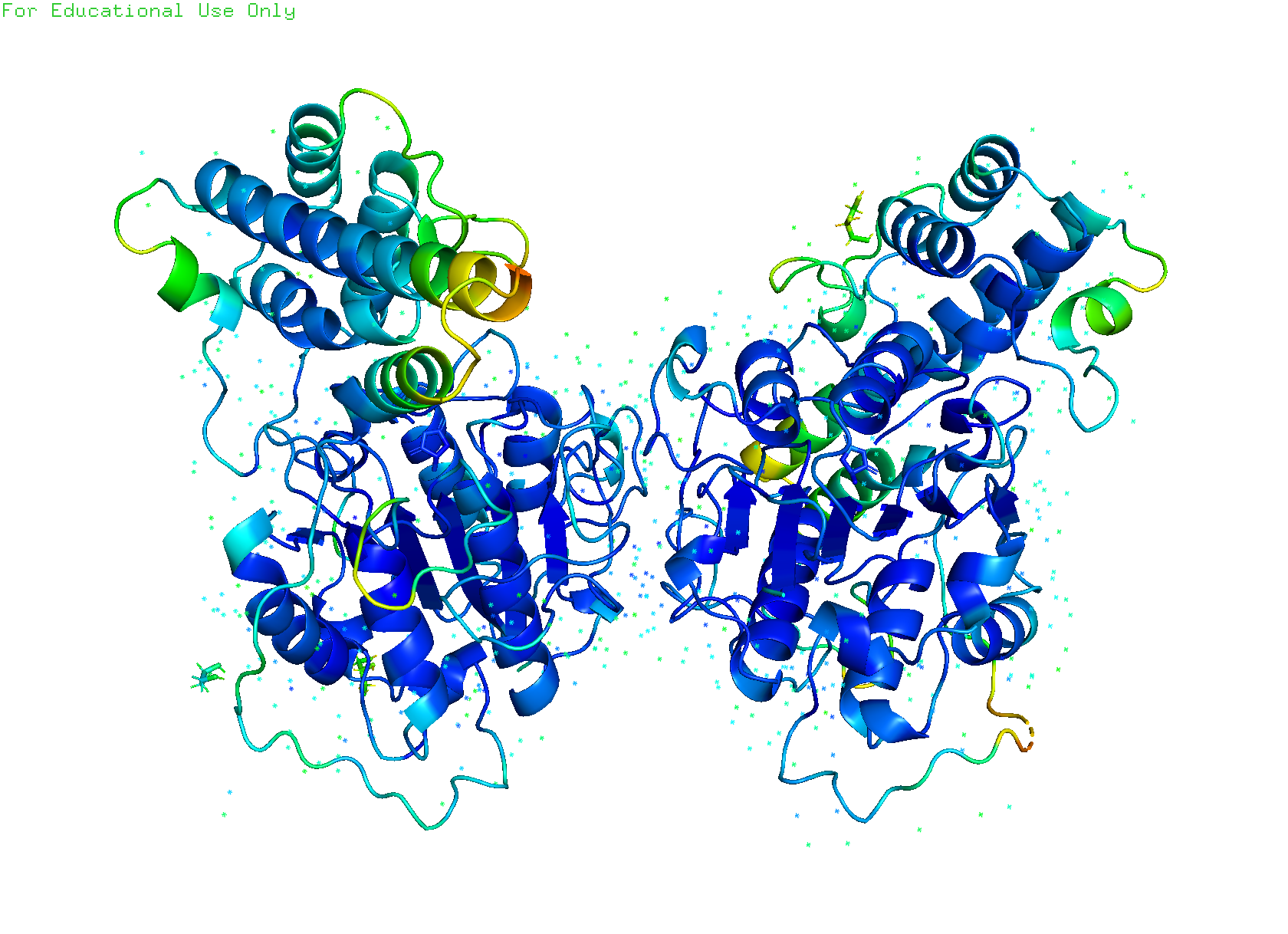

Figure S15 The crystal structure of VdiPAL1. The dimeric VdiPAL1 structure, with a resolution of 1.87 Å, allows for clear observation of the water molecules within the structure (PDB ID: 9VSX).

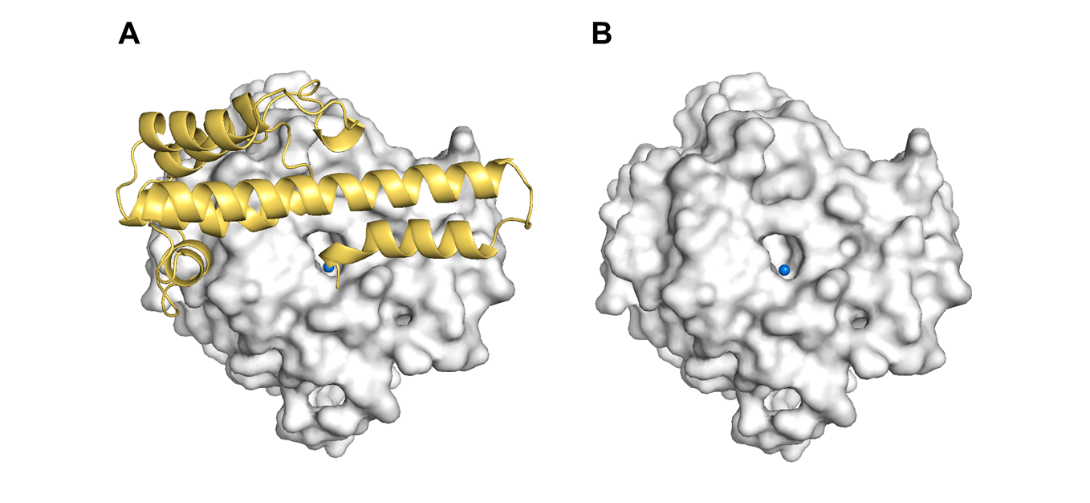

Figure S16 Water molecules in the catalytic hole of VdiPAL1 structure. (A) The VdiPAL1 proenzyme structure, with the cap domain displayed in yellow cartoon and the core domain in gray surface. (B) The core domain clearly revealed the water molecules in the catalytic hole.

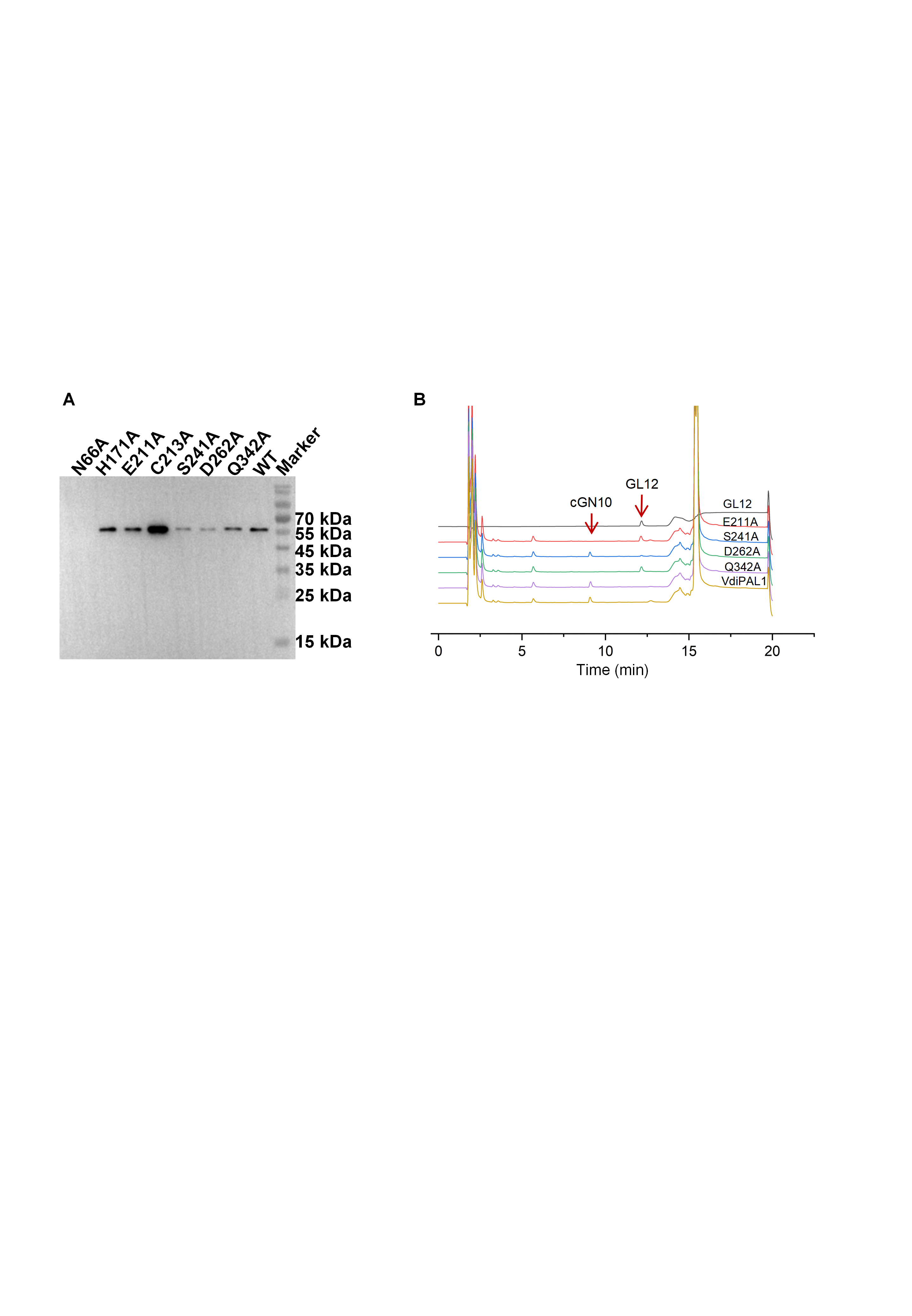

Figure S17 Expression and activity analysis of VdiPAL1 mutants. (A) Western blot analysis showing the expression levels of mutants with alanine substitutions in the catalytic pocket. (B) Cyclization activity of partial alanine mutants determined by HPLC.

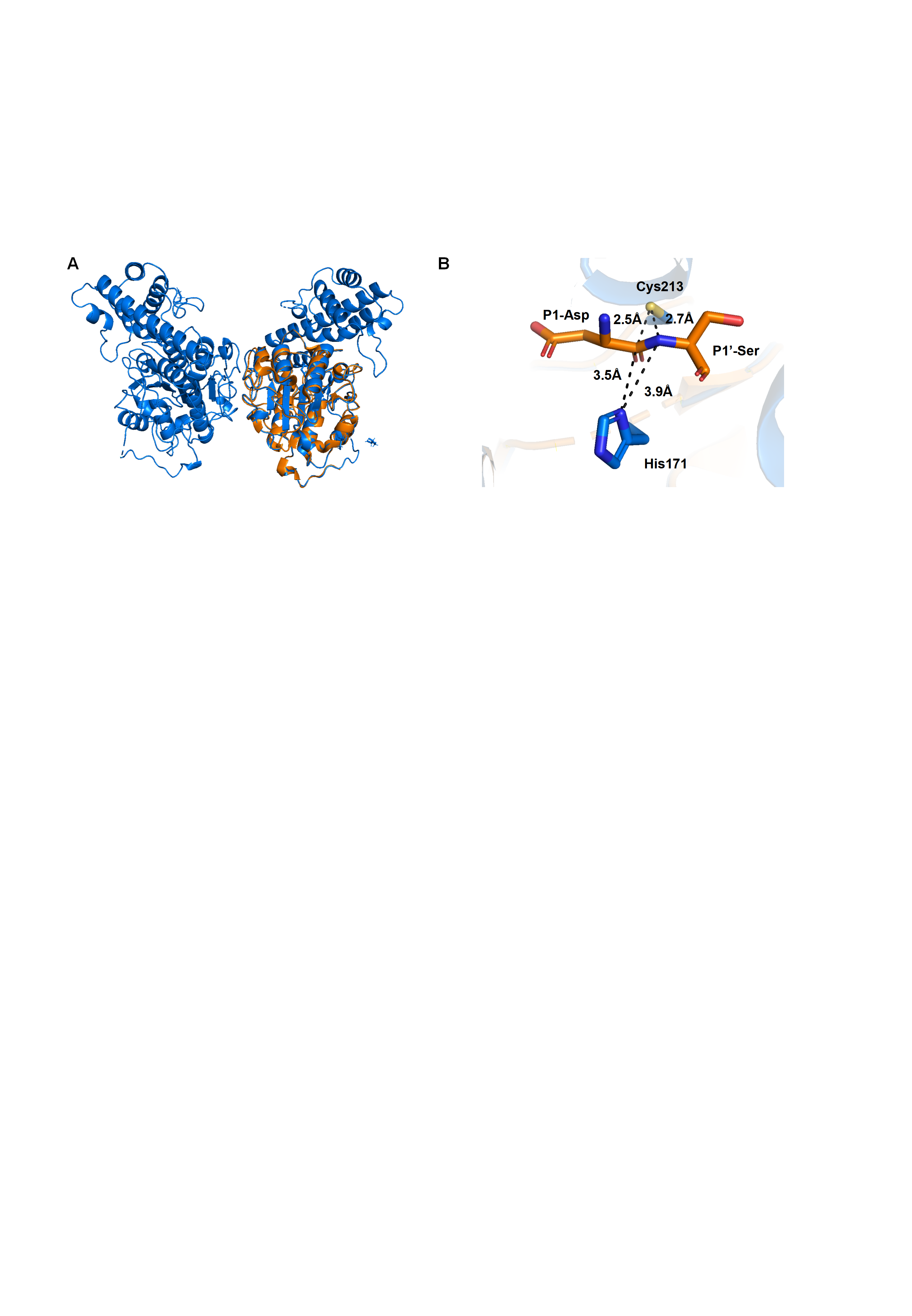

Figure S18 Comparison of the crystal structures of the VdiPAL1 and VyPAL2 core structure. (A) Overall structural alignment of VdiPAL1 (blue) and VyPAL2 (PDB ID: 7F5P, orange). (B) Alignment of the catalytic Cys and His (blue) with the substrate (orange).

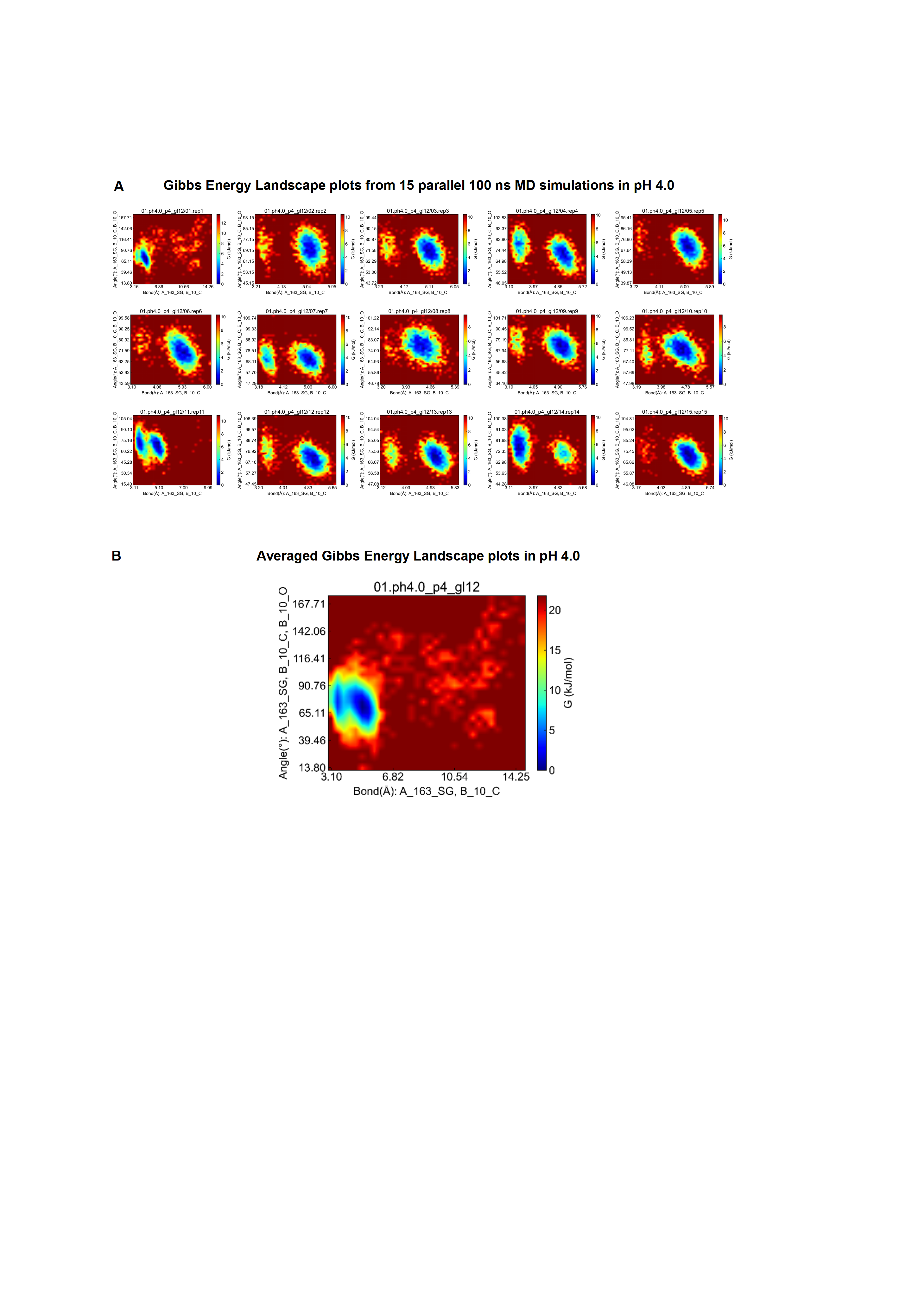
Figure S19 Gibbs free energy landscapes of the Cys nucleophilic attack at pH 4.0. (A) Gibbs free energy landscapes from 15 independent 100 ns MD simulations at pH 4.0. (B) Averaged Gibbs free energy landscape of the 15 replicates. The x-axis represents the distance of the forming carbon-sulfur bond in the intermediate, while the y-axis represents the BD angle. The color scale indicates the Gibbs free energy, with blue representing regions of lower free energy, corresponding to more favorable conformational states.

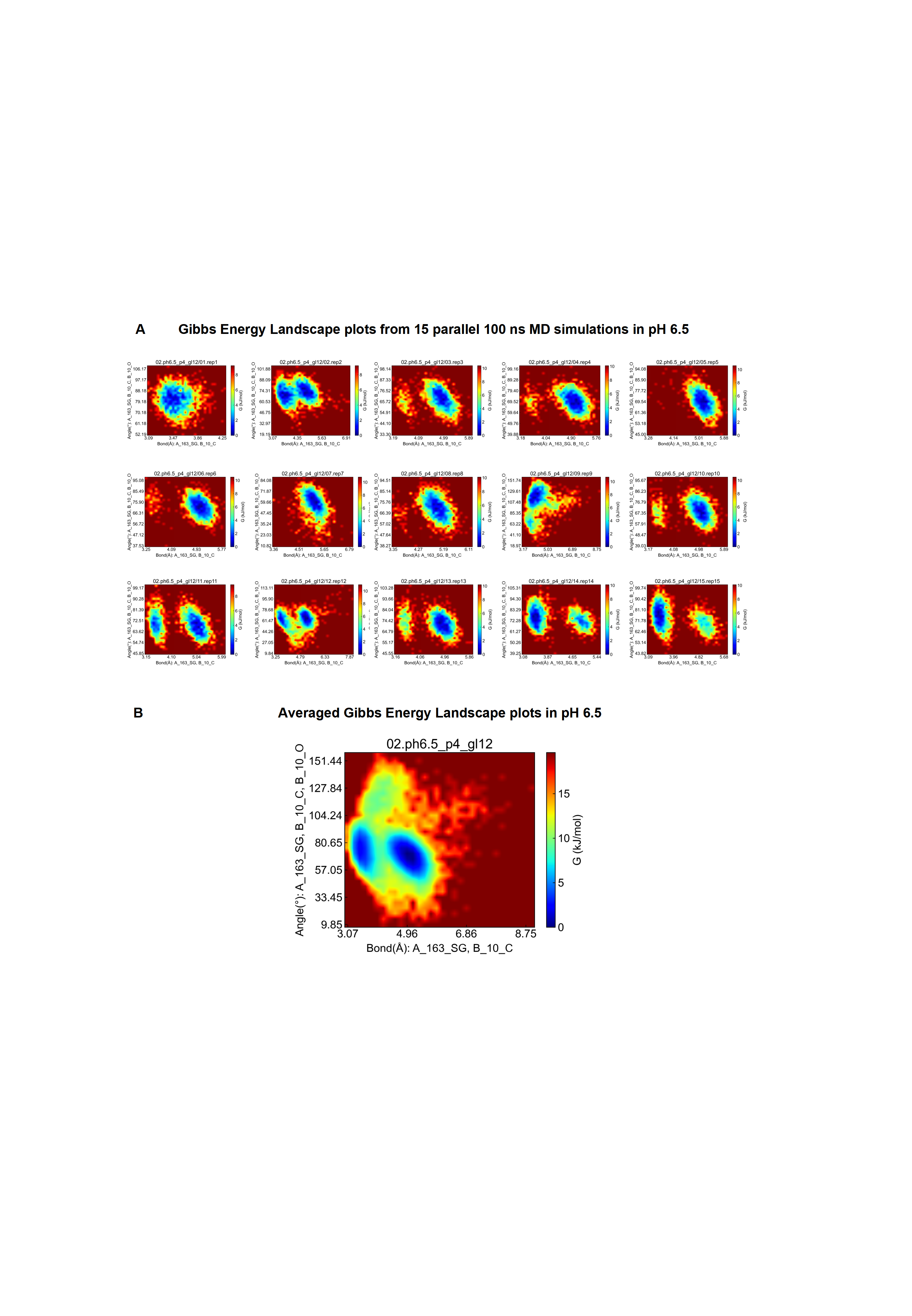

Figure S20 Gibbs free energy landscapes of the Cys nucleophilic attack at pH 6.5. (A) Gibbs free energy landscapes from 15 independent 100 ns MD simulations at pH 6.5. (B) Averaged Gibbs free energy landscape of the 15 replicates. The x-axis represents the distance of the forming carbon-sulfur bond in the intermediate, while the y-axis represents the BD angle. The color scale indicates the Gibbs free energy, with blue representing regions of lower free energy, corresponding to more favorable conformational states.

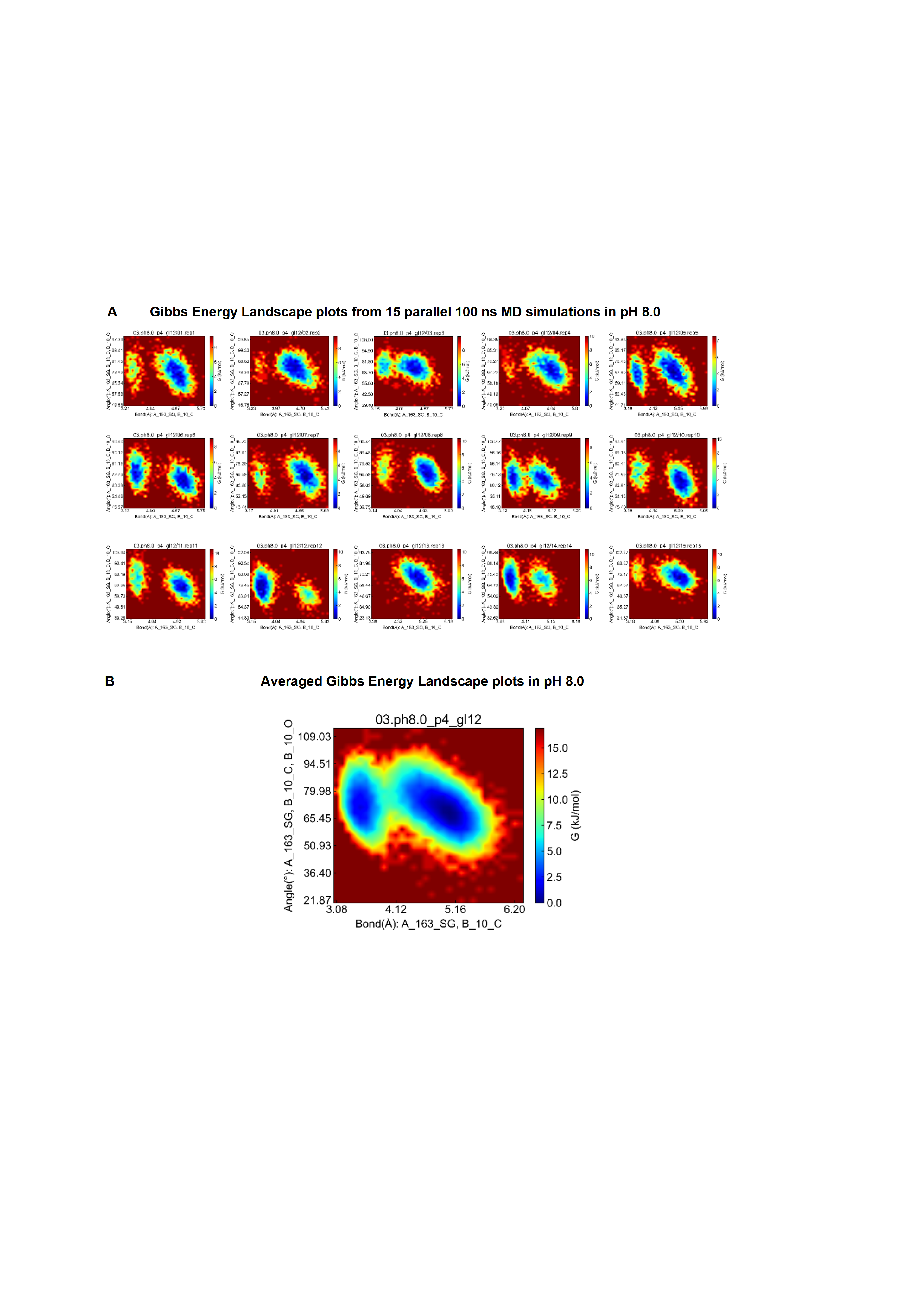

Figure S21 Gibbs free energy landscapes of the Cys nucleophilic attack at pH 8.0. (A) Gibbs free energy landscapes from 15 independent 100 ns MD simulations at pH 8.0. (B) Averaged Gibbs free energy landscape of the 15 replicates. The x-axis represents the distance of the forming carbon-sulfur bond in the intermediate, while the y-axis represents the BD angle. The color scale indicates the Gibbs free energy, with blue representing regions of lower free energy, corresponding to more favorable conformational states.

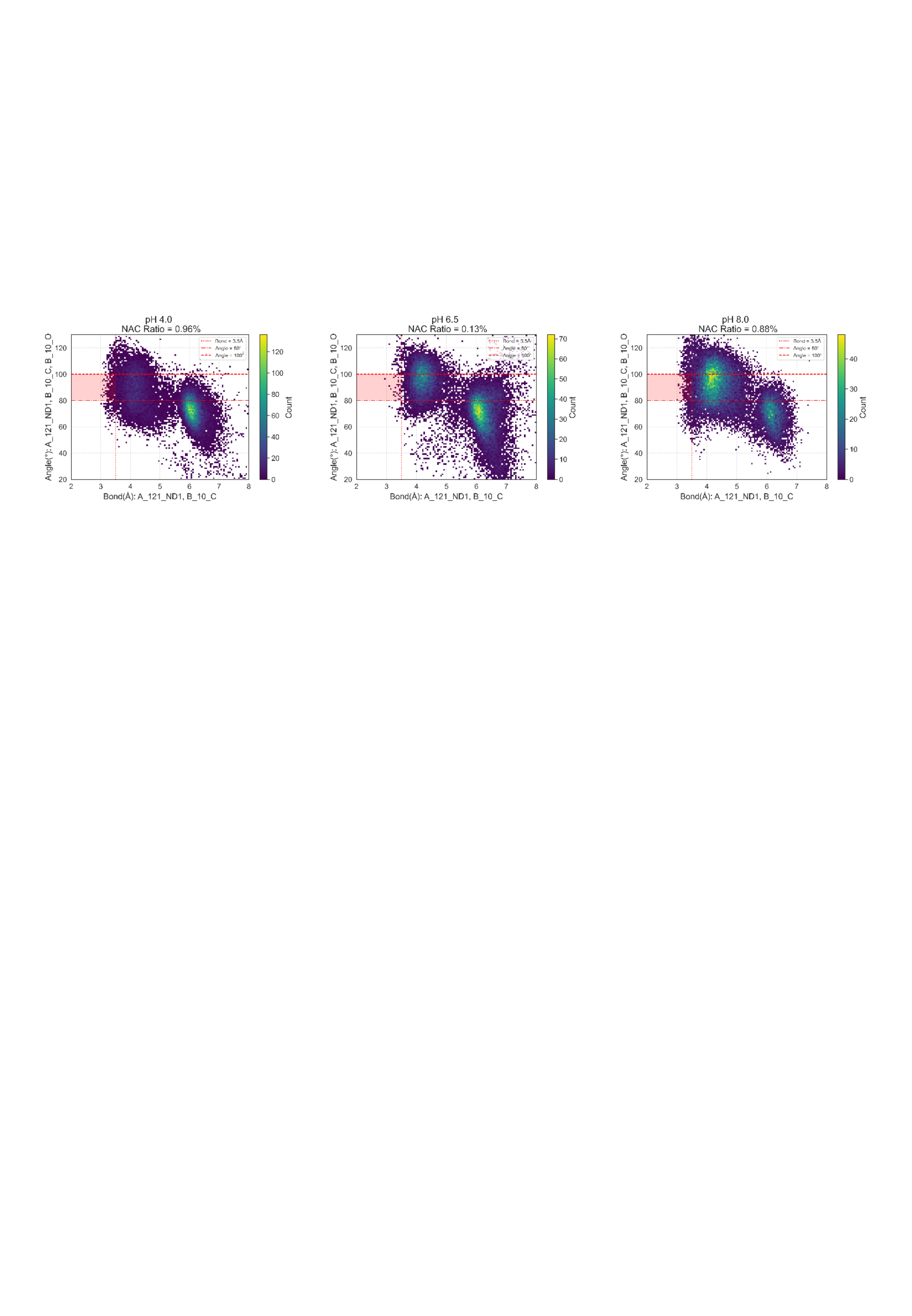

Figure S22 The NAC scatter plot for nucleophilic attack by histidine at different pH 100 ns MD simulations. The 2D histogram illustrating the statistical distribution of distances and angles for conformations of His171 that satisfy the Near-Attack Complex (NAC) criteria, collected from 100 ns × 15 replicated MD simulations at three different pH values. The red highlight indicates the percentage of the population that falls within the NAC geometry.

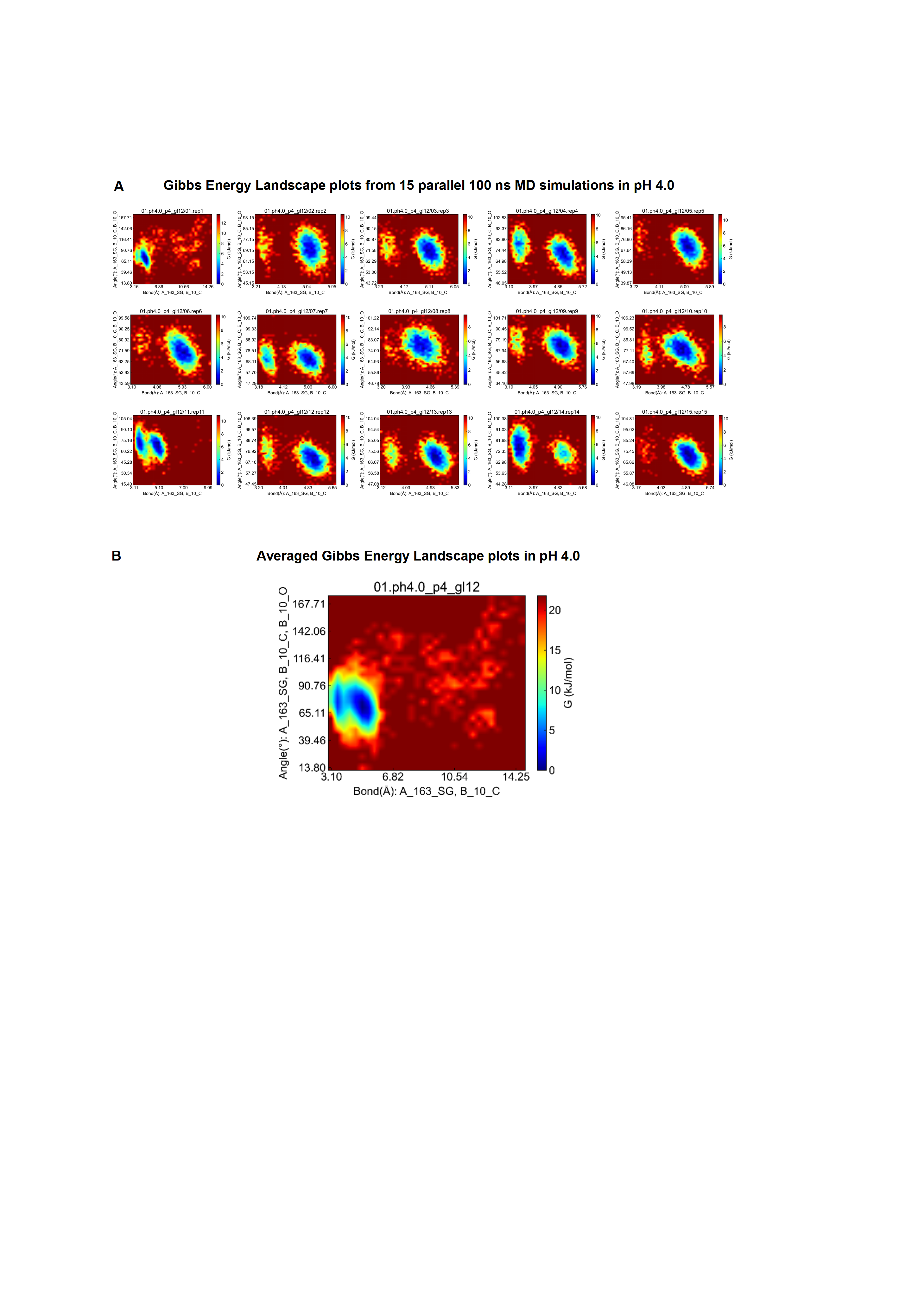

Figure S23 Gibbs free energy landscapes of the His nucleophilic attack at pH 4.0. (A) Gibbs free energy landscapes from 15 independent 100 ns MD simulations at pH 4.0. (B) Averaged Gibbs free energy landscape of the 15 replicates. The x-axis represents the distance of the forming carbon-sulfur bond in the intermediate, while the y-axis represents the BD angle. The color scale indicates the Gibbs free energy, with blue representing regions of lower free energy, corresponding to more favorable conformational states.

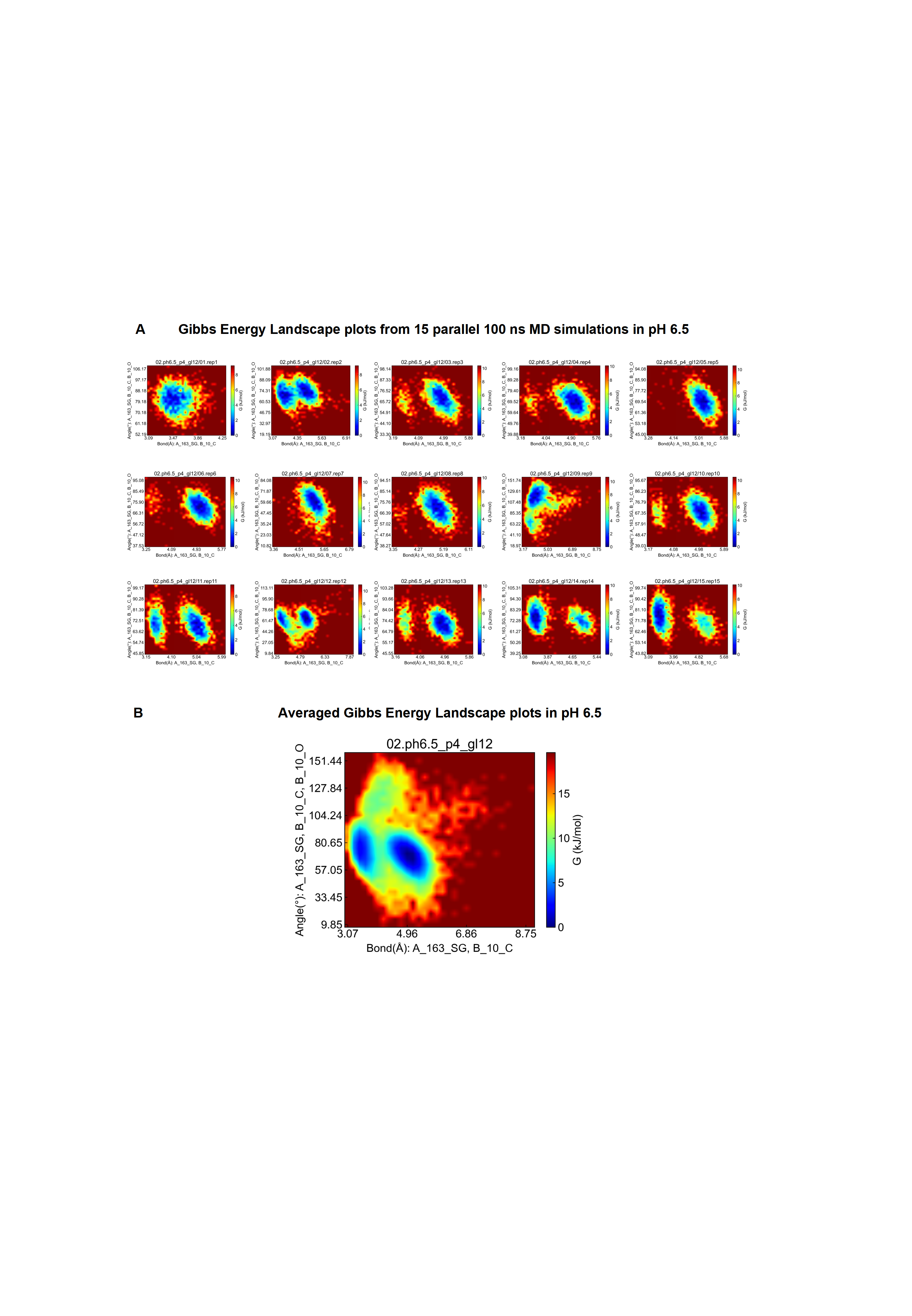

Figure S24 Gibbs free energy landscapes of the His nucleophilic attack at pH 6.5. (A) Gibbs free energy landscapes from 15 independent 100 ns MD simulations at pH 6.5. (B) Averaged Gibbs free energy landscape of the 15 replicates. The x-axis represents the distance of the forming carbon-sulfur bond in the intermediate, while the y-axis represents the BD angle. The color scale indicates the Gibbs free energy, with blue representing regions of lower free energy, corresponding to more favorable conformational states.

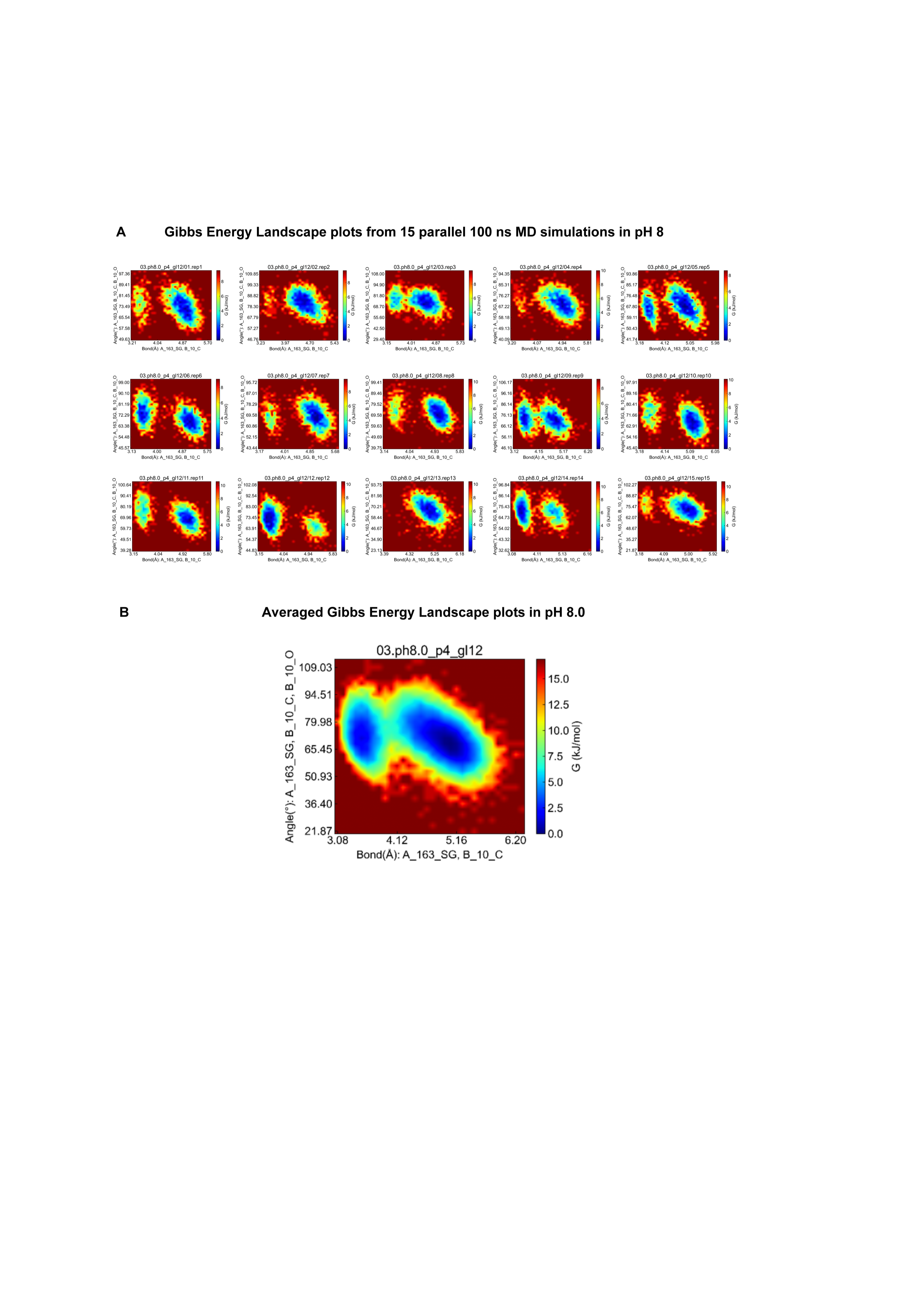

Figure S25 Gibbs free energy landscapes of the His nucleophilic attack at pH 8.0. (A) Gibbs free energy landscapes from 15 independent 100 ns MD simulations at pH 8.0. (B) Averaged Gibbs free energy landscape of the 15 replicates. The x-axis represents the distance of the forming carbon-sulfur bond in the intermediate, while the y-axis represents the BD angle. The color scale indicates the Gibbs free energy, with blue representing regions of lower free energy, corresponding to more favorable conformational states.

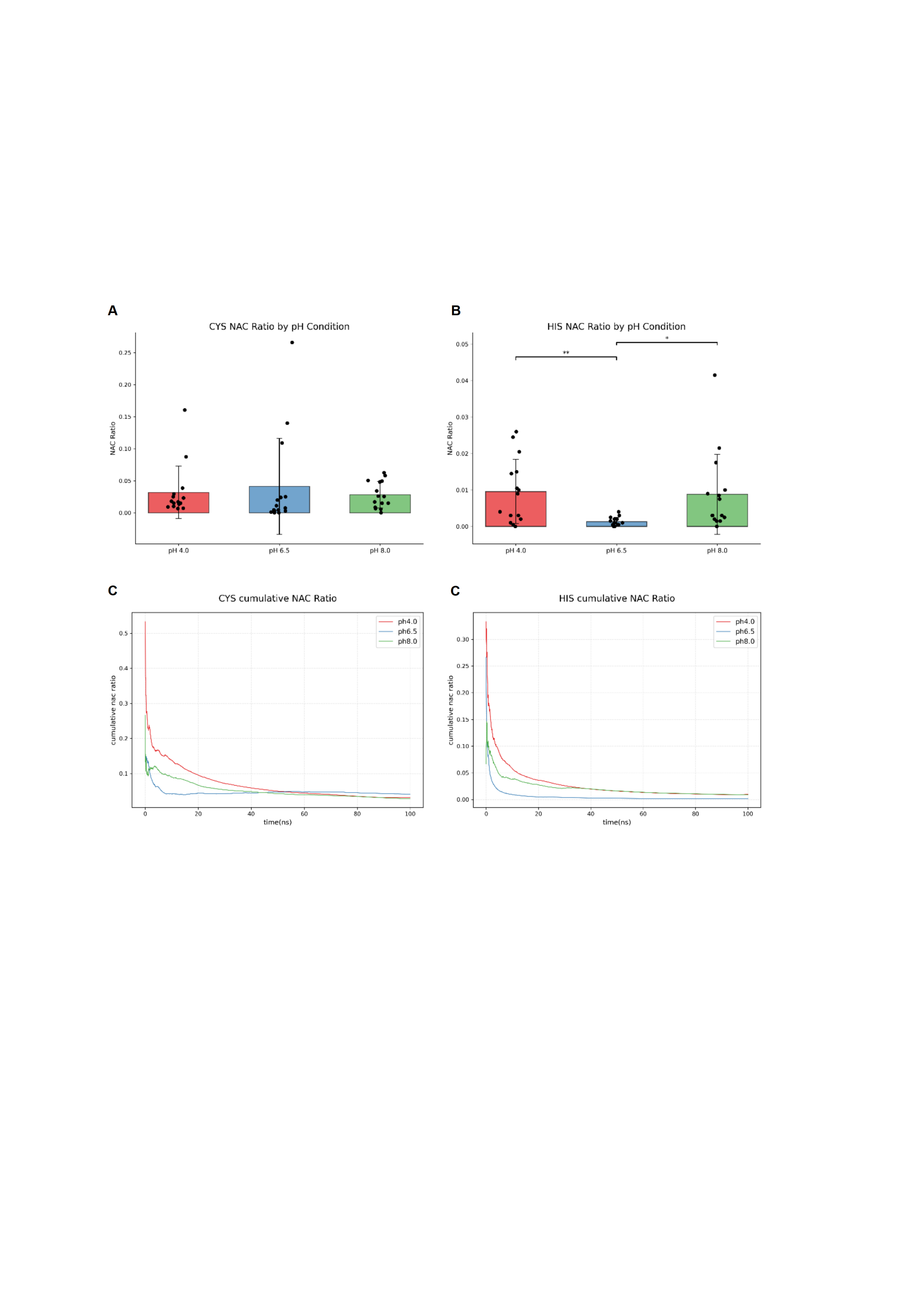

Figure S26 Proportions of NAC with Cys and His acting as attacking residues at three different pH values in 100 ns. (A) Proportions of NACs with Cys as the nucleophile in 15 parallels. (B) Proportions of NACs with His as the nucleophile in 15 parallels. (C) Time evolution of NACs with Cys as the nucleophile during 100 ns simulations. (D) Time evolution of NACs with His as the nucleophile during 100 ns simulations. Data are presented as mean ± SD of *n* = 15 independent simulations. Black dots represent individual data points. Statistical analysis was performed using Student’s t-test. No statistically significant differences were observed between pH conditions for Cys (*p*>0.05). There is a significant difference in the His-mediated catalytic attack between pH 4.0/8.0 and pH 6.5 (*p<0*.05). The NAC ratio and cumulative NAC ratio on all the Y-axis range from 0 to 1.

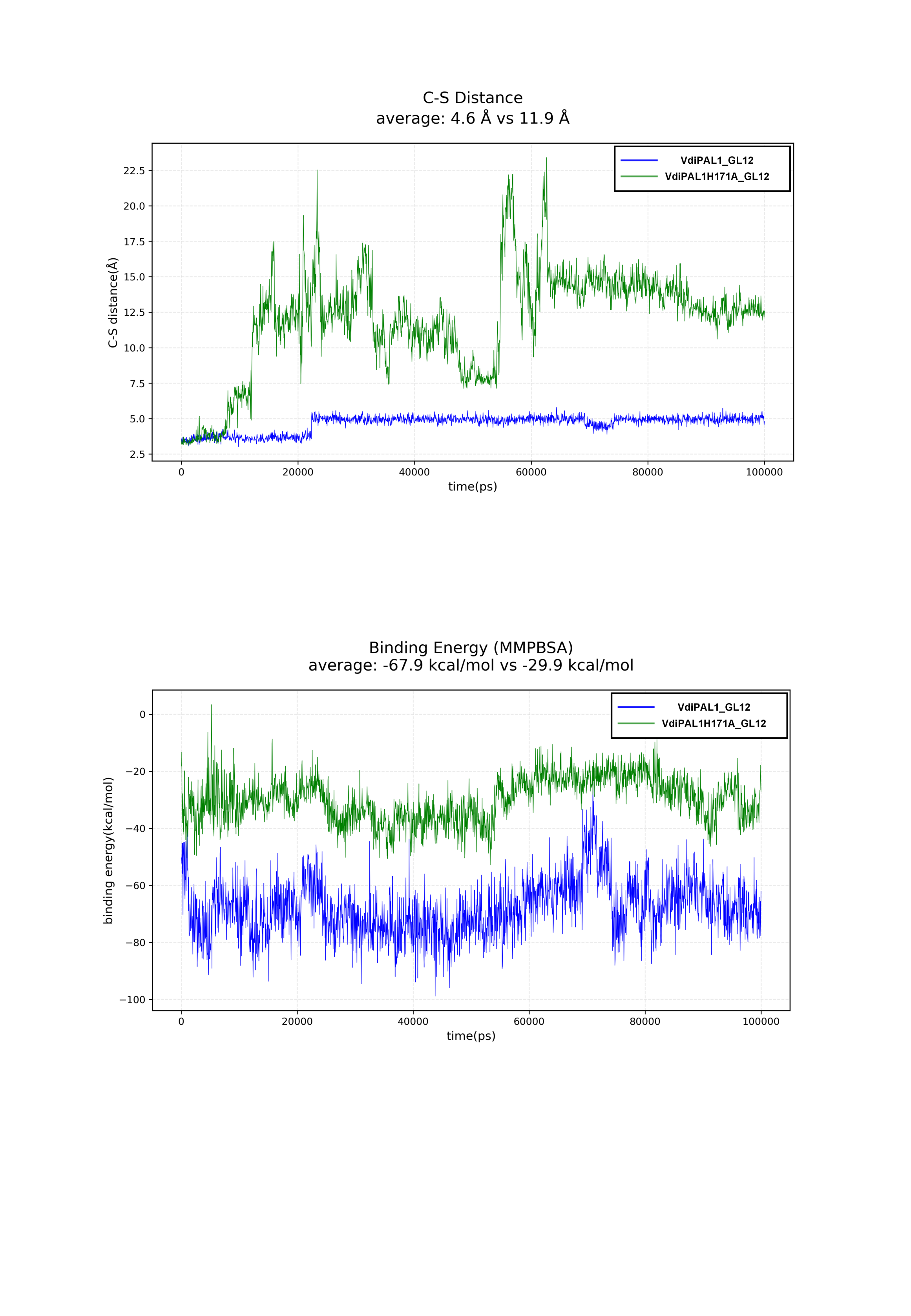

Figure S27 Analysis of the distance between the sulfur atom of catalytic Cys213 and the carbon atom of substrate GL12(Asn10). The blue and green trajectories represent the time evolution of this distance derived from molecular dynamics simulations of the wild-type VdiPAL1 and H171A mutant in complex with the GL12 substrate.

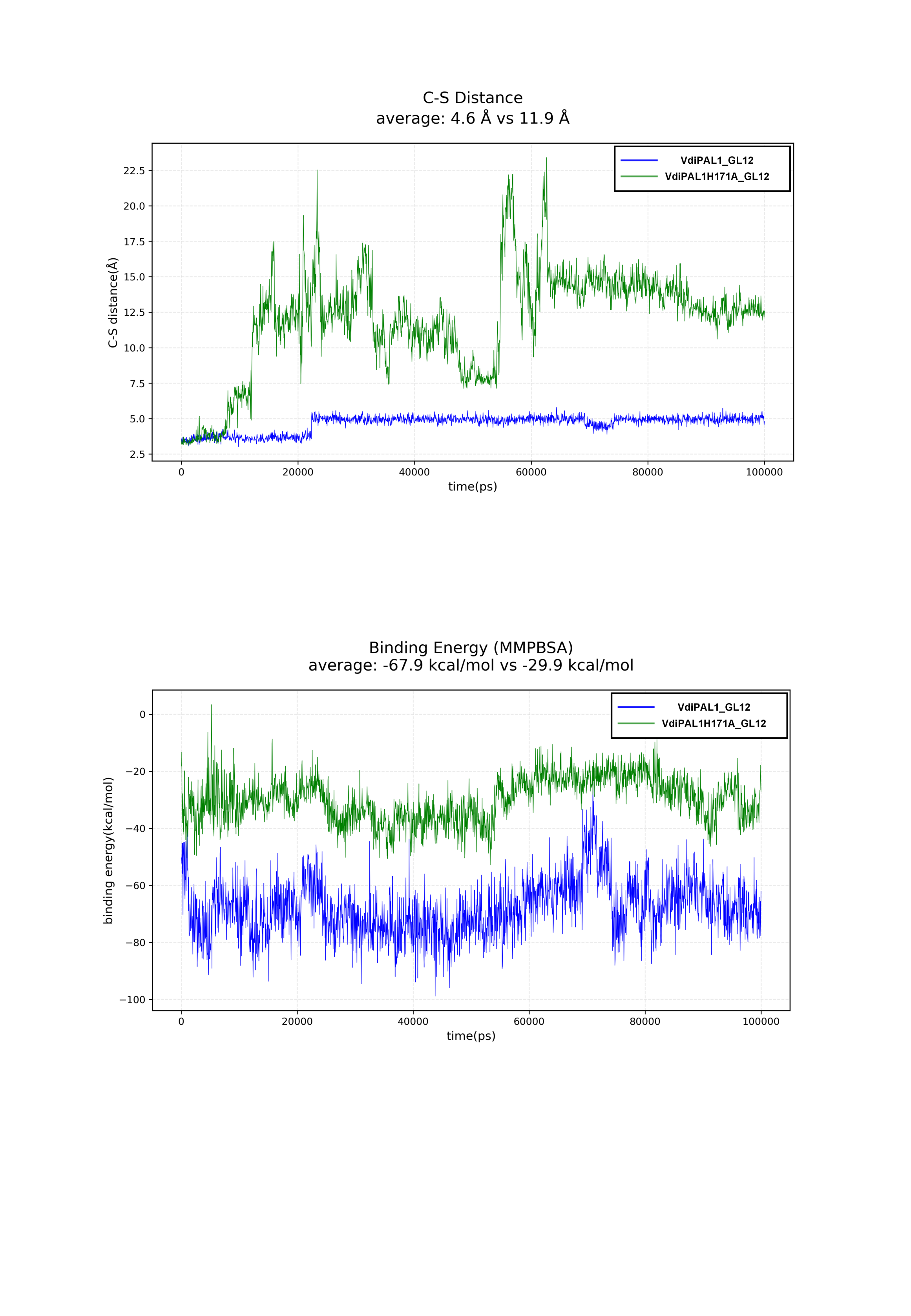

Figure S28 Time-dependent change of the binding free energy between the enzyme and substrate GL12. MM/PBSA calculations were performed based on the MD trajectories. Blue and green curves indicate the energies for the wild-type VdiPAL1 and H171A mutant complexes, respectively.

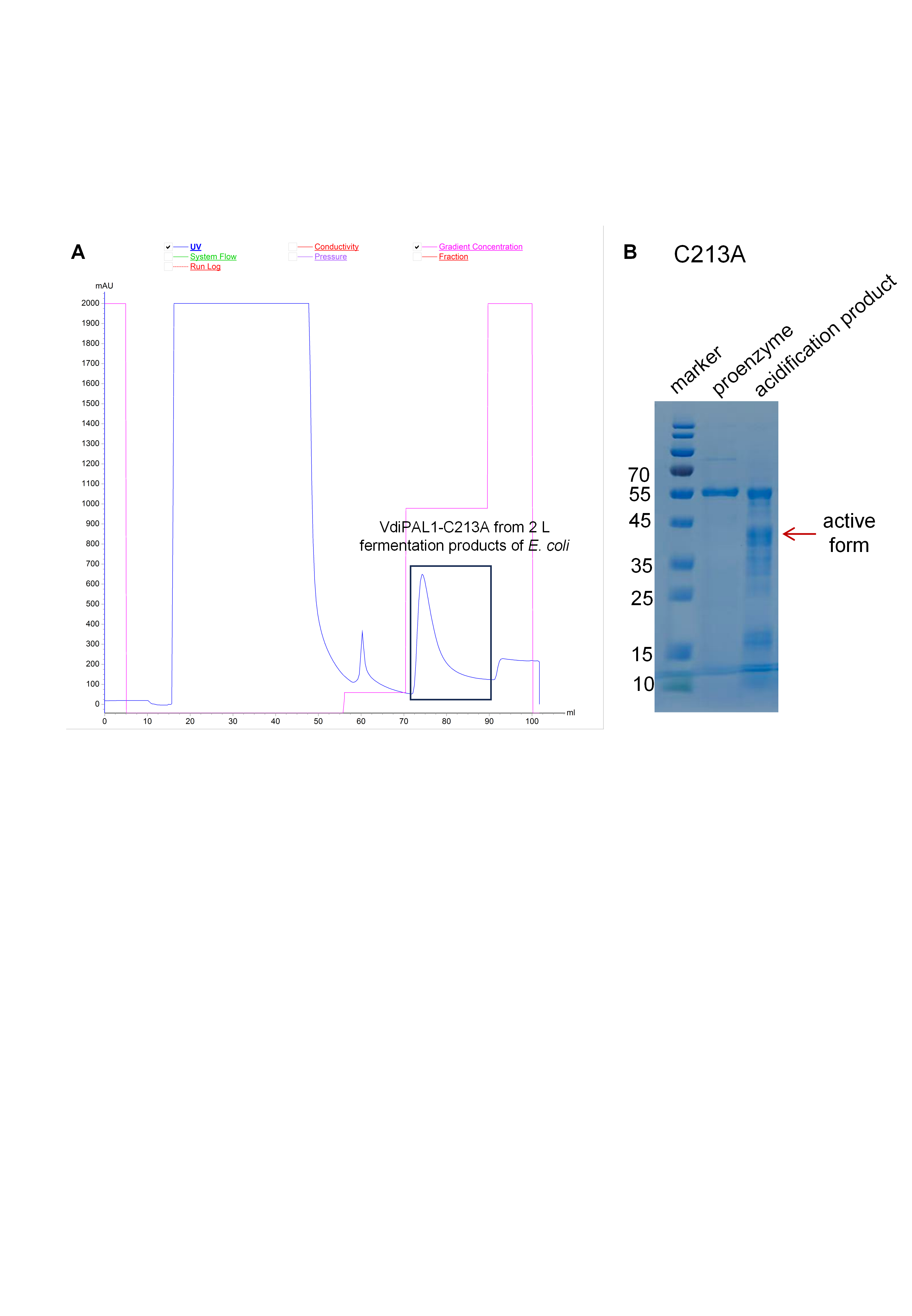

Figure S29 Purification of the VdiPAL1-C213A. (A) The blue line represents the UV absorbance at 280 nm, and the red line indicates the imidazole concentration. Fractions containing the target protein, purified from a 2 L *E. coli* fermentation using Ni-affinity chromatography, are enclosed in the black box. (B) The SDS-PAGE of purified VdiPAL1-C213A proenzyme (55 kDa) and acidification product (pH 4.1, 25 , overnight). The arrow indicates the incomplete active form.

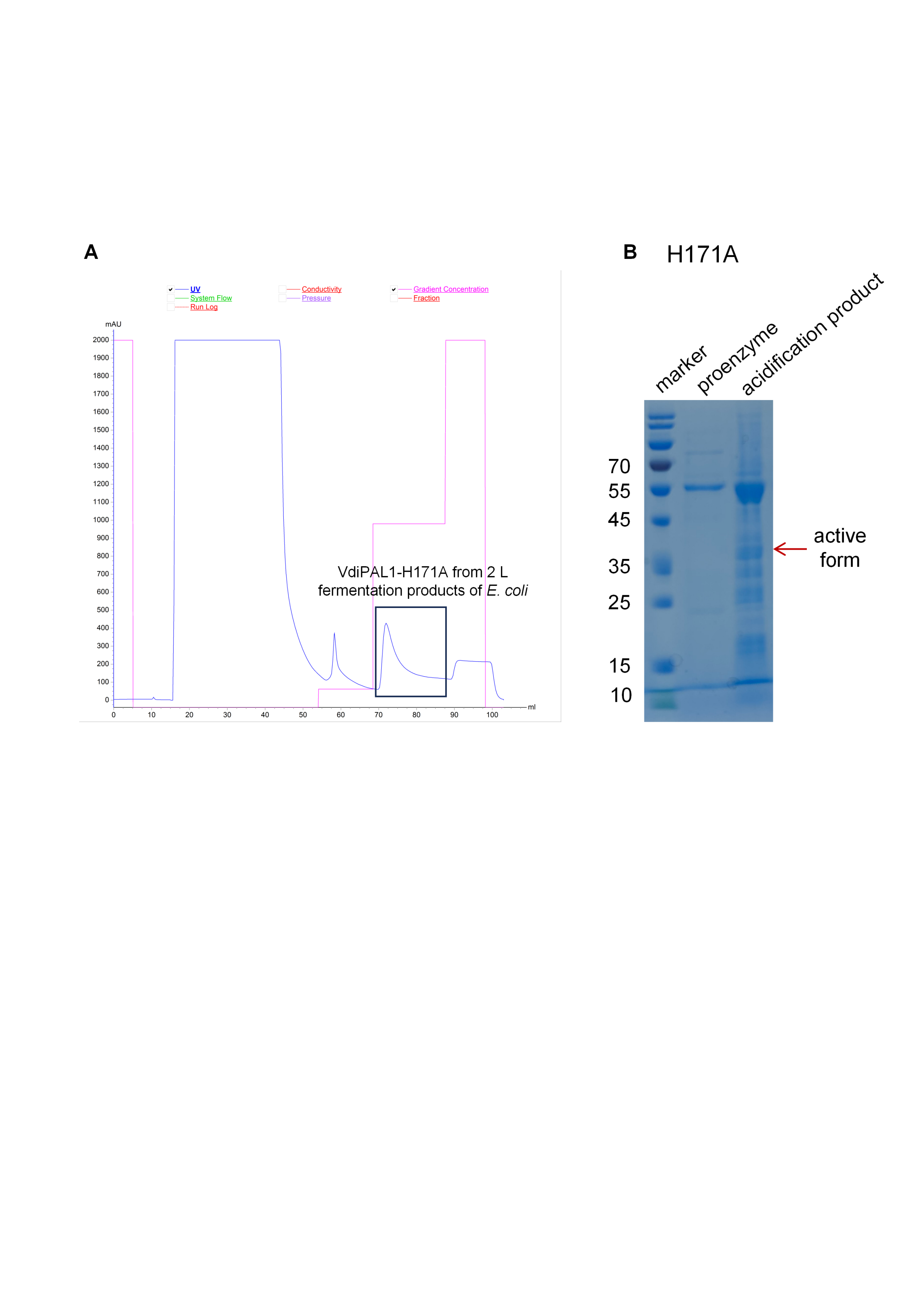

Figure S30 Purification of the VdiPAL1-H171A. (A) The blue line represents the UV absorbance at 280 nm, and the red line indicates the imidazole concentration. Fractions containing the target protein, purified from a 2 L *E. coli* fermentation using Ni-affinity chromatography, are enclosed in the black box. (B) The SDS-PAGE of purified VdiPAL1-H171A proenzyme (55 kDa) and acidification product. The arrow indicates the incomplete active form.

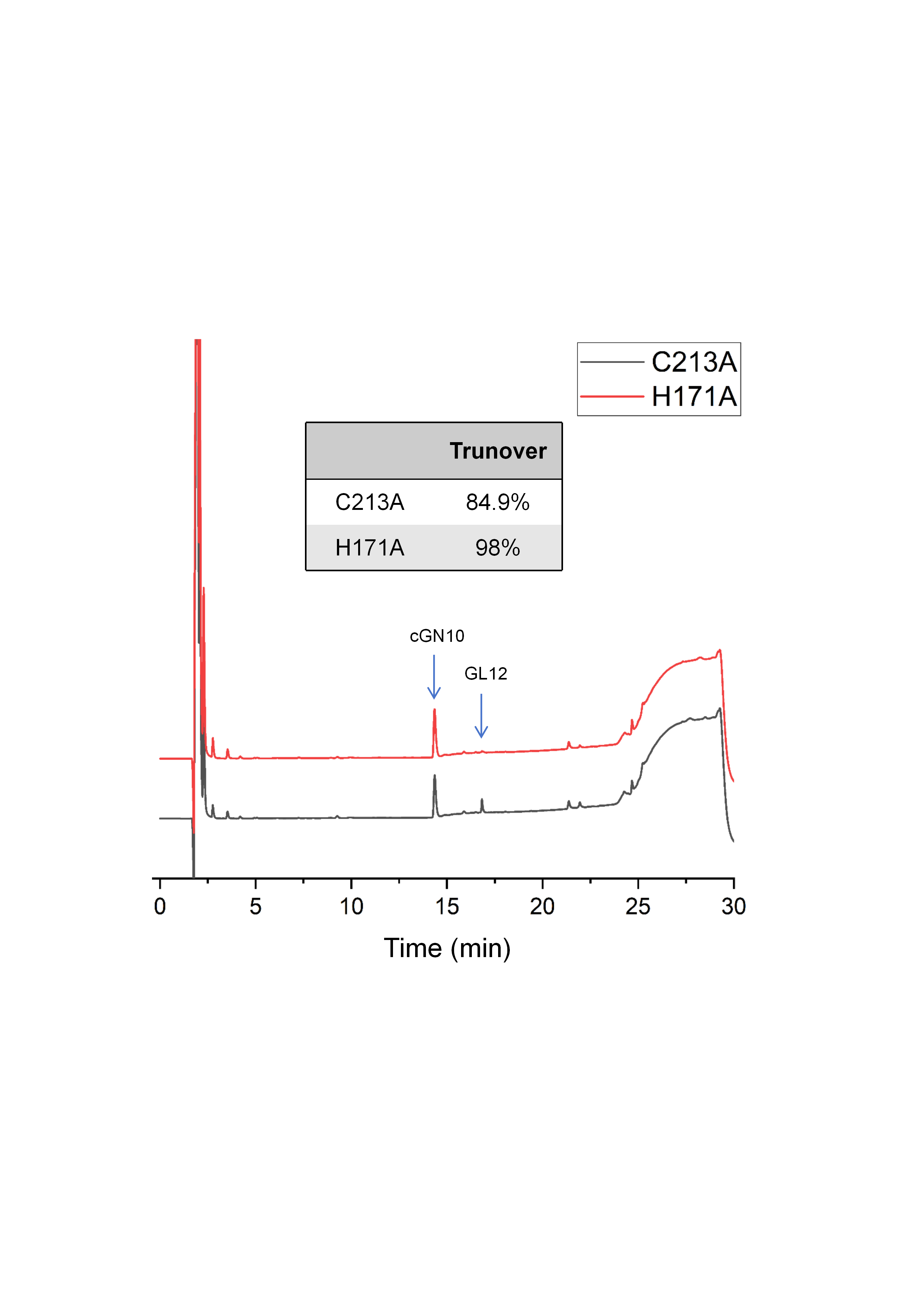

Figure S31 Activity analysis of VdiPAL1 mutants. Cyclization activity of C213A and H171A mutants determined by HPLC. The arrows indicate the substrate (GL12) and the cyclized product (cGN10), respectively.

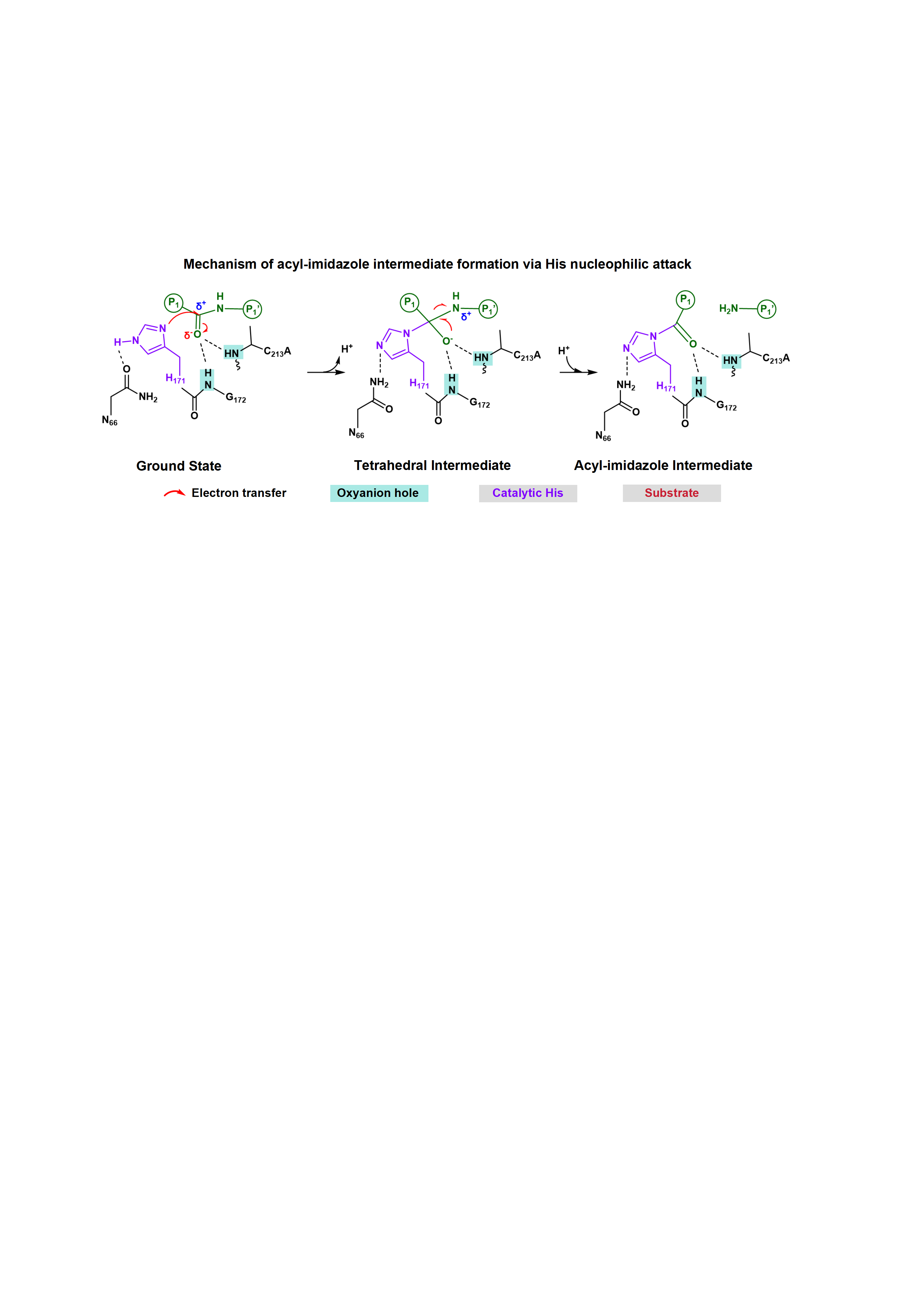

Figure S32 Mechanism of acyl-imidazole intermediate formation via nucleophilic attack. The substrate is shown in green and the catalytic His171 in purple. Key residues (Cys213Ala, Asn66, Gly172) are shown in black. The oxyanion hole, formed by backbone amides of Gly172 and Cys213Ala, is boxed in blue. Electron flow is indicated by red arrows. Schematic of initial attack by His forms an enzyme-substrate acyl-imidazole intermediate.

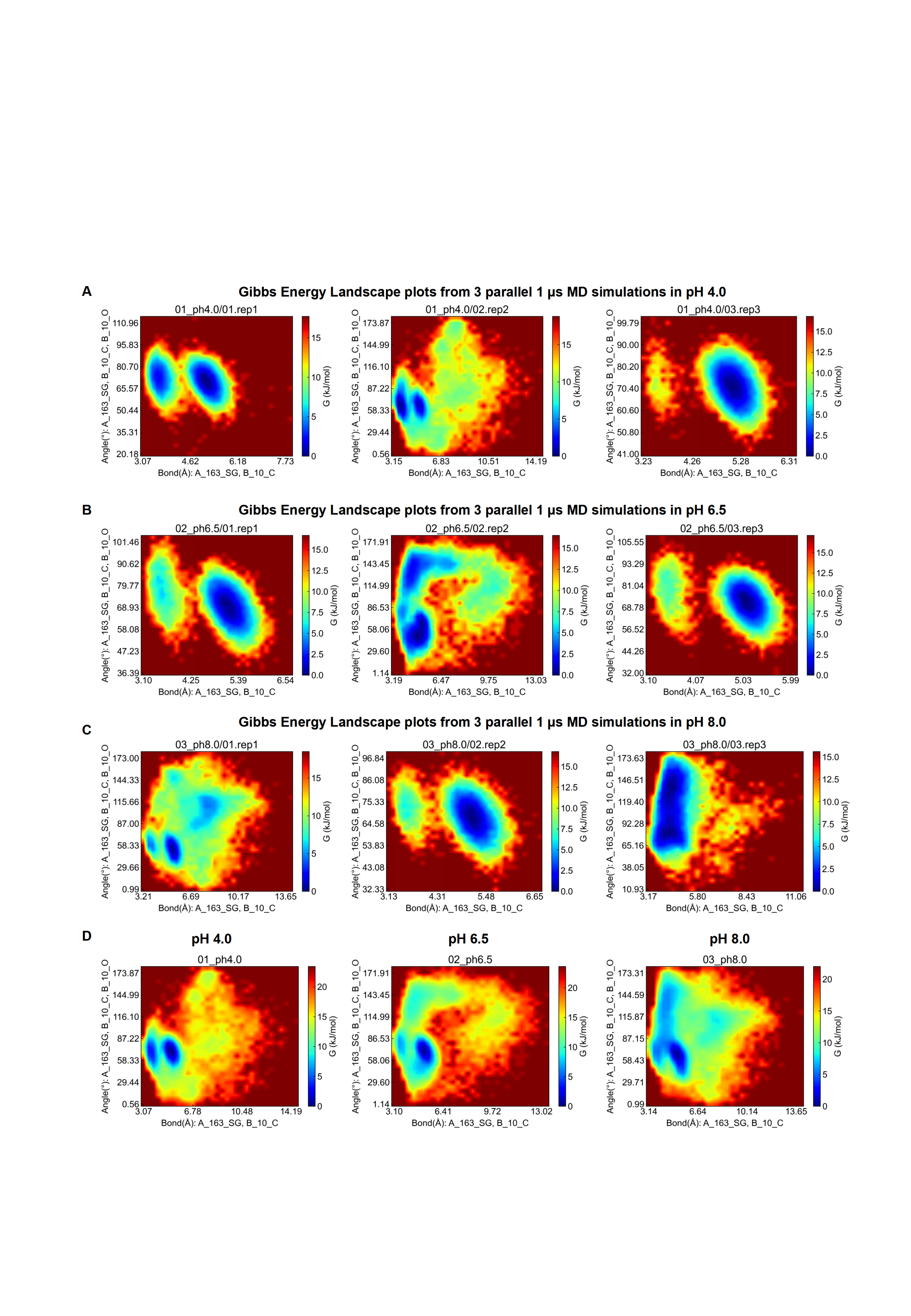

Figure S33 Gibbs free energy landscapes of the Cys nucleophilic attack at three different pH values in 1 μs MD simulations. (A-C) Results from three replicate MD simulations at pH 4.0, 6.5, and 8.0. (D) Averaged landscapes for each of the three pH conditions. The x-axis represents the distance of the forming carbon-sulfur bond in the intermediate, while the y-axis represents the BD angle. The color scale indicates the Gibbs free energy, with blue representing regions of lower free energy, corresponding to more favorable conformational states.

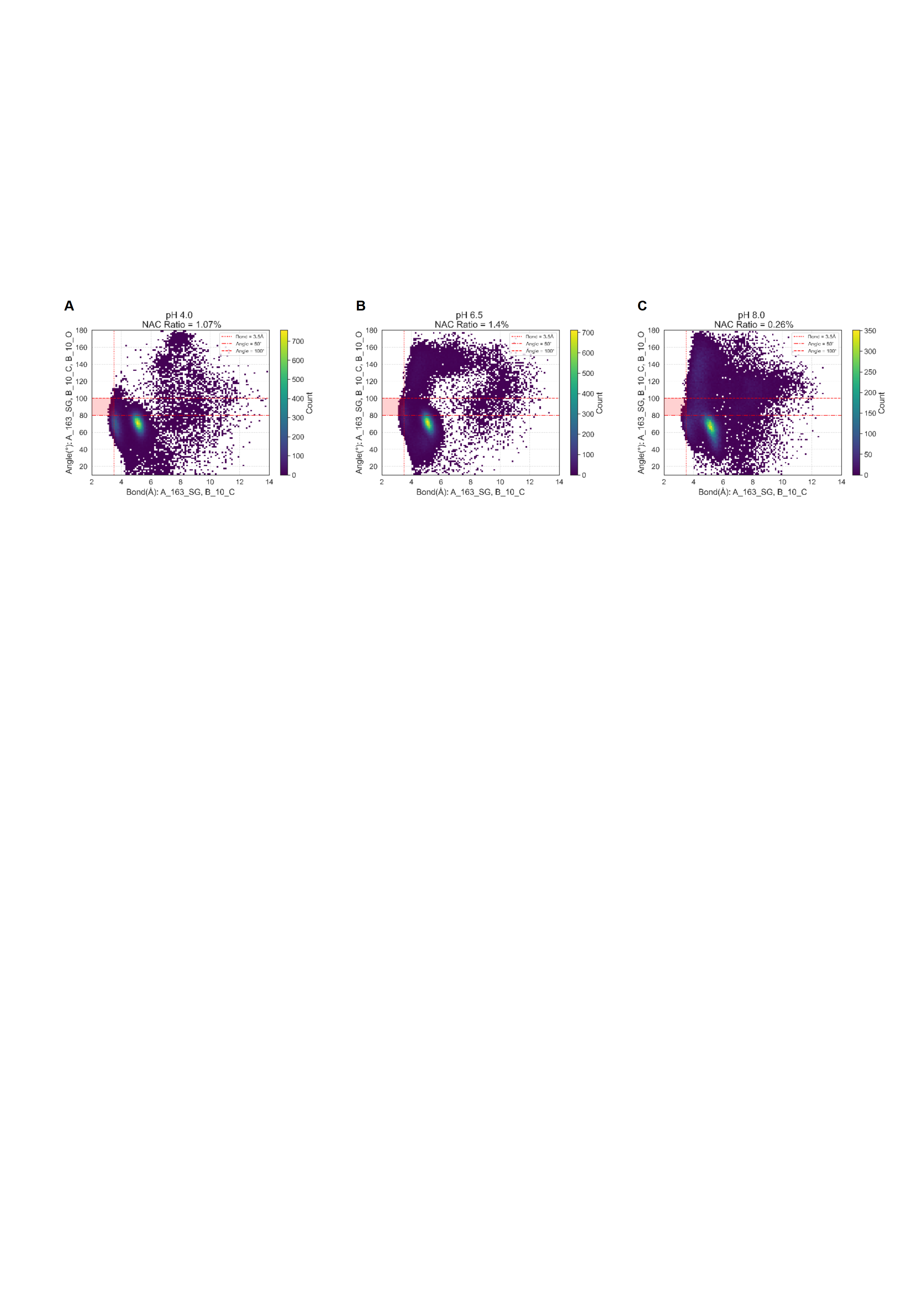

Figure S34 The NAC scatter plot for nucleophilic attack by Cys at different pH in 1 μs MD simulations. The 2D histogram illustrating the statistical distribution of distances and angles for conformations of C213 that satisfy the Near-Attack Complex (NAC) criteria, collected from MD simulations at three different pH values. The red highlight indicates the percentage of the population that falls within the NAC geometry.

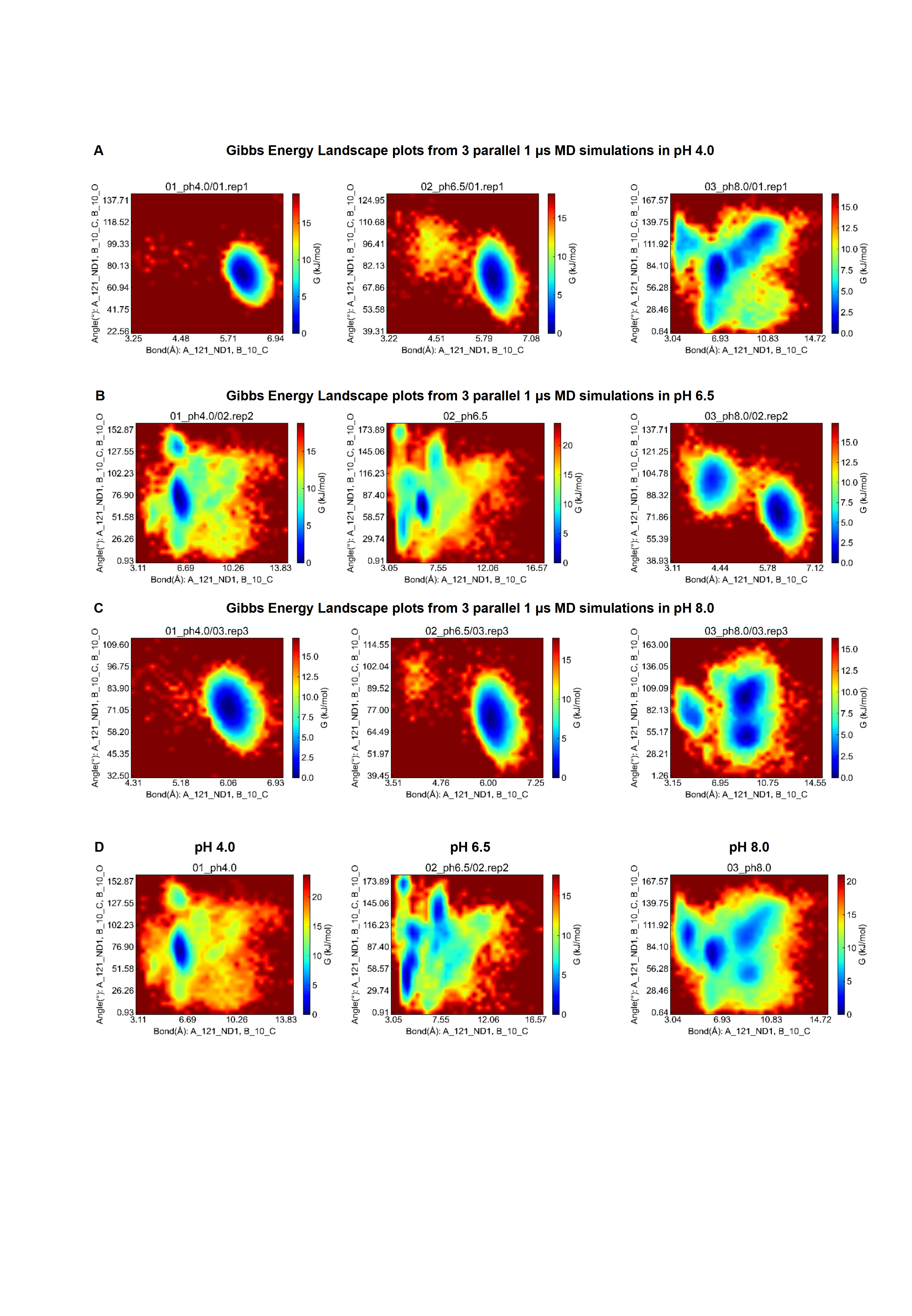

Figure S35 Gibbs free energy landscapes of the His nucleophilic attack at three different pH values in 1 μs MD simulations. (A-C) Results from three replicate MD simulations at pH 4.0, 6.5, and 8.0. (D) Averaged landscapes for each of the three pH conditions. The x-axis represents the distance of the forming carbon-sulfur bond in the intermediate, while the y-axis represents the BD angle. The color scale indicates the Gibbs free energy, with blue representing regions of lower free energy, corresponding to more favorable conformational states.

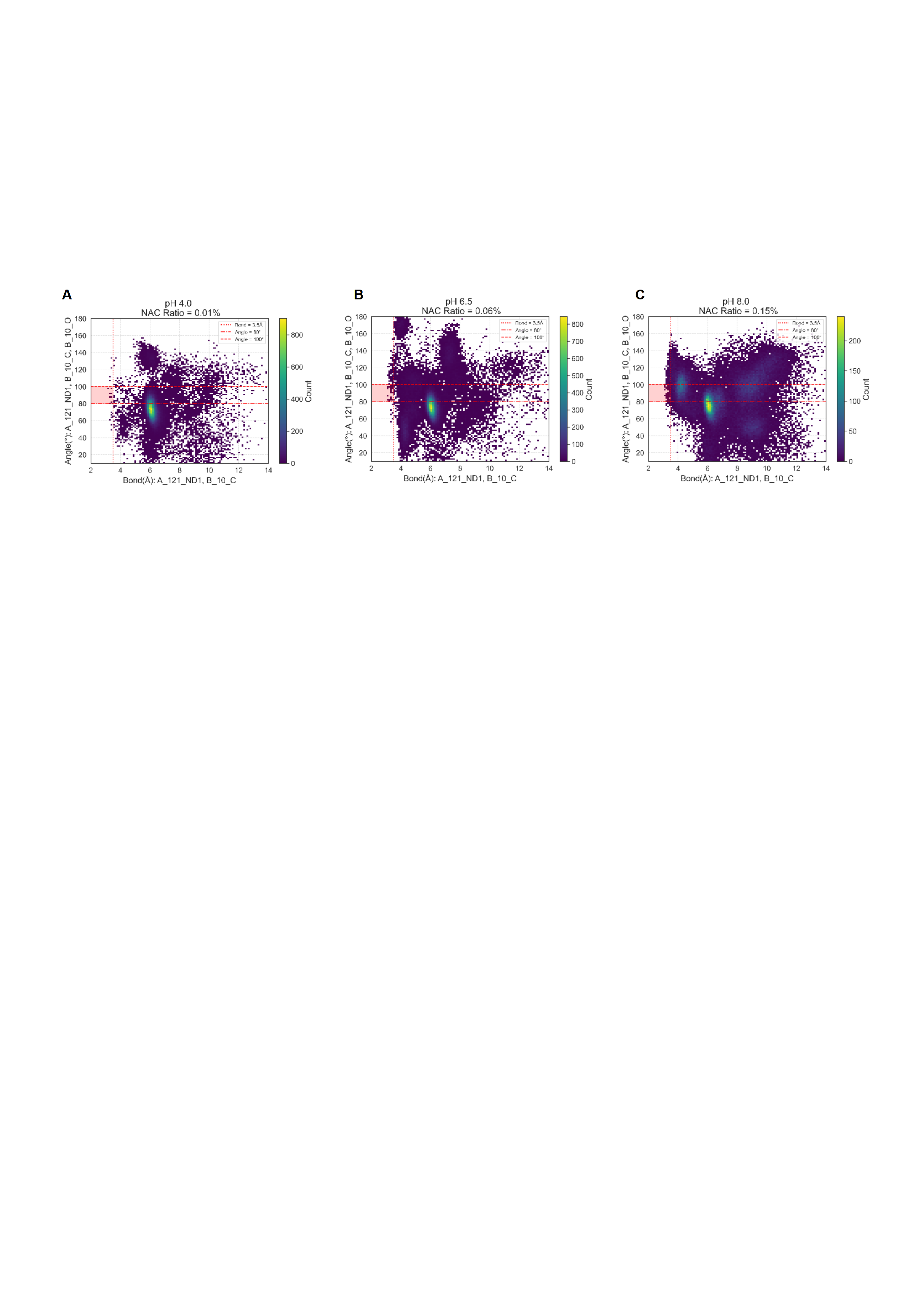

Figure S36 The NAC scatter plot for nucleophilic attack by His at different pH in 1 μs MD simulations. The 2D histogram illustrating the statistical distribution of distances and angles for conformations of H171 that satisfy the Near-Attack Complex (NAC) criteria, collected from MD simulations at three different pH values. The red highlight indicates the percentage of the population that falls within the NAC geometry.

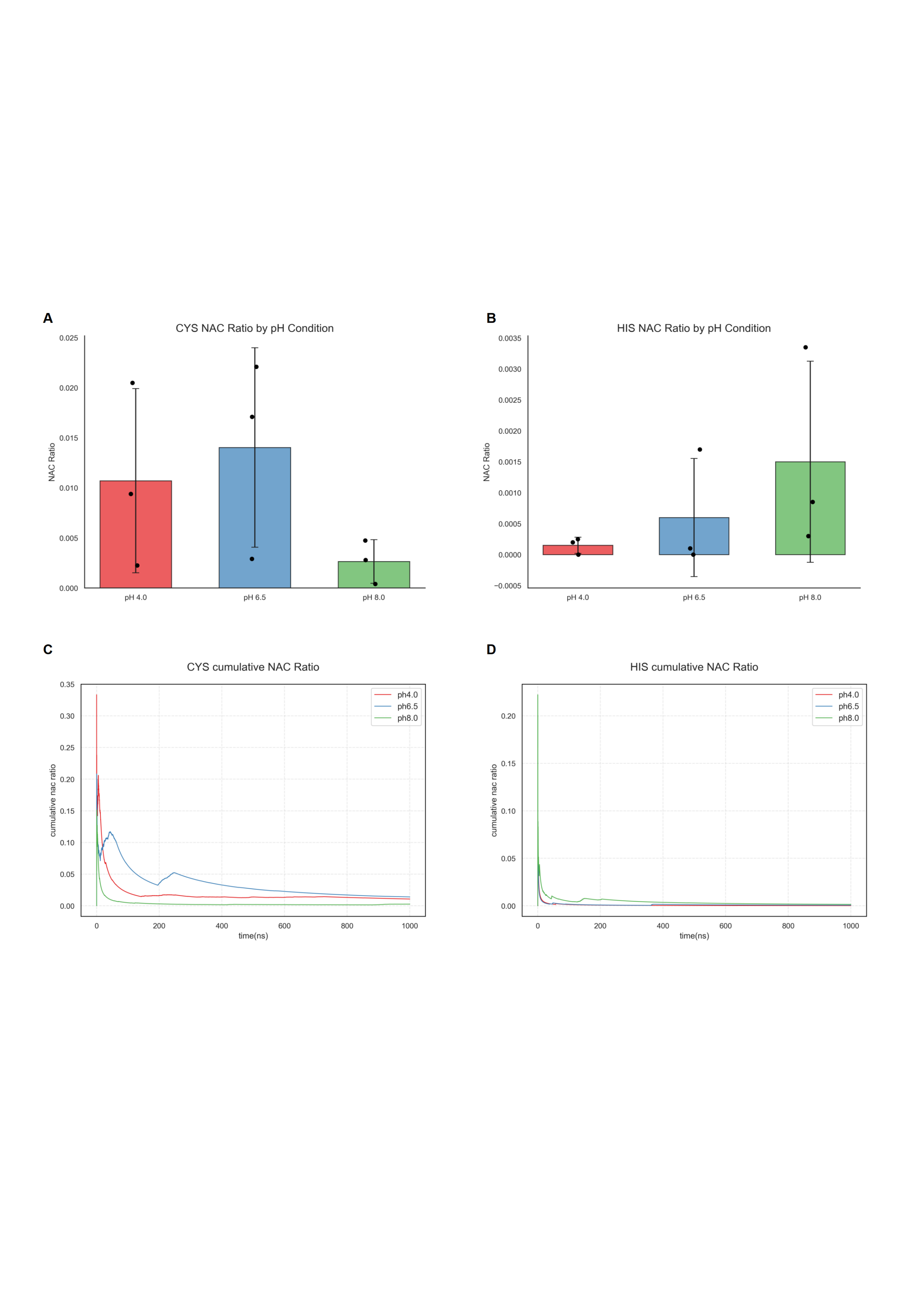

Figure S37 Proportions of NAC with Cys and His acting as attacking residues at three different pH values. (A) Proportions of NACs with Cys as the nucleophile. (B) Proportions of NACs with His as the nucleophile. (C) Time evolution of NACs with Cys as the nucleophile during 1 μs simulations. (D) Time evolution of NACs with His as the nucleophile during 1 μs simulations. Data are presented as mean ± SD of *n*=3 independent simulations. Black dots represent individual data points. Statistical analysis was performed using Student’s t-test. No statistically significant differences were observed between pH conditions for either Cys or His groups (all *p*>0.05). The NAC ratio and cumulative NAC ratio on all the Y-axis range from 0 to 1.

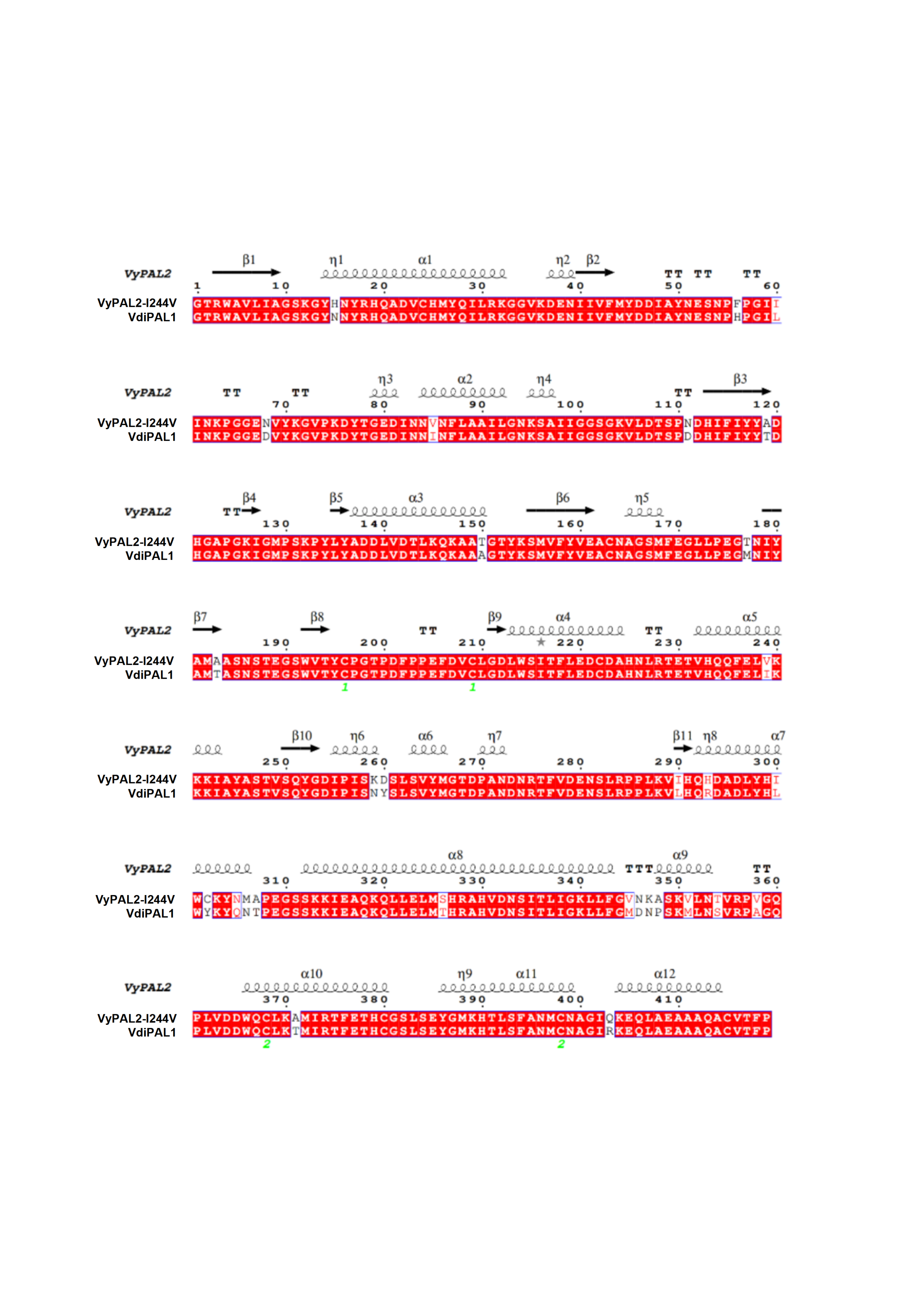
Figure S38 Sequence comparison of VdiPAL1 and VyPAL2. The figure below shows the sequence alignment of VdiPAL1 and VyPAL2. The sequence similarity between VdiPAL1 and VyPAL2 is as high as 92%, with only 30 amino acid residues differing according to the alignment results.

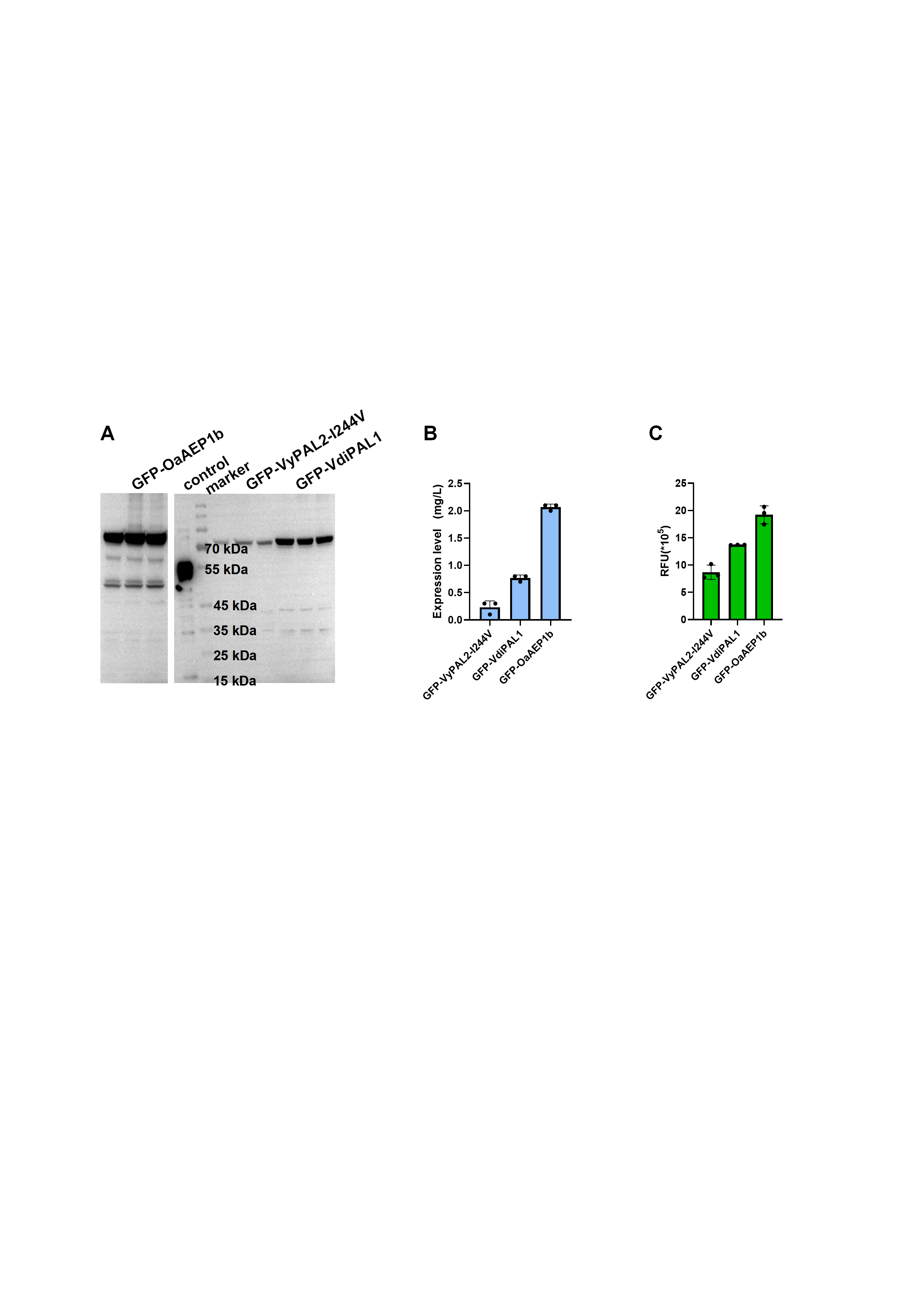

Figure S39 Assessing the correlation between expression levels and fluorescence signals of three PAL-GFP fusion proteins. (A) The WB results are the expression of VdiPAL1, VyPAL2-I244V and OaAEP1b-C247A alone and in fusion with GFP. The target protein was at 78 kDa. (B) Bar chart of expression levels of the three fusion proteins. (C) Bar chart of GFP fluorescence signals of the three fusion proteins.

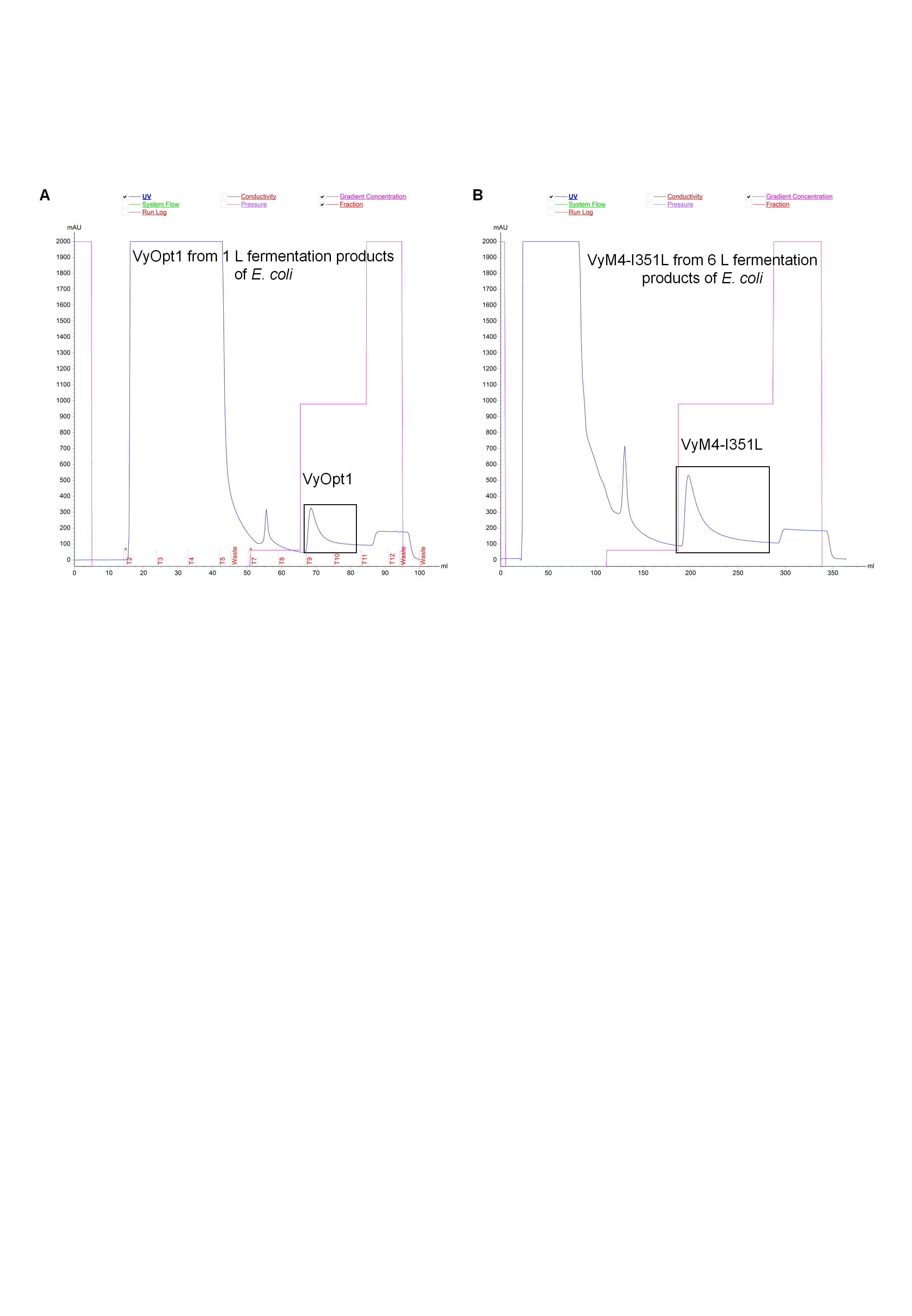

Figure S40 Protein purification results of VyOpt1 and VyM4-I351L using an AKTA-start. (A) Ni-NTA purification chromatogram of 1 L *E. coli* fermentation product of VyOpt1. (B) Ni-NTA purification chromatogram of 6 L *E. coli* fermentation product of VyM4-I351L. Each boxed fractions contain the target enzyme protein.

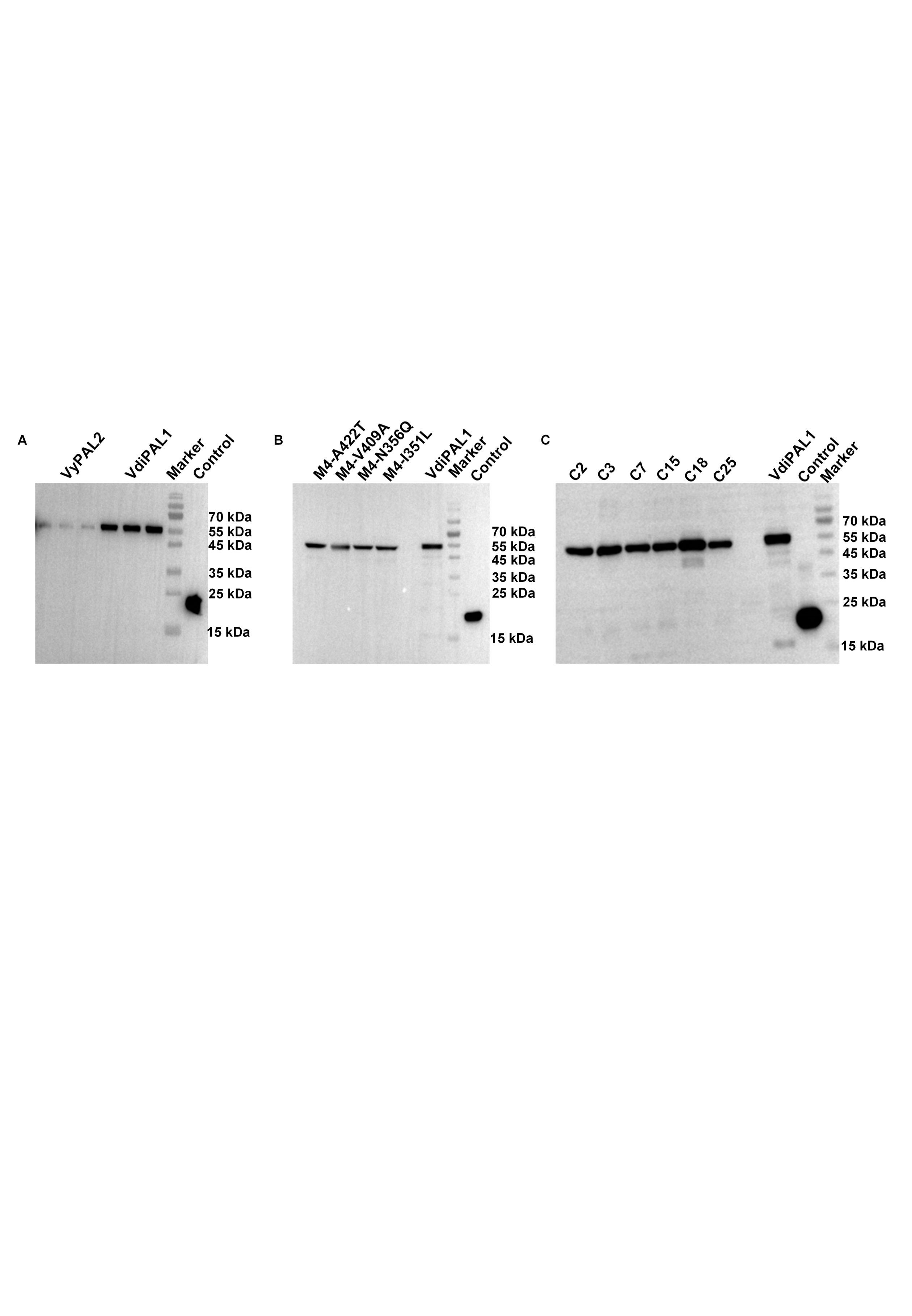

Figure S41 Western Blot (WB) analysis of VyPAL2 wild-type and its mutants. (A) WB of VyPAL2 and VdiPAL1. The target protein was at 55 kDa. (B) WB comparing the expression levels of the VyPAL2 combinatorial mutants from the fourth round of greedy iteration with VdiPAL1. (C) WB showing the expression levels of the computationally designed combinatorial mutant strains corresponding to the top 6 fluorescence signals. Each WB includes a control with an equal amount of enzyme as an internal reference.

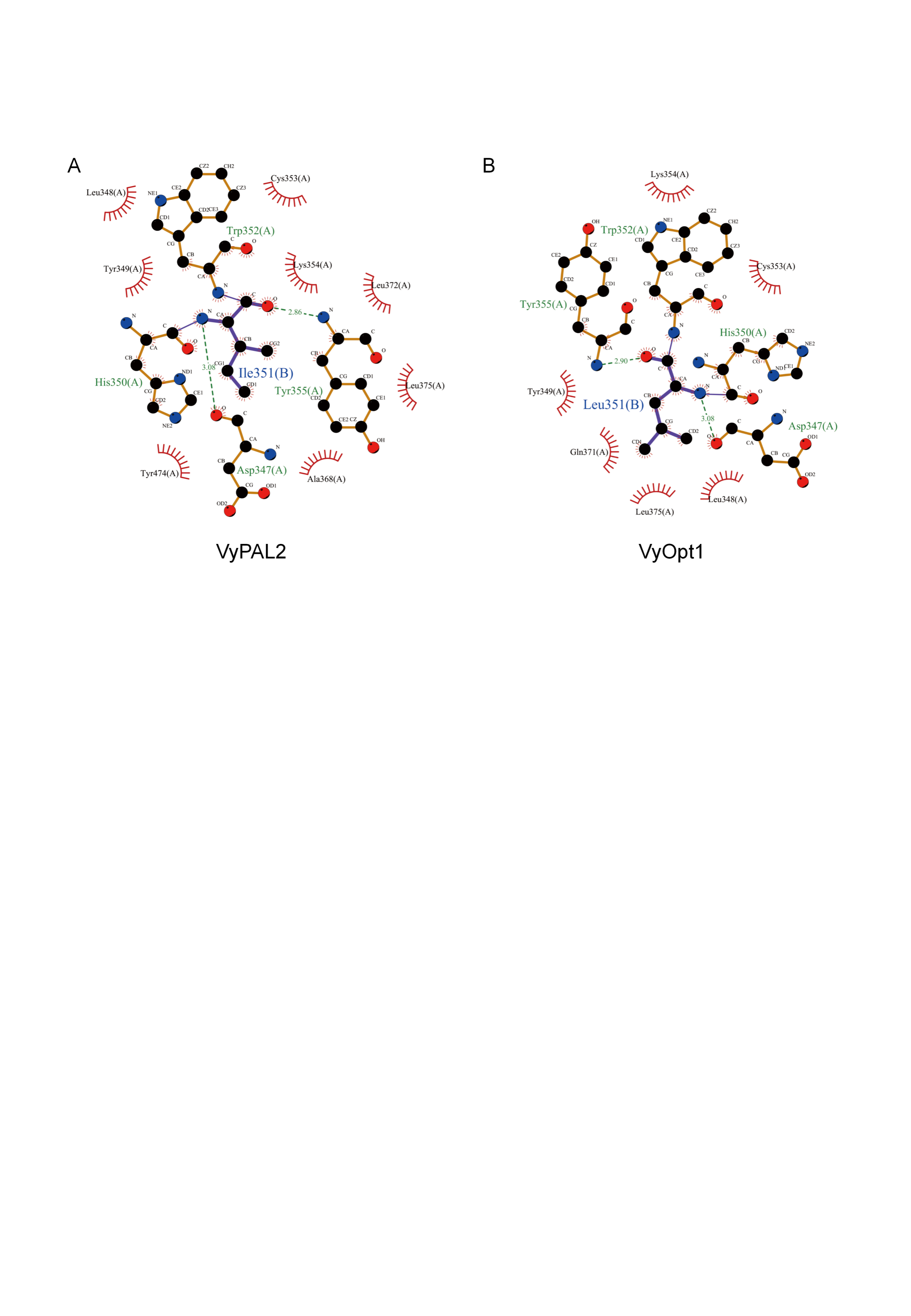

Figure S42 LigPlot diagram of the I351L mutant and its interactions with surrounding residues. (A) The LigPlot diagram illustrates the interaction network of Ile351 with surrounding residues in wild-type VyPAL2 (B) The LigPlot diagram illustrates the interaction of Leu351 with surrounding residues in mutant VyOpt1. Green dashed lines represent hydrogen bonds, and red spoked arcs indicate hydrophobic interactions. Carbon, oxygen, and nitrogen atoms are shown as black, red, and blue spheres, respectively.

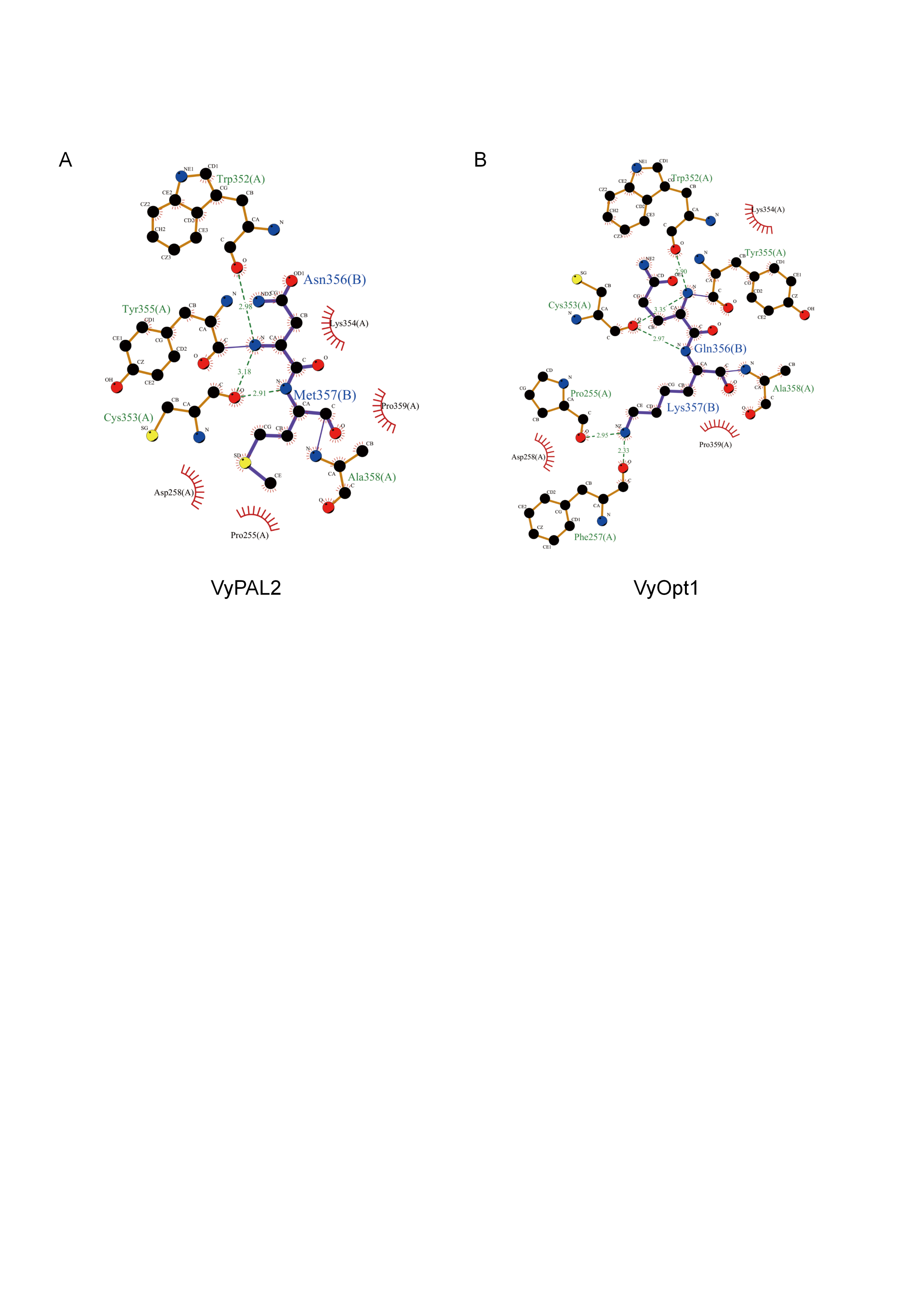
 Figure S43 LigPlot diagram of the N356Q and M357K mutants and its interactions with surrounding residues. (A) The interaction network of N356Q and M357K with surrounding residues in wild-type VyPAL2 (B) The interaction of N356Q and M357K with surrounding residues in mutant VyOpt1.

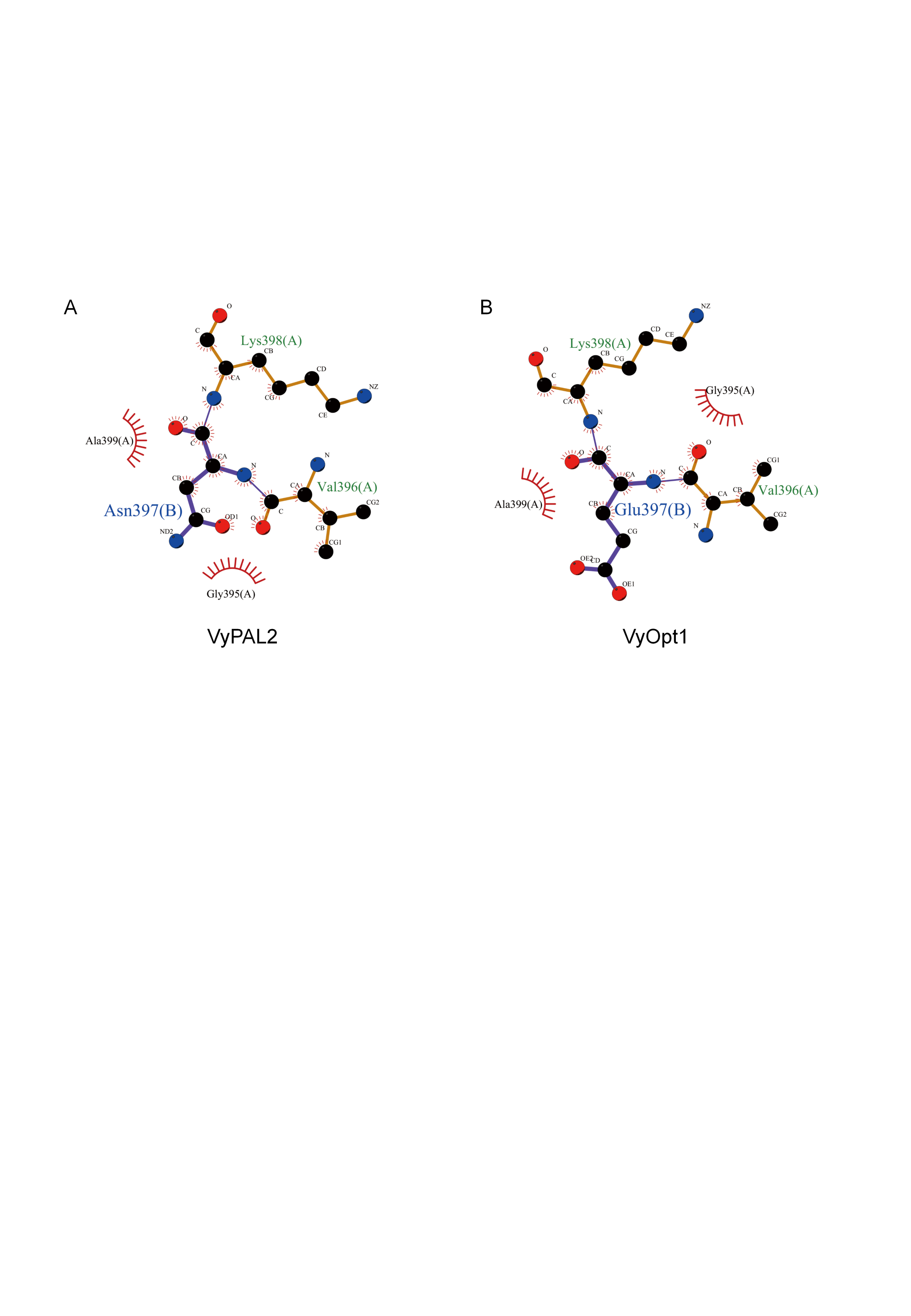

Figure S44 LigPlot diagram of the N397E mutant and its interactions with surrounding residues. (A) The interaction network of N397E with surrounding residues in wild-type VyPAL2 (B) The interaction of N397E with surrounding residues in mutant VyOpt1.

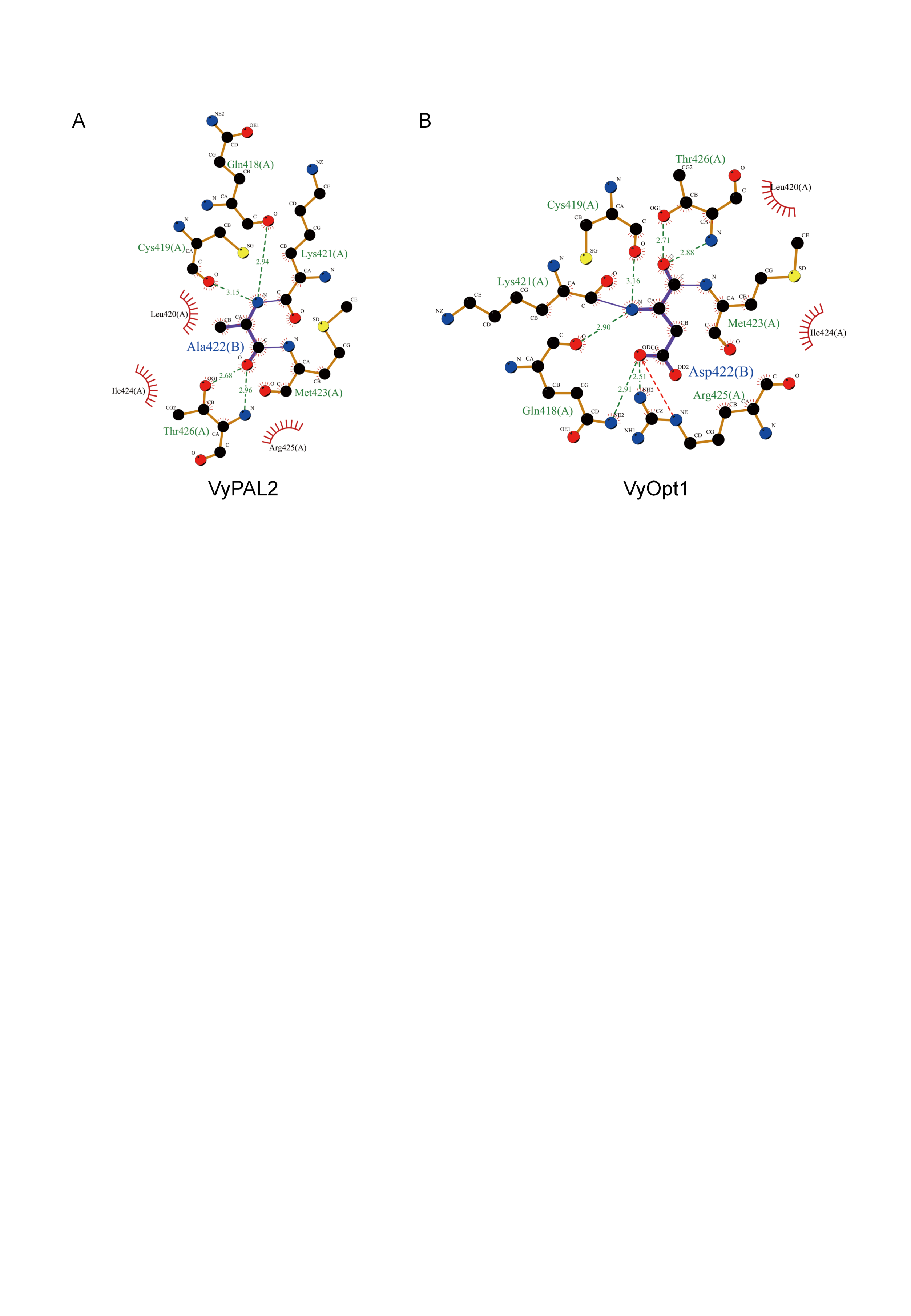

Figure S45 LigPlot diagram of the A422D mutant and its interactions with surrounding residues. (A) The interaction network of A422D with surrounding residues in wild-type VyPAL2 (B) The interaction of A422D with surrounding residues in mutant VyOpt1.

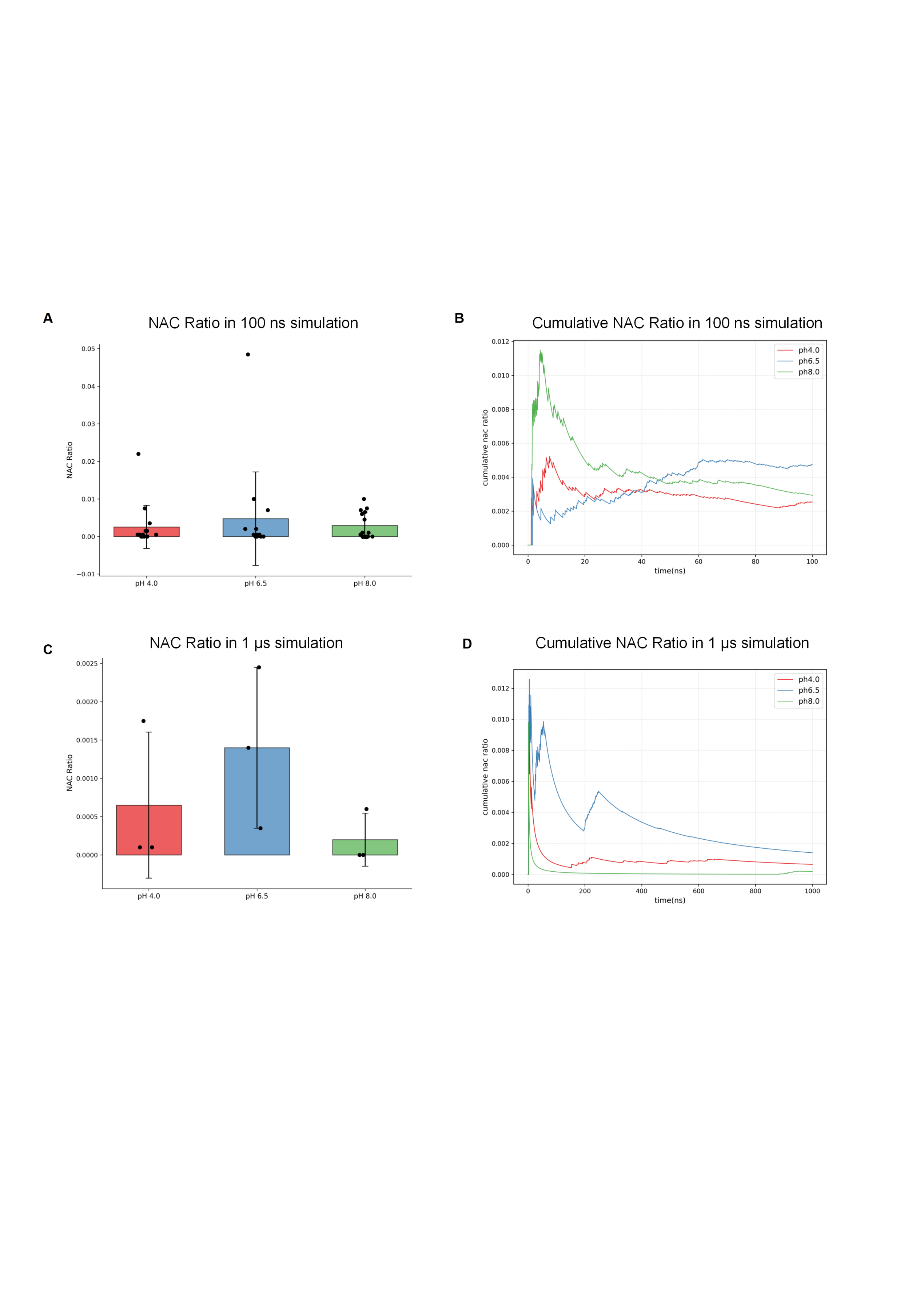

Figure S46 Proportions of Cys-attacking NAC including the hydrogen-bonding with oxyanion hole at three different pH values in 100 ns and 1 μs simulations. (A) Proportions of NACs in 100 ns simulations. (B) Time evolution of NACs during 100 ns simulations. Data are presented as mean ± SD of *n*=15 independent simulations. (C) Proportions of NACs in 1 μs simulations. (D) Time evolution of NACs during 100 ns simulations. Data are presented as mean ± SD of *n*=15 independent simulations. Data are presented as mean ± SD of *n*=3 independent simulations. Black dots represent individual data points. Statistical analysis was performed using Student’s t-test. No statistically significant differences were observed between pH conditions (all *p*>0.05). The NAC ratio and cumulative NAC ratio on all the Y-axis range from 0 to 1.

Supplementary tables

Table S1 PCR cloning primers of New Legumains.

| Primer | Nucleotide Sequence |
| --- | --- |
| VdiPAL1_F | CTGGTGCCGCGCGGCAGCCATCTCCGGTTGCCGTCAGAAG |
| VdiPAL1_R | GTGGTGGTGGTGGTGGTGCTCGAGCGCACTGAACCCTTCGG |
| VdiPAL3_F | CTGGTGCCGCGCGGCAGCCATCTCCAGTTGCCGTCAGAAGC |
| VdiPAL3_R | GTGGTGGTGGTGGTGGTGCTCGAGAGCACTGAACCCTTCGG |
| VdiAEP2_F | CTGGTGCCGCGCGGCAGCCATCTCCGCTTACCATCGGAAGC |
| VdiAEP2_R | GTGGTGGTGGTGGTGGTGCTCGAGTGCGCTGAACCCGTT |
| VdifPAL1_F | GTGCCGCGCGGCAGCCATCTCCGGTTGCCGTCAGAAG |
| VdifPAL1_R | GTGGTGGTGGTGGTGCTCGAGCGCACTGAACCCTTCGGC |
| VdivAEP2_F | GTGCCGCGCGGCAGCCATCTCCGGTTACCGTCGCAA |
| VdivAEP2_R | GTGGTGGTGGTGGTGCTCGAGAGCACTGAAACCTTTGACAAGAGA |
| VdivAEP3_F | GTGCCGCGCGGCAGCCATCTCCGCTTACCATCGGAAGC |
| VdivAEP3_R | GTGGTGGTGGTGGTGCTCGAGTGCGCTGAACCCGTTGCG |
| VdivPAL1_F | GTGCCGCGCGGCAGCCATCTCCGGTTGCCGTCAGAAG |
| VdivPAL1_R | GTGGTGGTGGTGGTGCTCGAGAGCACTGAACCCTTCGGCA |
| VpaPAL1_F | GTGCCGCGCGGCAGCCATCTCCGCTTGCCGTCAGAA |
| VpaPAL1_R | GTGGTGGTGGTGGTGCTCGAGAGCACTGAACCCTTCAGCAAGA |
| VbaPAL1_F | GTGCCGCGCGGCAGCCATCTCCGGTTGCCGTCAGAAG |
| VbaPAL1_R | GTGGTGGTGGTGGTGCTCGAGCGCACTGAACCCTTCGGC |
| VbaPAL2_F | GTGCCGCGCGGCAGCCATCTCCGCTTACCATCGGAAGC |
| VbaPAL2_R | GTGGTGGTGGTGGTGCTCGAGTGCGCTGAACCCGTTGAC |
| VbaPAL3_F | GTGCCGCGCGGCAGCCATCTCCGCTTGCCATCAGAAGC |
| VbaPAL3_R | GTGGTGGTGGTGGTGCTCGAGTGCGCTGAACCCGTTGAC |
| VinPAL1_F | GTGCCGCGCGGCAGCCATCTCCGGTTGCCGTCAGAAG |
| VinPAL1_R | GTGGTGGTGGTGGTGCTCGAGCGCACTGAACCCTTCGGC |
| VinPAL3_F | GTGCCGCGCGGCAGCCATCTCCGGTTGCCAACGGAA |
| VinPAL3_R | GTGGTGGTGGTGGTGCTCGAGACTGAACCCTTCGGCAAGAGA |
| VinPAL2_F | GTGCCGCGCGGCAGCCATCTCCGGTTGCCGTCAGAAG |
| VinPAL2_R | GTGGTGGTGGTGGTGCTCGAGTGCACTGAACCCTTCGGCA |
| VgrPAL1_F | GTGCCGCGCGGCAGCCATCTCCGATTGCCGTCAGAAGC |
| VgrPAL1_R | GTGGTGGTGGTGGTGCTCGAGCGCACTGAACCCTTCCGC |
| VgrPAL2_F | GTGCCGCGCGGCAGCCATCTCCGGTTGCCAACGGAA |
| VgrPAL2_R | GTGGTGGTGGTGGTGCTCGAGTGCACTGAAACCTTTTTTGAGAGA |
| VcoPAL1_F | GTGCCGCGCGGCAGCCATCTCCGGTTGCCGTCAGAAG |
| VcoPAL1_R | GTGGTGGTGGTGGTGCTCGAGAGCACTGAACCCTTCGGCA |
| VcoPAL2_F | GTGCCGCGCGGCAGCCATCTCCGGTTGCCGTCAGAAG |
| VcoPAL2_R | GTGGTGGTGGTGGTGCTCGAGCGCACTGAACCCTTCGGC |
| VcoAEP3_R | GTGCCGCGCGGCAGCCATCTCCGCTTACCATCGGAAGC |
| VcoAEP3_R | GTGGTGGTGGTGGTGCTCGAGTGCGCTGAACCCGTTGCC |
| VcoAEP4_F | GTGCCGCGCGGCAGCCATTCTTCAGCTGCAACTAAGTGGGC |
| VcoAEP4_R | GTGGTGGTGGTGGTGCTCGAGAGCACTGAAACCTTTGGCAAGA |
| VmoPAL1_F | GTGCCGCGCGGCAGCCATCTCCGGTTGCCGTCAGAAG |
| VmoPAL1_R | GTGGTGGTGGTGGTGCTCGAGCGCACTGAACCCTTCGGC |
| VmoPAL2_F | GTGCCGCGCGGCAGCCATCTCCGGTTGCCGTCAGAAG |
| VmoPAL2_R | GTGGTGGTGGTGGTGCTCGAGCGCACTGAACCCTTCTGCA |
| VmoAEP3_R | GTGCCGCGCGGCAGCCATCTCCGGTTACCGTCGTTAGAAT |
| VmoAEP3_R | GTGGTGGTGGTGGTGCTCGAGTGCACTGAAACCTTTGACAAGAGA |
| VmoPAL4_F | GTGCCGCGCGGCAGCCATCTCCGGTTGCCAACGGAT |
| VmoPAL4_R | GTGGTGGTGGTGGTGCTCGAGTGCACTGAAACCTTTTTTGAGAGA |
| VmoAEP7_R | GTGCCGCGCGGCAGCCATCTCCGGTTACCGTCGCAA |
| VmoAEP7_R | GTGGTGGTGGTGGTGCTCGAGAGCACTGAAACCTTCGACAAGAG |
| VmoPAL5_F | GTGCCGCGCGGCAGCCATCTCCGGTTGCCGTCAGAAG |
| VmoPAL5_R | GTGGTGGTGGTGGTGCTCGAGAGCACTGAACCCTTCGGCA |
| VmoPAL6_F | GTGCCGCGCGGCAGCCATCTCCGGTTGCCGTCAGAAG |
| VmoPAL6_R | GTGGTGGTGGTGGTGCTCGAGCGCACTGAACCCTTCGGC |
| VmoAEP8_R | GTGCCGCGCGGCAGCCATCTCCGGTTGCCAACGGAA |
| VmoAEP8_R | GTGGTGGTGGTGGTGCTCGAGTGCACTGAAACCTTTTTTGAGAGA |
| VmoAEP9_R | GTGCCGCGCGGCAGCCATCTCCGCTTACCATCGGAAGC |
| VmoAEP9_R | GTGGTGGTGGTGGTGCTCGAGTGCGCTGAACCCGTTGCC |
| VmoAEP10_F | GTGCCGCGCGGCAGCCATCTCCGCTTACCATCGGAAGC |
| VmoAEP10_R | GTGGTGGTGGTGGTGCTCGAGTGCGCTGAACCCGTTGCC |
| VmoAEP11_F | GTGCCGCGCGGCAGCCATCTCCGGTTACCGTCGTCAGA |
| VmoAEP11_R | GTGGTGGTGGTGGTGCTCGAGTGCACTGAAACCTTTGACAAGAGA |
| VmoAEP12_F | GTGCCGCGCGGCAGCCATATTCTGATGCCAACCGATAGAGTC |
| VmoAEP12_R | GTGGTGGTGGTGGTGCTCGAGGGCACTATAACCTCGGATAAACG |
| VmoAEP13_F | GTGCCGCGCGGCAGCCATATTCTGATGCCAACCGATAGAATC |
| VmoAEP13_R | GTGGTGGTGGTGGTGCTCGAGGGCACTATAACCACGGATAAACG |
| VbiPAL1_F | GTGCCGCGCGGCAGCCATCTCCGGTTGCCGTCAGAAA |
| VbiPAL1_R | GTGGTGGTGGTGGTGGTGCTCGAGCGCACTGAACCCTTCGGC |
| VodAEP1_F | TGGTGCCGCGCGGCAGCCATATGAGACGTTTTGTCGCCGG |
| VodAEP1_R | CTCAGTGGTGGTGGTGGTGGTGCTCGAGTGCGCTGAACCCGTTGC |
| VodAEP2_F | TGGTGCCGCGCGGCAGCCATATGAGACGTTTTGTCGCCGG |
| VodAEP2_R | CTCAGTGGTGGTGGTGGTGGTGCTCGAGTGCTGATGCCTCGGCCA |
| VodPAL1_F | GGTGCCGCGCGGCAGCCATATGAAACTACTCGCCGCCG |
| VodPAL1_R | TCTCAGTGGTGGTGGTGGTGGTGCTCGAGAGCACTGAACCCTTCGGC |
| VodPAL2_F | GGTGCCGCGCGGCAGCCATATGAAACTACTCGCCGCCG |
| VodPAL2_R | TCTCAGTGGTGGTGGTGGTGGTGCTCGAGAGCACTGAACCCTTCGGC |
| VodPAL3_F | GGTGCCGCGCGGCAGCCATATGAAACTACTCGCCGCCG |
| VodPAL3_R | TCTCAGTGGTGGTGGTGGTGGTGCTCGAGAGCACTGAACCCTTCGGC |
| VfaPAL1_F | GTGCCGCGCGGCAGCCATCTCCGGTTGCCGTCAGAAG |
| VfaPAL1_R | GTGGTGGTGGTGGTGCTCGAGCGCACTGAACCCTTCGGC |
| VfaAEP2_F | GTGCCGCGCGGCAGCCATCTCCGGTTGCCAACGGAA |
| VfaAEP2_R | GTGGTGGTGGTGGTGCTCGAGTGCACTGAAACCTTTCTTGAGAGA |
| VyuAEP1_F | CTGGTGCCGCGCGGCAGCCATCTCCGCTTACCATCGGAAGTTT |
| VyuAEP1_R | GTGGTGGTGGTGGTGCTCGAGTGCACTGAAACCTTTCTTGAGAGA |
| VyuAEP2_F | GTGCCGCGCGGCAGCCATCTCCGCTTACCATCGGAAGC |
| VyuAEP2_R | GTGGTGGTGGTGGTGCTCGAGAGCACTGAAACCTTTGACAAGAGA |
| VyuAEP3_F | GTGCCGCGCGGCAGCCATCTCCGGTTGCCAACGGAA |
| VyuAEP3_R | GTGGTGGTGGTGGTGCTCGAGTGCACTGAAACCTTTCTTGAGAGA |
| VyuAEP4_F | GTGCCGCGCGGCAGCCATCTCCGGTTACCGTCGCAA |
| VyuAEP4_R | GTGGTGGTGGTGGTGCTCGAGAGCACTGAAACCTTTGACAAGAGA |
| VyuAEP5_F | GTGCCGCGCGGCAGCCATCTCCGGTTACCGTCGTCAGA |
| VyuAEP5_R | GTGGTGGTGGTGGTGCTCGAGTGCACTGAAACCTTTGACAAGAGA |
| VyuAEP6_F | GTGCCGCGCGGCAGCCATCTCCGGTTGCCGTCAGAAG |
| VyuAEP6_R | GTGGTGGTGGTGGTGCTCGAGCGCACTGAACCCTTCGGC |
| VdaPAL1_F | GTGCCGCGCGGCAGCCATCTCCGCTTGCCGTCAGAA |
| VdaPAL1_R | GTGGTGGTGGTGGTGCTCGAGAGCACTGAACCCTTCGGCA |
| VdaAEP1_F | GTGCCGCGCGGCAGCCATCTCCGGTTGCCAACGGAA |
| VdaAEP1_R | GTGGTGGTGGTGGTGCTCGAGTGCACTGAAACCTTTTTTGAGGG |
| VduPAL1_F | GTGCCGCGCGGCAGCCATCTCCGGTTGCCGTCAGAAG |
| VduPAL1_R | GTGGTGGTGGTGGTGCTCGAGCGCACTGAACCCTTCGGC |
| VduAEP1_F | GTGCCGCGCGGCAGCCATCTCCGGTTGCCAACGGAA |
| VduAEP1_R | GTGGTGGTGGTGGTGCTCGAGTGCACTGAAACCTTTTTTGAGAGA |
| VduAEP2_F | GTGCCGCGCGGCAGCCATCTCCGGTTACCGTCGCAA |
| VduAEP2_R | GTGGTGGTGGTGGTGCTCGAGTGCACTGAAACCTTTGACAAGAGA |
| VpiAEP1_F | GTGCCGCGCGGCAGCCATCTCCGGTTACCGTCGTTCG |
| VpiAEP1_R | GTGGTGGTGGTGGTGCTCGAGTGCACTGAAACCTTTGACAAGAGA |
| VpiAEP2_F | GTGCCGCGCGGCAGCCATCTCCGGTTGCCAACGGAA |
| VpiAEP2_R | GTGGTGGTGGTGGTGCTCGAGTGCACTGAAACCTTTCTTGAGAGA |
| VphAEP1_F | GTGCCGCGCGGCAGCCATATTCTGATGCCAACCGATAAAGTC |
| VphAEP1_R | GTGGTGGTGGTGGTGCTCGAGGGCACTATAACCCCGGATAAACG |
| VphPAL1a_F | GTGCCGCGCGGCAGCCATCTCCGGTTGCCGTCAGAAG |
| VphPAL1a_R | GTGGTGGTGGTGGTGCTCGAGCGCACTGAACCCTTCGGC |
| VphPAL1b_F | GTGCCGCGCGGCAGCCATCTCCGGTTGCCGTCAGAAG |
| VphPAL1b_R | GTGGTGGTGGTGGTGCTCGAGAGCACTGAACCCTTCGGCA |

Table S2 Novel Legumain Sequences Identified in *Viola* Species.

|  | Plant species | Legumain | LAD2 | LAD1 | Label |
| --- | --- | --- | --- | --- | --- |
| 1 | *Viola dissecta* | VdiPAL1 | AP | WVT | PAL |
| 2 |  | VdiPAL3 | AP | WIT | PAL |
| 3 |  | VdiAEP2 | GP | WGT | AEP |
| 4 | *Viola diffusa* | VdifPAL1 | AP | WVT | PAL |
| 5 | *Viola dissecta* var. *incisa* | VdivAEP2 | YA | WVV | AEP |
| 6 |  | VdivAEP3 | GP | WGT | AEP |
| 7 |  | VdivPAL1 | AP | LIT | PAL |
| 8 | *Viola patrinii* | VpaPAL1 | AP | WIT | PAL |
| 9 | *Viola banksii* | VbaPAL1 | AP | WIT | PAL |
| 10 |  | VbaPAL2 | AP | WIT | PAL |
| 11 |  | VbaPAL3 | AP | WIT | PAL |
| 12 | *Viola inconspicua* | VinPAL1 | AP | WIT | PAL |
| 13 |  | VinPAL3 | AP | WIT | PAL |
| 14 |  | VinPAL2 | AP | WIT | PAL |
| 15 | *Viola grypoceras* | VgrPAL1 | AP | LIA | PAL |
| 16 |  | VgrPAL2 | AP | WIT | PAL |
| 17 | *Viola collina* | VcoPAL1 | AP | WIT | PAL |
| 18 |  | VcoPAL2 | AP | WIT | PAL |
| 19 |  | VcoAEP3 | GA | WGL | AEP |
| 20 |  | VcoAEP4 | YP | WTV | AEP |
| 21 | *Viola moupinensis* | VmoPAL1 | AP | LIT | PAL |
| 22 |  | VmoPAL2 | AP | WIT | PAL |
| 23 |  | VmoAEP3 | GP | WGT | AEP |
| 24 |  | VmoPAL4 | AP | WIT | PAL |
| 25 |  | VmoAEP7 | YP | WAV | AEP |
| 26 |  | VmoPAL5 | AP | LIT | PAL |
| 27 |  | VmoPAL6 | AP | WVT | PAL |
| 28 |  | VmoAEP8 | GP | WGT | AEP |
| 29 |  | VmoAEP9 | GA | WGT | AEP |
| 30 |  | VmoAEP10 | YP | WVV | AEP |
| 31 |  | VmoAEP11 | YP | WVT | AEP |
| 32 |  | VmoAEP12 | GP | WGT | AEP |
| 33 |  | VmoAEP13 | GP | WGT | AEP |
| 34 | *Viola biflora* | VbiPAL1 | AP | WIT | PAL |
| 35 | *Viola odorata L.* | VodAEP1 | GA | WGV | AEP |
| 36 |  | VodAEP2 | GA | WGT | AEP |
| 37 |  | VodPAL1 | AP | LIT | PAL |
| 38 |  | VodPAL2 | AP | LIT | PAL |
| 39 |  | VodPAL3 | AP | LIT | PAL |
| 40 | *Viola fargesii* | VfaPAL1 | AP | WIT | PAL |
| 41 |  | VfaAEP2 | GP | WGT | AEP |
| 42 | *Viola yunnanensis* | VyuAEP1 | GP | WGT | AEP |
| 43 |  | VyuAEP2 | GP | WGT | AEP |
| 44 |  | VyuAEP3 | GP | WGT | AEP |
| 45 |  | VyuAEP4 | YP | WVV | AEP |
| 46 |  | VyuAEP5 | YP | WVT | AEP |
| 47 |  | VyuAEP6 | YP | WVT | AEP |
| 48 | *Viola davidii* | VdaPAL1 | AP | LIT | PAL |
| 49 |  | VdaAEP1 | GP | WGT | AEP |
| 50 | *Viola duclouxii* | VduPAL1 | AP | WIT | PAL |
| 51 |  | VduAEP1 | GP | WGT | AEP |
| 52 |  | VduAEP2 | YP | WVV | AEP |
| 53 | *Viola pilosa* | VpiAEP1 | YP | WVT | AEP |
| 54 |  | VpiAEP2 | GP | WGT | AEP |
| 55 | *Viola philippica* | VphAEP1 | GP | WGT | AEP |
| 56 |  | VphPAL1a | AP | LIA | PAL |
| 57 |  | VphPAL1b | AP | LIA | PAL |

Table S3 Data collection and refinement statistics.

|  | VdiPAL1 |
| --- | --- |
| PDB Entry | 9VSX |
| Space group | P 21 21 2 |
| a, b, c (Å) | 115.31 128.90 70.96 |
| α, β, γ (°) | 90.00 90.00 90.00 |
| Resolution (Å)^a^ | 85.95-1.86(1.96-1.87) |
| Unique reflections | 89403(12891) |
| Redundancy | 12.6(10.1) |
| Completeness (%)^a^ | 100(100) |
| I/σI | 11.4(0.9) |
| Rmerge | 0.154(3.975) |
| CC1/2^b^ | 0.997(0.444) |
| Refinement |  |
| Resolution (Å)^a^ | 35.48-1.87 |
| Rwork/Rfree ^c^ | 0.1751/ 0.2120 |
| No.atoms | 7292 |
| Protein | 6683 |
| Ligand/ion | 18 |
| Water | 591 |
| B-factor (Å) | 38.63 |
| Protein | 38.09 |
| Ligand/ion | 58.83 |
| Water | 44.18 |
| Bond length (Å)^d^ | 0.009 |
| Bond angle (°)^d^ | 1.011 |
| Ramachandran favored (%) | 97.39% |

^a^ The values in parentheses for resolution range, completeness, Rmerge and I/σ (I) correspond to the highest resolution shell. ^b^ Rmerge(I) = ΣhklΣj | I(hkl)j - <I(hkl)> | / Σhkl Σj I(hkl)j, where I(hkl)j is the jth measurement ^c^ R = Σhkl | |Fobs| - |Fcalc| |/Σhkl |Fobs|, of the intensity of reflection hkl and <I(hkl)> is the average intensity, where Rfree is calculated without a sigma cut off for a randomly chosen 5 % of reflections, which were not used for structure refinement, and Rwork is calculated for the remaining reflections. ^d^ Deviations from ideal bond lengths/angles.

Table S4 Computational strategy to guide the combination of mutations.

| Plasmid | Δddg(REU) | No. Mutation | Non-native* | Ratio | Mutation |
| --- | --- | --- | --- | --- | --- |
| C1 | -4.706 | 5 | 2 | 0.4 | M357K_V409P_N397D_N356Q_A422D |
| C23 | -4.705 | 5 | 2 | 0.4 | M357K_V409P_N397D_N356Q_A422E |
| C14 | -4.214 | 4 | 2 | 0.5 | M357K_N397D_N356Q_V409P |
| C2 | -3.616 | 4 | 2 | 0.5 | M357K_N397D_N356Q_A422D |
| C3 | -3.171 | 5 | 2 | 0.4 | V409P_I351L_N397D_N356R_A422D |
| C21 | -3.121 | 4 | 2 | 0.5 | M357K_N397D_V409A_A422D |
| C19 | -2.857 | 2 | 1 | 0.5 | M357K_N356Q |
| C9 | -2.844 | 4 | 2 | 0.5 | N397D_N356Q_V409P_A422D |
| C4 | -2.649 | 4 | 2 | 0.5 | M357K_N397D_N356Q_A422E |
| C5 | -2.421 | 4 | 2 | 0.5 | M357K_N356Q_V409A_A422D |
| C22 | -2.401 | 5 | 2 | 0.4 | M357K_V409P_I351L_N356Q_A422E |
| C6 | -2.326 | 5 | 2 | 0.4 | M357K_V409P_I351L_N356Q_A422D |
| C7 | -2.258 | 5 | 2 | 0.4 | M357K_V409P_I351L_N397D_A422D |
| C20 | -2.234 | 2 | 1 | 0.5 | M357K_N397D |
| C8 | -2.188 | 4 | 2 | 0.5 | I351L_N356Q_V409P_A422D |
| C10 | -2.154 | 3 | 2 | 0.67 | M357K_N397D_N356Q |
| C18 | -2.059 | 5 | 2 | 0.4 | M357K_I351L_N397E_N356Q_A422D |
| C16 | -1.948 | 4 | 2 | 0.5 | I351L_M357K_N356Q_V409P |
| C11 | -1.939 | 4 | 2 | 0.5 | I351L_M357K_N356Q_A422D |
| C12 | -1.866 | 3 | 2 | 0.67 | I351L_N356Q_A422D |
| C13 | -1.862 | 5 | 3 | 0.6 | M357K_V409A_N397D_N356Q_A422D |
| C17 | -1.739 | 2 | 1 | 0.5 | I351L_N356K |
| C24 | -1.687 | 2 | 1 | 0.5 | N397D_A422D |
| C25 | -1.583 | 4 | 2 | 0.5 | N397D_N356Q_V409P_A422E |
| C15 | -1.536 | 5 | 2 | 0.4 | M357K_V409P_I351L_N397D_A422E |

* Number of mutation sites with predicted amino acid composition different from the native sequence of VdiPAL1.
